# Does Human Transthyretin Aggregate in Blood Plasma or in Cardiac Tissue? A Mathematical Modeling Study

**DOI:** 10.1101/2025.07.29.667417

**Authors:** Andrey V. Kuznetsov

## Abstract

Transthyretin (TTR) amyloidosis is a progressive disease characterized by the destabilization of TTR tetramers, leading to monomer dissociation, oligomer formation, and fibril deposition in cardiac tissue. This study presents a mathematical model that captures the kinetics of TTR aggregation and deposition, integrating clinically relevant parameters to distinguish between scenarios of TTR aggregation in blood plasma and cardiac tissue. The governing equations are derived by applying conservation principles to the number of TTR tetramers, monomers, and oligomers within defined control volumes, such as the blood plasma or heart. The model predicts that the scenario with fibril accumulation within cardiac tissue better reflects the clinical progression of wild-type TTR amyloidosis, aligning with observed patient survival times and measured tetramer concentrations. In this scenario, the model successfully recapitulates published physiological data, accurately predicting a total plasma TTR concentration of approximately 25 mg/dL. Sensitivity analyses reveal that the rate of tetramer dissociation and the half-deposition time of TTR oligomers are key regulators of disease progression. Lower tetramer dissociation rates, mimicking the stabilizing effect of tafamidis, reduce fibril accumulation and biological aging. Conversely, faster oligomer deposition into fibrils leads to reduced oligomer concentrations but accelerated tissue fibril accumulation. The model predicts that, after five years of disease progression, the volumetric ratio of deposited TTR fibrils to baseline myocardial volume may reach 58% in the tissue aggregation scenario, compared to 10% when aggregation occurs in plasma. These predictions are consistent with clinical observations that link higher amyloid burden to increased mortality. Overall, the findings support a pathogenic mechanism in which TTR aggregation predominantly occurs in cardiac tissue, highlighting potential therapeutic targets to modulate disease progression. A correlation is proposed for estimating biological age in humans, which incorporates both calendar age and the volumetric ratio of deposited TTR fibrils to baseline myocardial volume.

## 1. Introduction

Transthyretin (TTR) is a transport protein primarily produced in the liver that carries the hormone thyroxine and vitamin A (retinol) in the bloodstream, both of which are otherwise insoluble under normal conditions (Kittleson et al., 2020; Gao et al., 2022). However, TTR is also an amyloidogenic protein capable of forming insoluble deposits in various tissues—most notably in the heart—leading to the formation of amyloid fibrils. The pathological accumulation of transthyretin amyloid within the cardiac muscle increases myocardial stiffness, which progressively impairs the heart’s ability to pump effectively and ultimately results in congestive heart failure (Ruberg et al., 2019; Smiley et al., 2022). These deposits contribute to progressive diastolic and systolic dysfunction, ultimately resulting in congestive heart failure and, if left untreated, death. This condition is known as cardiac amyloidosis (Ruberg and Berk, 2012).

(Criddle et al., 2024) proposed two competing hypotheses for how insoluble amyloid fibrils composed of TTR appear in tissue. In the first, TTR monomers misfold and aggregate into protofibrils in the bloodstream, which then circulate and deposit in specific tissues; in the second, TTR monomers circulate until they bind directly to tissue sites, where they form amyloid deposits *in situ*.

One of the objectives of this paper is to investigate how the accumulation of TTR fibrils in the heart relates to biological age. Aging is characterized by the gradual accumulation of harmful changes at molecular, cellular, tissue, and organ levels, collectively referred to as “damage.” While the body has repair mechanisms that counteract some of these changes, their effectiveness declines over time. It is important to note, however, that not all age-related alterations are detrimental—some may be adaptive, and others may have a neutral impact on health and function (Gladyshev et al., 2021).

Because aging involves a complex interplay of many interrelated biological processes, calendar age does not always accurately reflect an individual’s physiological condition. To capture this complexity, researchers have developed various computational tools known as aging clocks, which estimate biological age based on biomarkers that reflect different aspects of the aging process (Min et al., 2024). The diversity of these clocks emphasizes the multifaceted nature of aging, as no single biomarker can capture it fully. Existing models target different biological mechanisms, including epigenetic clocks that track DNA methylation changes, telomere-based clocks that monitor telomere shortening, and epitranscriptomic clocks that measure alterations in RNA methylation (Wang et al., 2023; Han, 2024; Crimmins et al., 2024). Each provides a distinct perspective on the biological underpinnings of aging.

Recently, (Dormann and Lemke, 2024) introduced the concept of an aging clock centered on the aggregation of intrinsically disordered proteins (IDPs)—such as amyloid beta (Aβ), tau, α-synuclein, and TDP-43. Unlike structured proteins, IDPs lack a stable folded conformation, making them prone to aggregate, as their aggregated forms are thermodynamically more favorable than their native disordered states (Vetri and Foderà, 2015). The appeal of an IDP-based aging clock stems from the fact that protein aggregation is intimately linked to aging and serves as a hallmark of neurodegenerative diseases like Alzheimer’s and Parkinson’s, for which advanced age remains the greatest risk factor. Building on this idea, (Kuznetsov, 2025a) proposed tracking the cumulative neurotoxicity of Aβ oligomers as a potential biomarker for the biological age of neural cells.

The accumulation of misfolded, aggregation-prone proteins is a systemic phenomenon, not confined to the brain. For example, although TTR is not an inherently disordered protein, it can aggregate when its native structure becomes destabilized. This paper introduces a new biomarker defined as the volumetric ratio of deposited TTR fibrils to baseline myocardial volume, 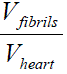. This biomarker captures the time-dependent progression of TTR fibril deposition in cardiac tissue and its effect on biological age.

## 2. Materials and models

### 2.1. Governing equations

#### 2.1.1. Conservation of TTR tetramers in the blood plasma

TTR tetramers are stable and are not prone to aggregation. It is well established that the dissociation of TTR tetramers into monomers is the initial step in TTR fibril formation (Almeida et al., 2024). Furthermore, due to the high stability of TTR tetramers (Sanguinetti et al., 2022), their dissociation into monomers represents the rate-limiting step in amyloid fibril formation (Esperante et al., 2021; Rinauro et al., 2024). Tetramer dissociation appears to be irreversible, as reported in (Quintas et al., 1999).

The mathematical framework is based on mass balance equations that govern the dynamics of tetramers, monomers, free oligomers, and oligomers incorporated into TTR amyloid fibrils. The model uses the lumped capacitance approach, assuming a uniform spatial distribution of all TTR species in both blood plasma and cardiac tissue, with time (*t*) as the only independent variable. In this framework, time is interchangeable with calendar age. Table 1 defines the dependent variables, while Table 2 provides a summary of the model parameters.

**Table 1.** Summary of dependent variables in the model.

| Symbol | Definition | Unit |
| --- | --- | --- |
| $[B]_{plasma}$ | Molar concentration of free TTR oligomers in the blood plasma | $\mu\text{M}$ |
| $[B]_{tissue}$ | Molar concentration of free TTR oligomers in the cardiac tissue | $\mu\text{M}$ |
| $[I]_{plasma}$ | Molar concentration of TTR oligomer units sequestered in amyloid fibrils circulating in the blood plasma. This variable represents the total molar concentration of TTR oligomers that have transitioned from a soluble state and been incorporated into stable amyloid filaments | $\mu\text{M}$ |
| $[I]_{tissue}$ | Molar concentration of TTR oligomer units sequestered in amyloid fibrils that infiltrated the cardiac tissue. This variable represents the total molar concentration of TTR oligomers that have transitioned from a soluble state and been incorporated into stable amyloid filaments. It is a measure of the cumulative amyloid burden expressed in terms of its constituent oligomeric units | $\mu\text{M}$ |
| $[M]_{plasma}$ | Molar concentration of TTR monomers in the blood plasma | $\mu\text{M}$ |
| $[M]_{tissue}$ | Molar concentration of TTR monomers in the cardiac tissue | $\mu\text{M}$ |
| $[T]$ | Molar concentration of TTR tetramers, representing the native form of TTR, in the blood plasma | $\mu\text{M}$ |
| $[TTR]_{total}$ | Total molar concentration of all TTR species in the blood plasma, expressed in monomeric equivalents (where one mole of tetramers corresponds to four moles of monomers) | $\mu\text{M}$ |

**Table 2.**
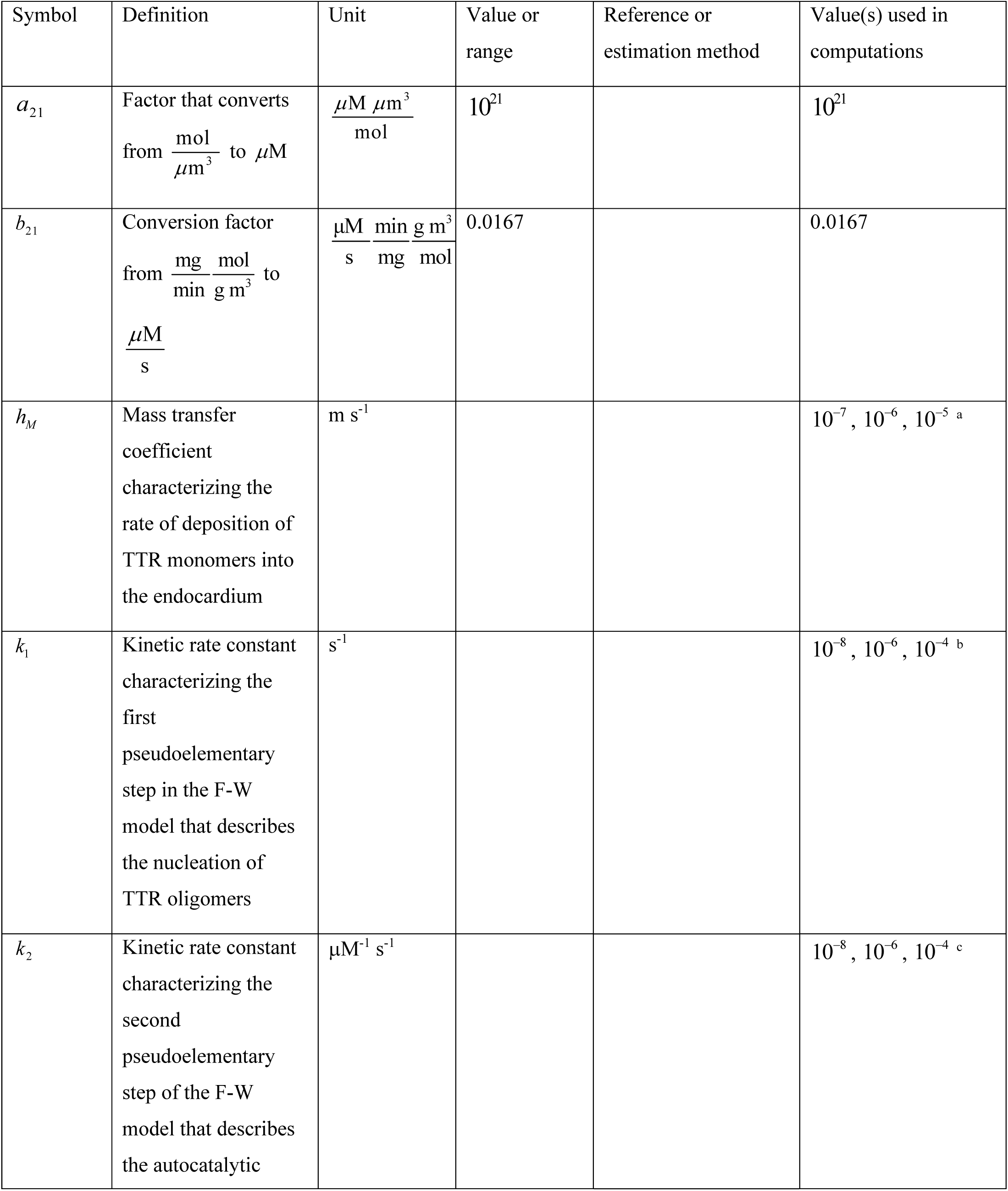
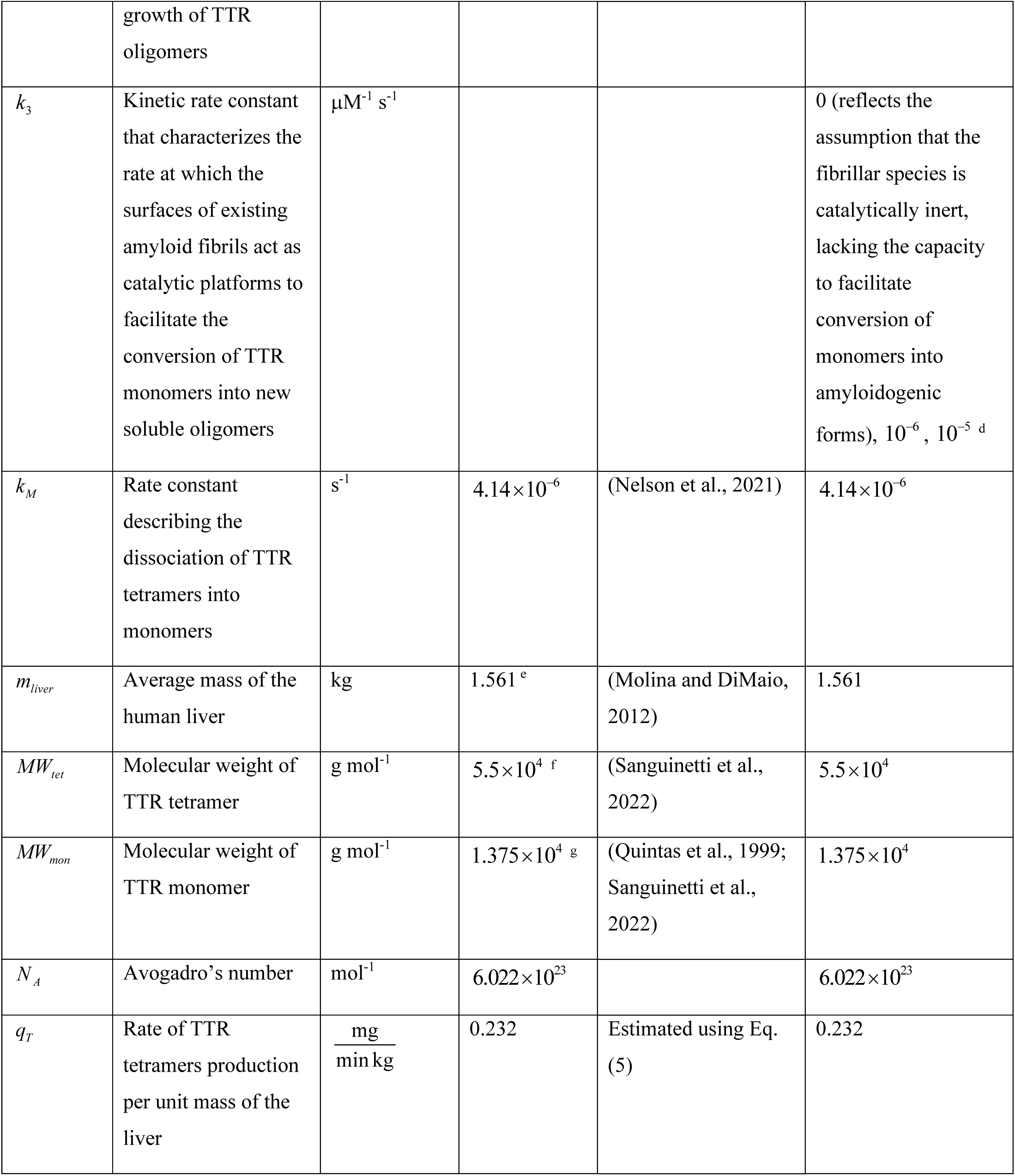
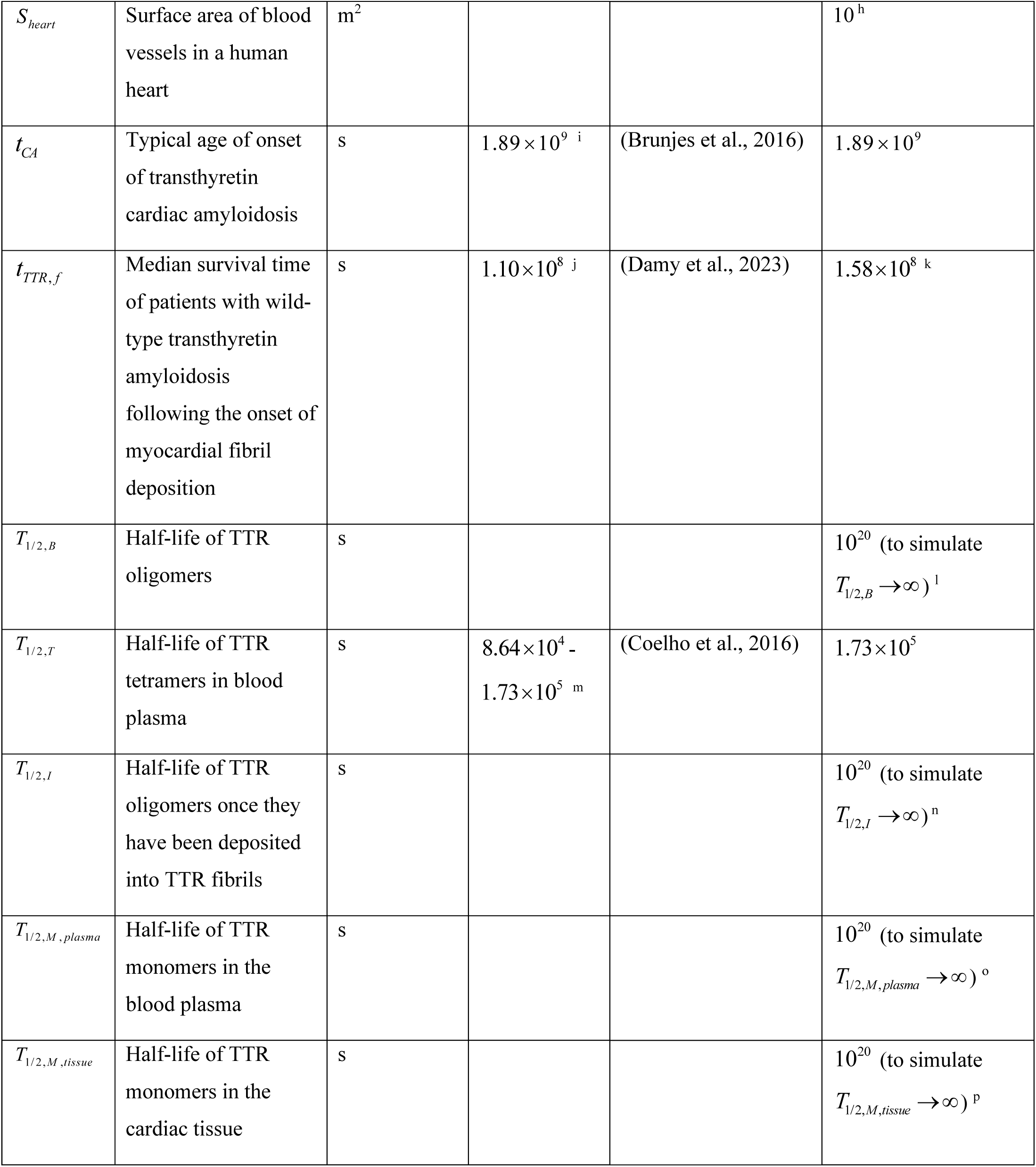
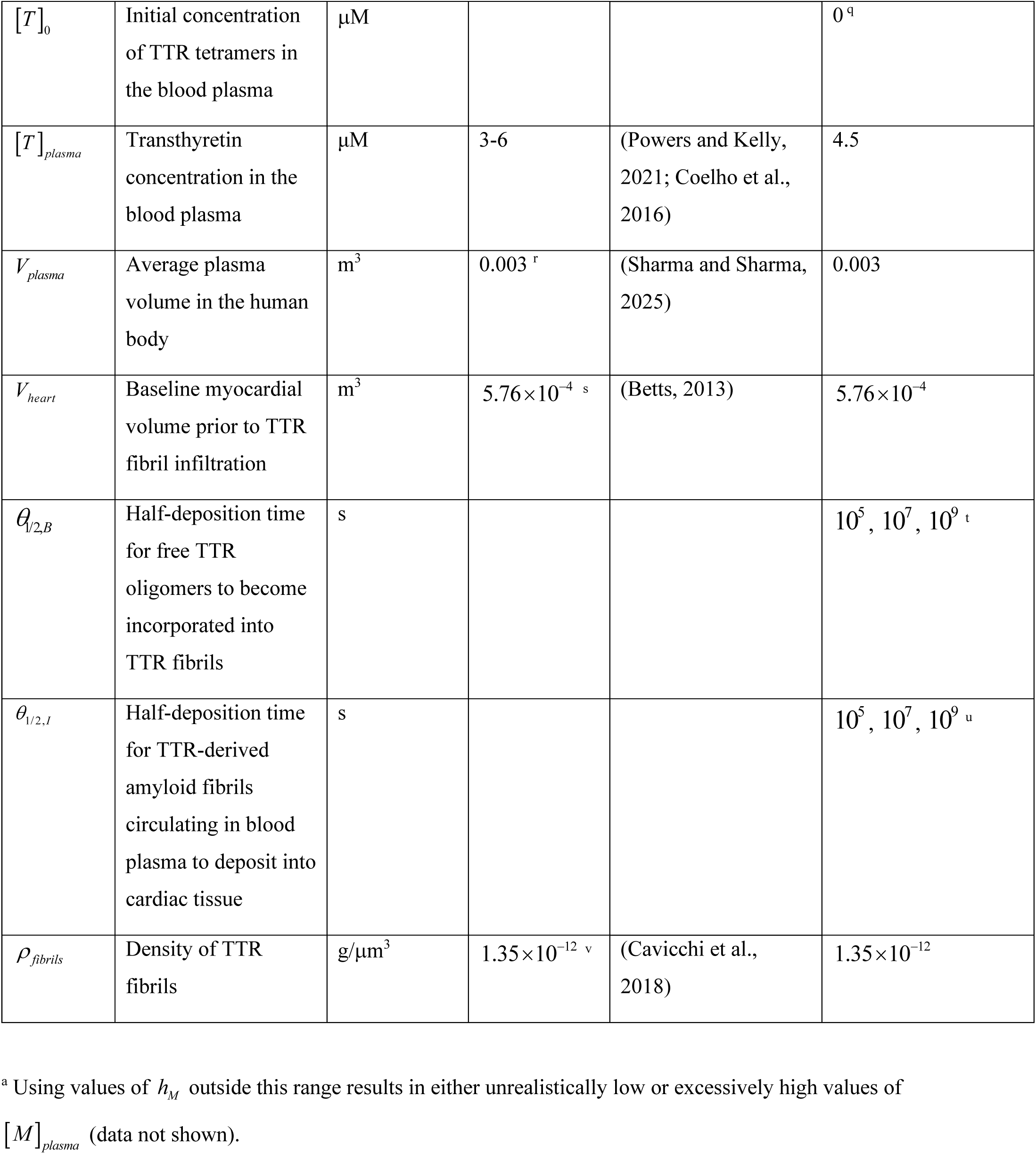
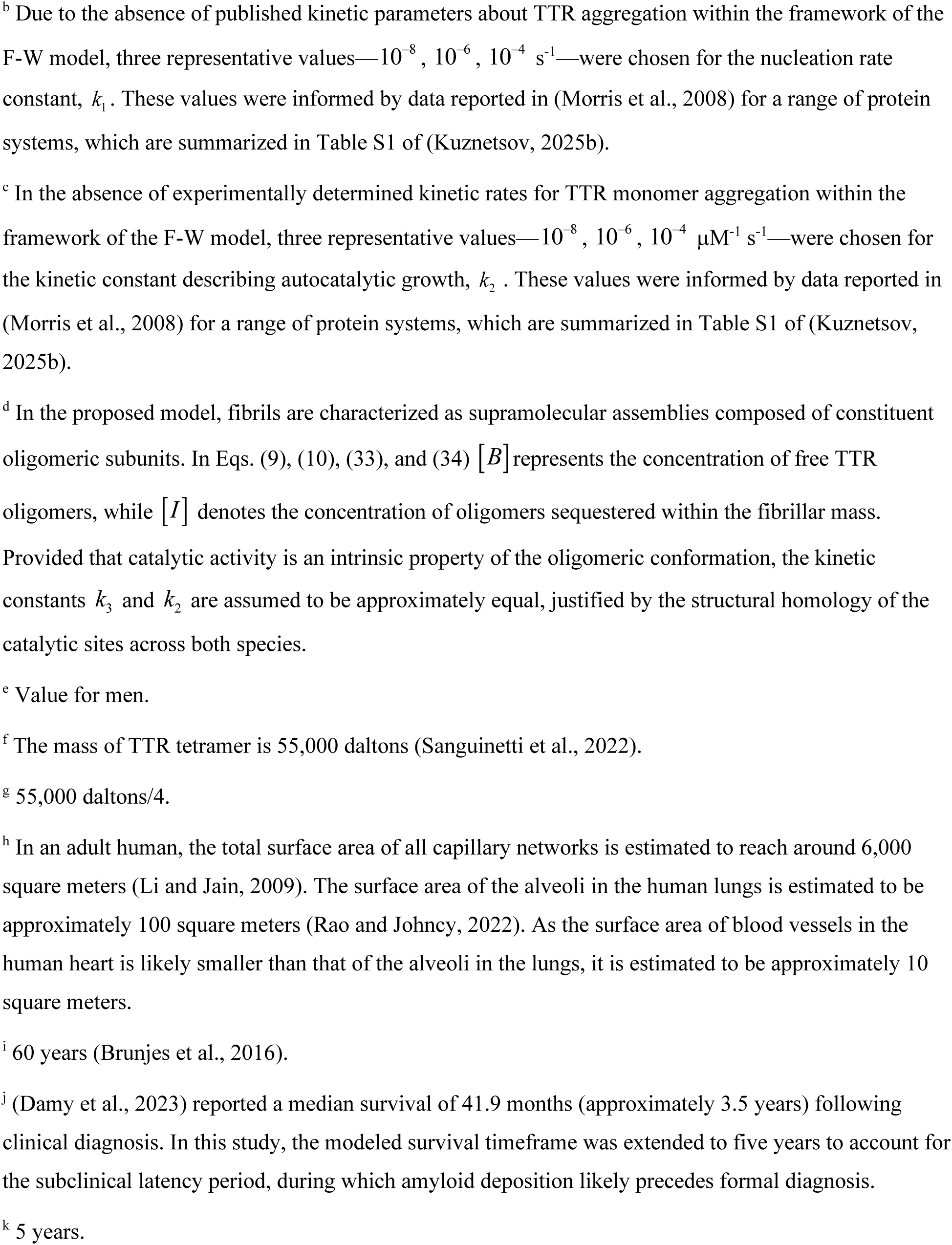
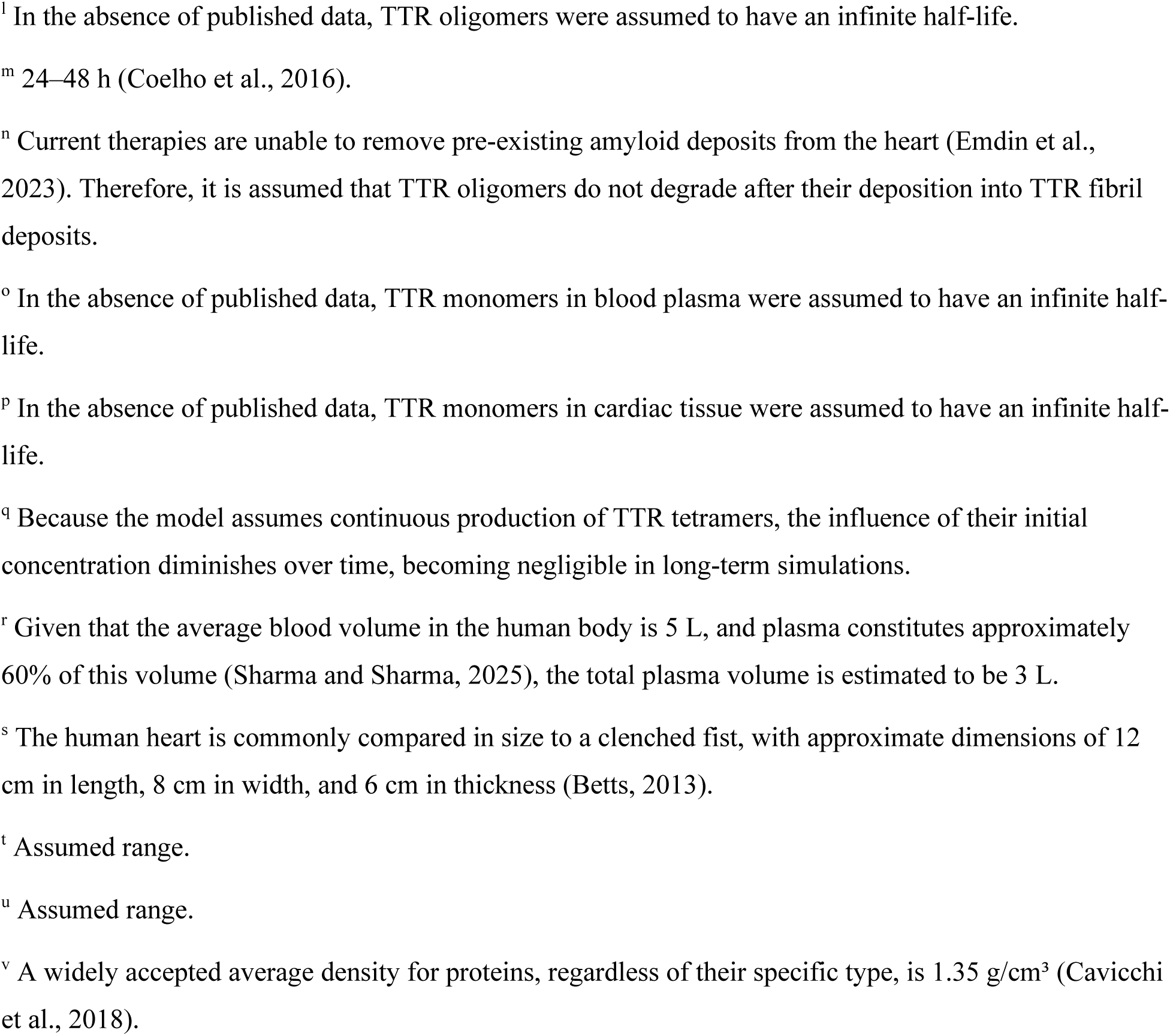
Summary of model parameters along with their estimated values.

The dissociation of TTR occurs in blood plasma and is described by the following equation:

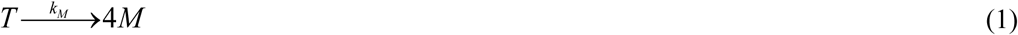

Expressing the conservation of TTR tetramers in blood plasma and dividing both sides of the equation by *V_plasma_* yields the following equation:

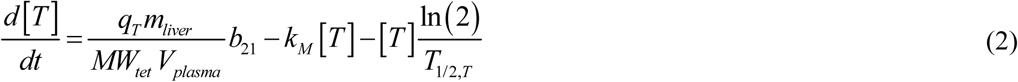

where *q_T_* denotes the rate of TTR tetramer production per unit mass of liver tissue. In Eq. (2), the first term on the right-hand side represents tetramer production in the liver; the second term accounts for the reduction in tetramer concentration due to their conversion into monomers; and the third term reflects the decline in tetramer concentration resulting from their limited half-life.

Eq. (2) must be solved with the following initial condition:

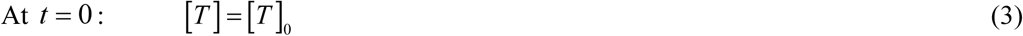

As noted in footnote “q” of Table 2, the effect of the initial condition becomes negligible when the duration of TTR aggregation is sufficiently long, allowing [*T*]_0_ to be set to zero.

Under steady-state conditions, the solution of Eq. (2) is:

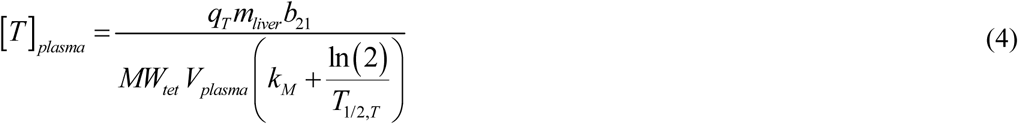

Eq. (4) can be used to estimate the value of *q_T_* if [*T*]*_plasma_* is known:

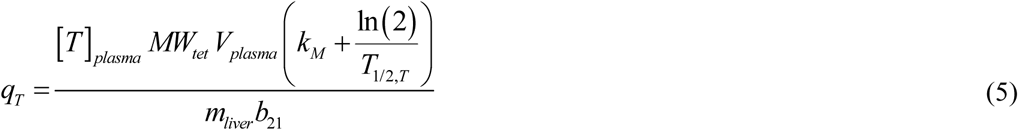

Subsequent equations within the model are contingent upon the assumed site of TTR monomer aggregation. Two scenarios are considered: (1) TTR monomers aggregate in the blood plasma, and the resulting protofibrils deposit in cardiac tissue, where they are uniformly distributed by diffusion (Fig. 1a); (2) TTR monomers first deposit in cardiac tissue, where they subsequently aggregate (Fig. 1b). These scenarios are consistent with those explored in (Criddle et al., 2024), who employed a different model that did not include autocatalysis. Similarly, (Criddle et al., 2020) did not incorporate autocatalysis in their simulations either.

**Fig. 1.**
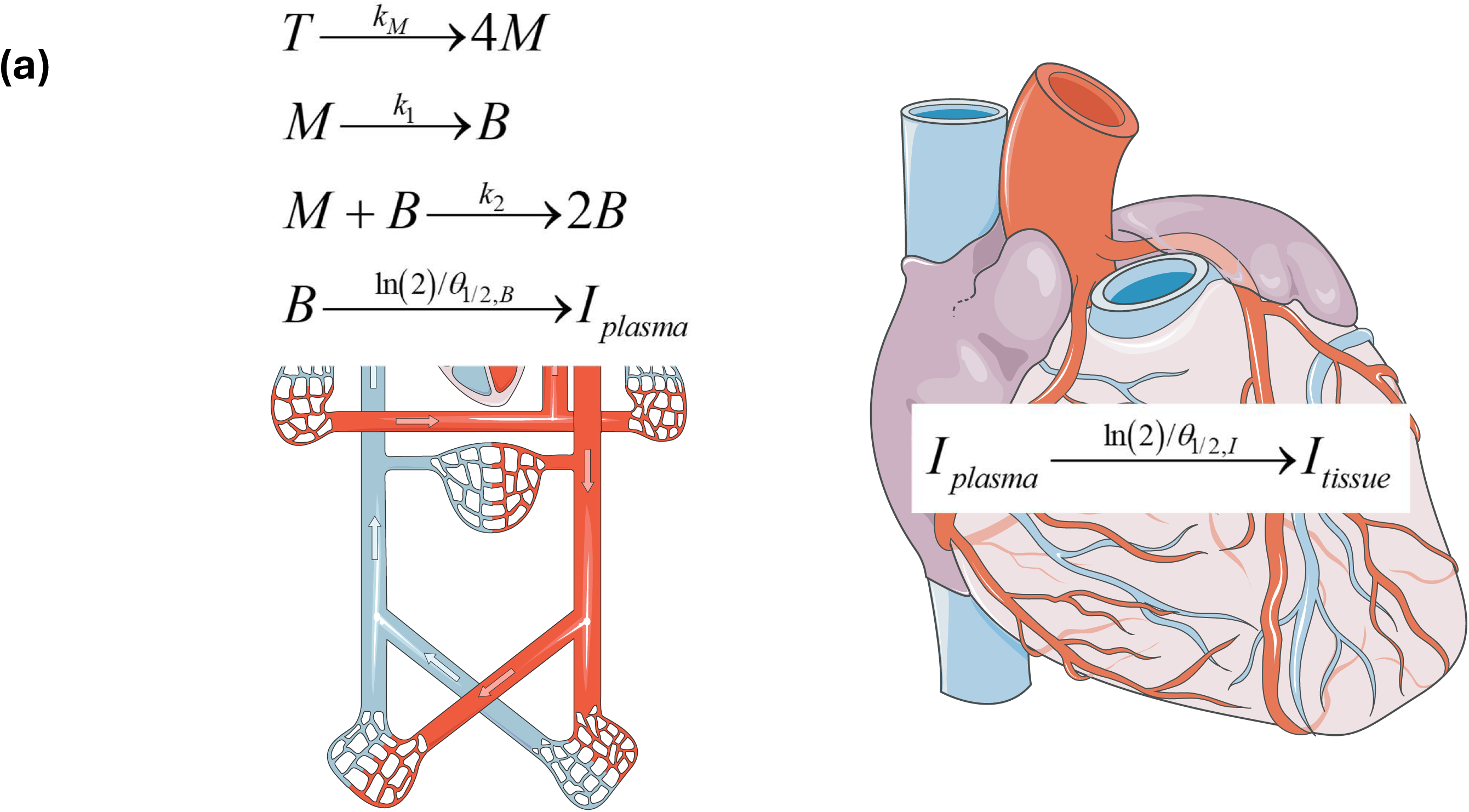
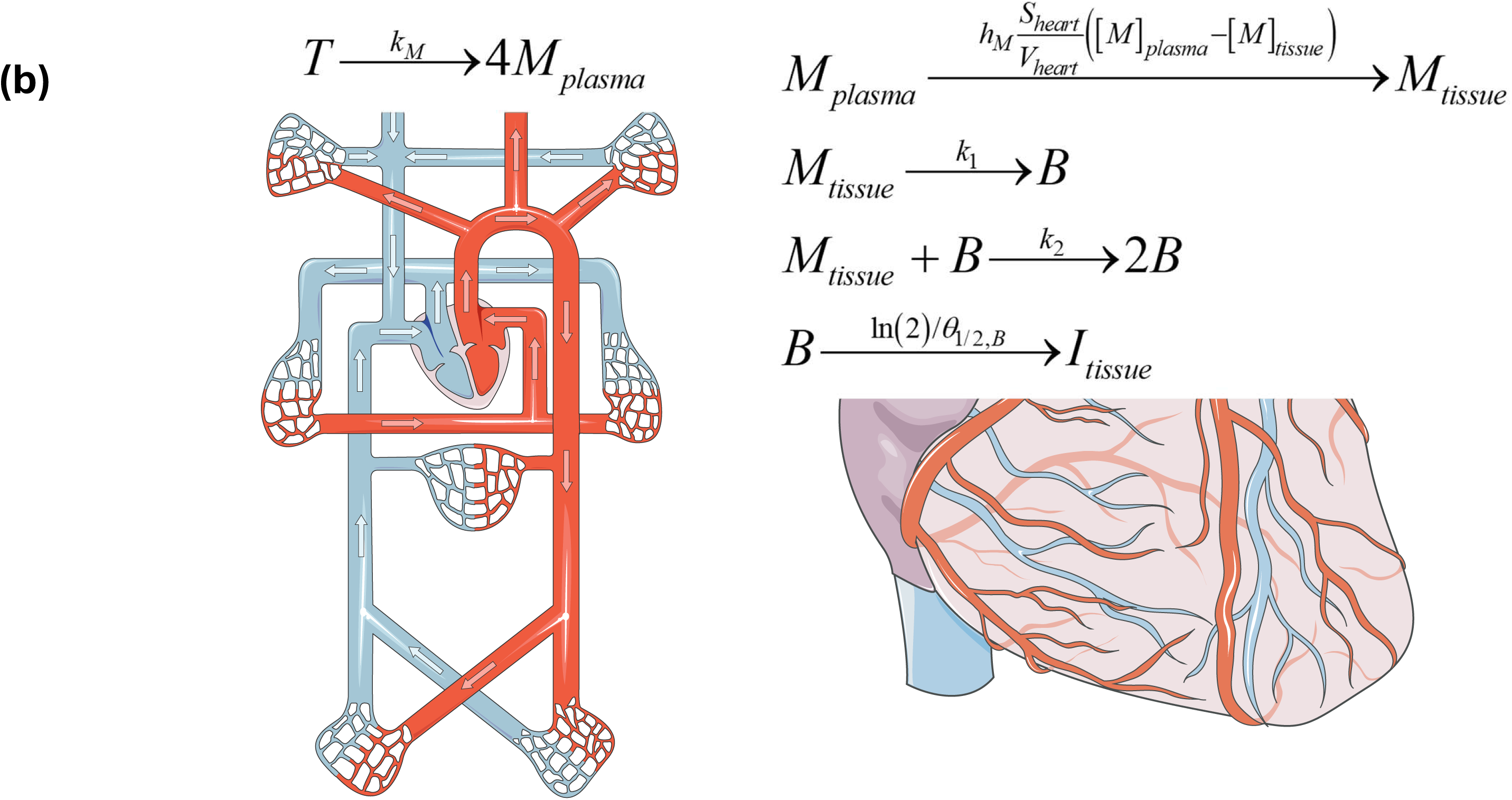
Schematic diagram illustrating two possible scenarios by which TTR fibril deposits may form in the heart: (a) TTR monomers misfold and assemble into protofibrillar formations within the bloodstream before depositing in the heart. (b) TTR monomers remain in circulation until they directly attach to sites within the heart, where they form amyloid deposits *in situ*. *Figure generated with the aid of servier medical art, licensed under a creative commons attribution 3.0 generic license.* http://Smart.servier.com.

#### 2.1.2. Conservation equations for the scenario when amyloid fibrils form in blood plasma

In many protein misfolding diseases, the aggregation of misfolded proteins is described by two key mechanisms: primary and secondary nucleation. Secondary nucleation is an autocatalytic process in which new aggregates form when monomers bind to and nucleate on the surface of existing fibrils (Rinauro et al., 2024).

After TTR tetramers dissociate into monomers—which are prone to aggregation—the monomers assemble into oligomers and subsequently form amyloid fibrils (Gao et al., 2022; Criddle et al., 2020). Aggregation of TTR monomers into oligomers is simulated using the Finke-Watzky (F-W) model. The F-W model is a minimal two-step model that reduces a complex multistep aggregation process to two pseudoelementary steps: parallel nucleation and autocatalytic growth (Morris et al., 2008; Iashchishyn et al., 2017; Finke et al., 2020). It has been effectively used to model the aggregation of various neurological proteins (Morris et al., 2008). When the two-step model is applied to simulate the formation of amyloid aggregates (Rinauro et al., 2024), the first pseudoelementary step describes the nucleation process of TTR oligomers:

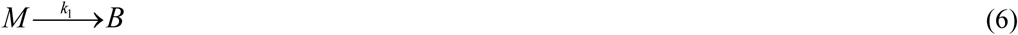

It is worth noting that, as a minimal model, the F-W model does not differentiate between the size (and mass) of a monomer and an oligomer. An oligomer is simply regarded as an activated monomer that is capable of autocatalysis. Once TTR oligomers (phase *B*) are formed, their surfaces are assumed to catalyze the formation of new oligomers from monomers (Aragonès Pedrola et al., 2025) in a prion-like manner—an autocatalytic process similar to that observed in several other amyloid disorders (Criddle et al., 2020). The potential involvement of a seed-dependent mechanism in TTR cardiac amyloidosis has been reviewed in (Morfino et al., 2022). Accordingly, and in line with the F-W model, the second pseudo-elementary step represents the autocatalytic conversion of monomers into TTR oligomers and is described as follows:

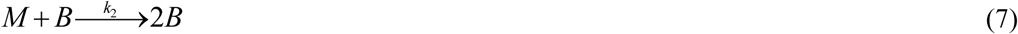

Recent studies suggest that amyloid fibrils may catalyze the formation and release of new soluble oligomers from monomers at their surfaces, making fibrils a dynamic reservoir and catalyst for toxic oligomer species (Dear et al., 2024; Xu et al., 2024). This process is described by the following equation:

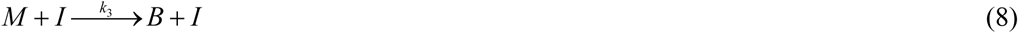

By formulating the conservation of TTR monomers in the blood plasma and normalizing the equation by

*V_plasma_*, the following equation is obtained:

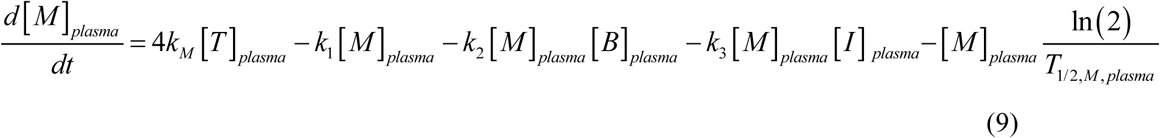

where [*T*]*_plasma_* is a constant value given by Eq. (4). The first term on the right-hand side represents the production of TTR monomers through the dissociation of tetramers. The second and third terms account for the reduction in monomer concentration due to their conversion into oligomers via nucleation and autocatalytic processes, respectively. The fourth term models the catalytic conversion of monomers into oligomers facilitated by the fibril surface. The fifth term captures the decline in monomer concentration due to their limited half-life. Accordingly, the model assumes that TTR monomers are cleared from the plasma either by aggregation into oligomers or through proteolytic degradation, consistent with (Criddle et al., 2020).

The conversion of free TTR oligomers into protofibrils is modeled analogously to coagulation processes occurring in colloidal suspensions (Boltachev and Ivanov, 2020). In this approach, free TTR oligomers (*B*) are characterized by a half-deposition time *θ*_1/ 2, *B*_, reflecting the time required for half of the oligomer population to incorporate into protofibrils. The total molar quantity of free TTR oligomers in the plasma is defined as the product of their molar concentration and the plasma volume. Normalizing the conservation equation by *V_plasma_* yields the following expression:

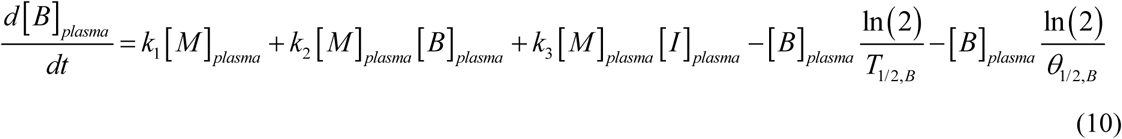

where the first and second terms on the right-hand side represent the formation of free TTR oligomers from monomers via nucleation and autocatalytic processes, respectively. The third term accounts for surface-mediated catalysis, representing the conversion of monomers to oligomers on the fibrillar lattice. The fourth term accounts for the decrease in free oligomer concentration due to their limited half-life, while the fifth term reflects their loss through deposition into protofibrils. While the transition from soluble oligomers to the fibrillar state is modeled using first-order kinetics, this does not imply that the resulting fibril is an oligomer. Instead, the first-order rate constant describes the sequestering of oligomer units into a solid-phase aggregate.

The conservation of TTR oligomers deposited into protofibrils circulating in the bloodstream, when normalized by *V_plasma_*, yields the following equation:

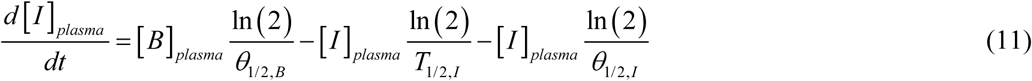

where the first term on the right-hand side represents the increase in the concentration of oligomers deposited into protofibrils as free oligomers incorporated into these protofibrils. The second term accounts for the reduction in protofibril concentration in the bloodstream due to their limited half-life, while the third term represents the decrease in their concentration due to capture by cardiac tissue.

To clarify the nature of the amyloid species in this model, [*I*] represents the molar concentration of oligomer units sequestered within fibrils, rather than the molar concentration of the fibrillar filaments themselves. In this kinetic framework, a fibril is defined as a continuous filament composed of these deposited oligomers. As this is a lumped-parameter kinetic model, it is designed to track the total mass of deposited protein (the amyloid burden) and does not explicitly resolve the length, branching, or stoichiometry of individual fibrillar polymers.

In the proposed scenario, protofibrils originate in the bloodstream, travel through circulation, and deposit in specific tissues with compatible binding sites. To enable this process, circulating protofibrils must strike a balance: they must be sufficiently small to move freely within the vasculature, yet large enough to expose a recognizable pattern of amino acid side chains that facilitate selective tissue adhesion. TTR protofibrils composed of approximately 10 to 50 monomer units appear to fulfill both of these criteria (Criddle et al., 2024).

In this scenario, the total molar concentration of all TTR species in the blood plasma, reported as the equivalent molar concentration of TTR monomers, is given by:

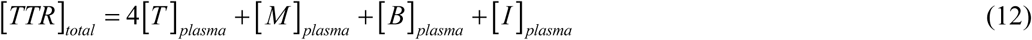

TTR concentrations expressed in μM (SI units) can be translated to clinical units (mg/dL) using the following conversion:

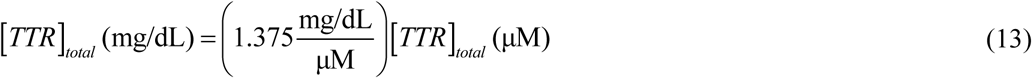

Stating the conservation of TTR oligomers deposited into fibrils that infiltrated the cardiac tissue, upon normalization by *V_heart_*, leads to the following equation:

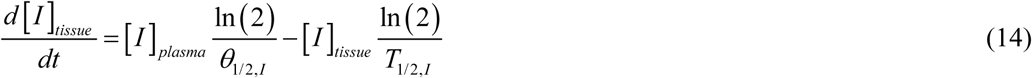

where the first term on the right-hand side models the increase in the concentration of TTR fibrils in cardiac tissue due to the capture of circulating protofibrils and their infiltration into the cardiac tissue. The second term accounts for a potential decrease in concentration of deposited fibrils resulting from their degradation. It is important to note that this degradation term reflects the anticipated effects of future therapeutic interventions—such as monoclonal antibody treatments—since current therapies are unable to remove pre-existing amyloid deposits from the heart (Emdin et al., 2023).

Eqs. (2), (9)–(11), and (14) are solved using initial condition (3) along with the following additional initial conditions:

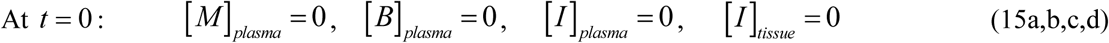

The numerical solution methodology is detailed in Section S1 of Supplementary Materials.

#### 2.1.3. Approximate analytical solution for the scenario in which amyloid fibrils form in blood plasma

Approximate solutions are derived for the case in which fibrillar species are inert, in the sense that they cannot further catalyze monomer conversion to amyloid form (*k*_3_ = 0). In addition, all half-lives are assumed to be infinitely long, while half-deposition times remain finite.

Assuming that 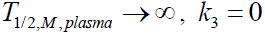, and considering the steady-state conditions, Eq. (9) can be rewritten as:

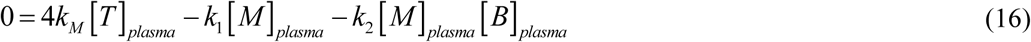

Assuming that *T*_1/ 2,*B*_ →∞ and *k*_3_ = 0, Eq. (10) can be rewritten as:

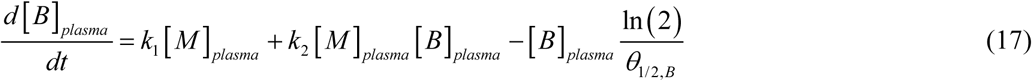

By adding Eqs. (16) and (17), the following is obtained:

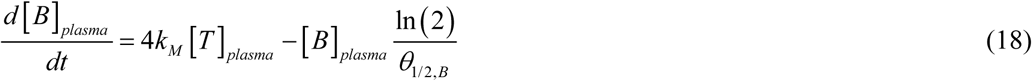

Solving Eq. (18) subject to initial condition (15b) yields:

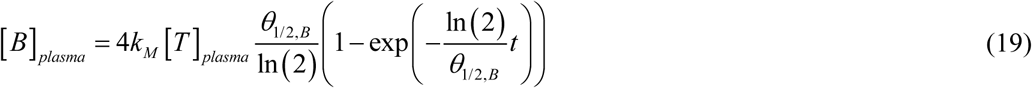

By substituting Eq. (19) into Eq. (16) and assuming steady-state conditions (*t* →∞), the following approximate solution for [*M*]*_plasma_* is obtained:

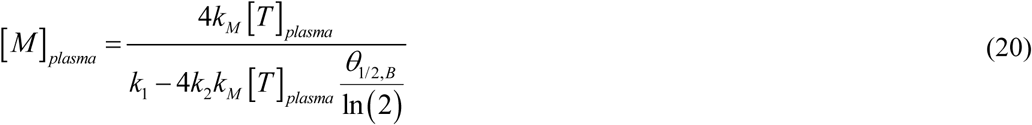

Assuming that *T*_1/ 2,*I*_ →∞, Eq. (11) can be rewritten as:

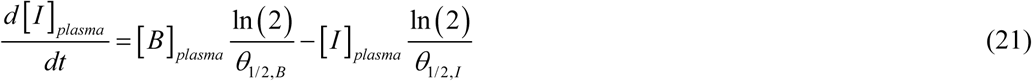

By substituting Eq. (19) for [*B*]*_plasma_* (*t*) into Eq. (21) and solving Eq. (21) with initial condition (15c), the following is obtained:

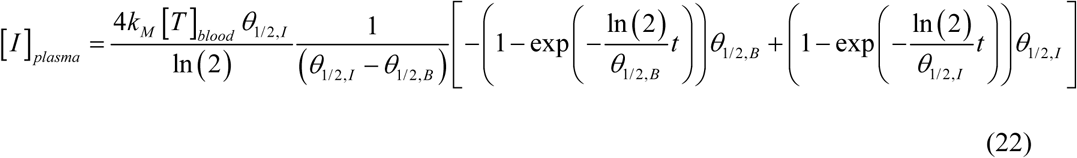

Note thats

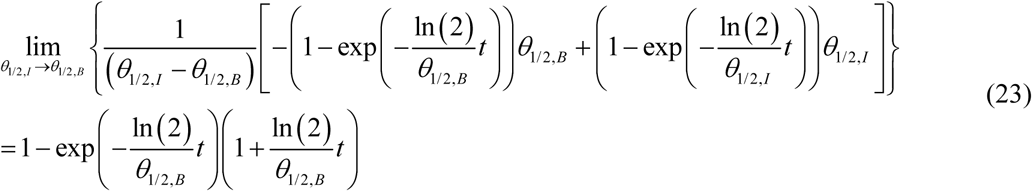

This implies that for *θ*_1/2,*I*_ → *θ*_1/2,*B*_, Eq. (22) simplifies to:

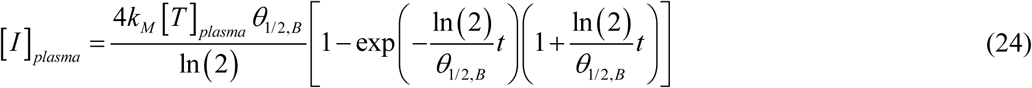

For *T*_1/ 2,*I*_ →∞ Eq. (14) simplifies to the following form:

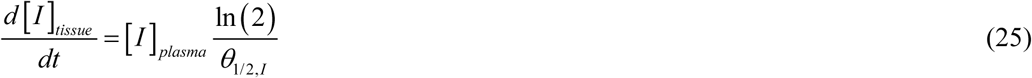

Substituting Eq. (24) for [*I*]*_plasma_* (*t*) into Eq. (25) and solving it with the initial condition (15d) yields the following result:

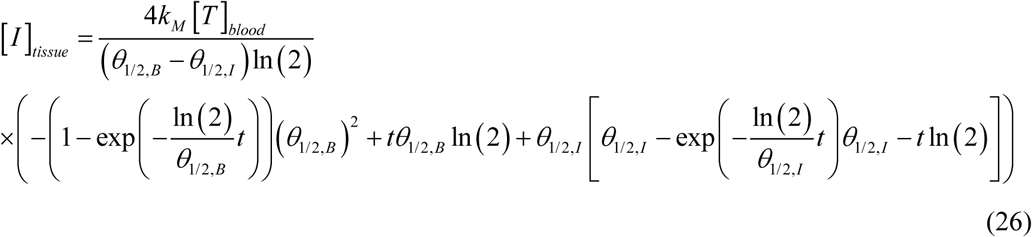

It is worth noting that:

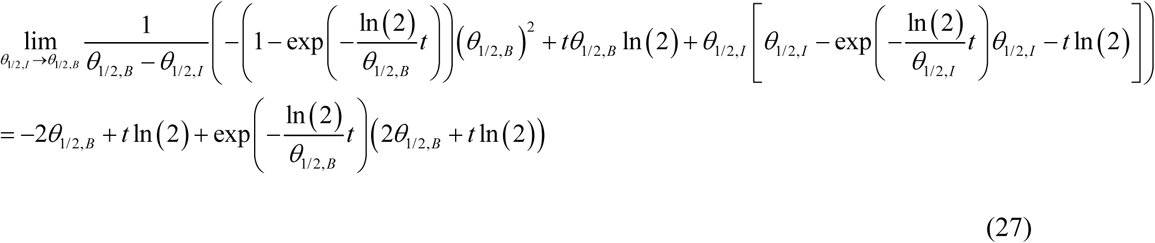

Thus for *θ*_1/2,*I*_ → *θ*_1/2,*B*_, Eq. (26) reduces to the following simplified form:

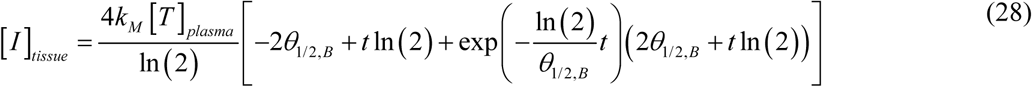

For large values of *t*

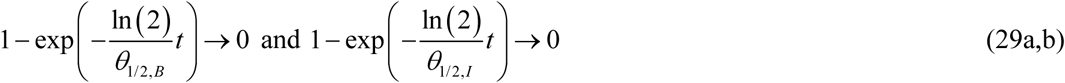

Thus, for large *t*, Eq. (26) approaches the following asymptotic form:

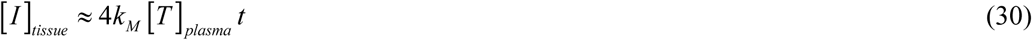

#### 2.1.4. Conservation equations for the scenario when amyloid fibrils form in cardiac muscle tissue

The second scenario is supported by a review of (Williams et al., 2022), who stated that TTR monomers can be taken up by myocardial tissue, where they aggregate and form amyloid fibrils.

Stating the conservation of TTR monomers in the blood plasma and dividing both sides of the equation by *V_plasma_*, yields the following equation:

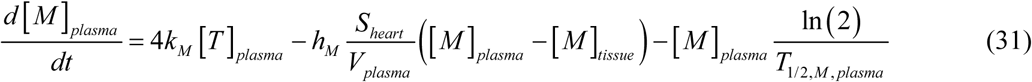

The total molar concentration of all TTR species in the blood plasma, reported as the equivalent molar concentration of TTR monomers, under this scenario is given by:

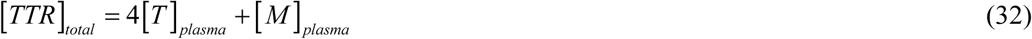

To align the modeled values with clinical reference ranges, TTR concentration can be converted from μM to mg/dL using Eq. (13).

By applying the principle of conservation of TTR monomers in cardiac tissue and normalizing both sides of the equation by *V_heart_*, which is defined as the heart volume prior to TTR fibril infiltration, the resulting equation is

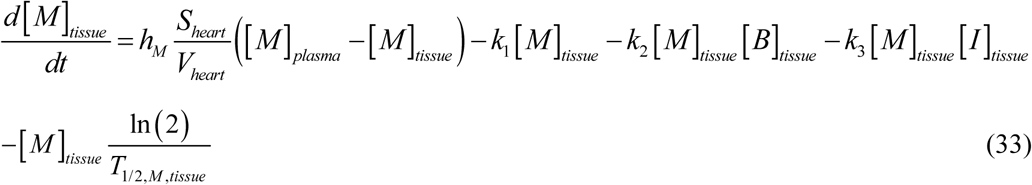

The first term on the right-hand side represents the volumetric rate of TTR monomer deposition into cardiac tissue, where 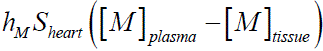 characterizes the total rate of monomer deposition into the heart. The second and third terms represent the loss of monomers as they are transformed into oligomers through nucleation and autocatalytic mechanisms, respectively. The fourth term accounts for surface-mediated catalysis, representing the rate at which existing fibrils facilitate the conversion of monomers into oligomeric species. The fifth term reflects monomer clearance resulting from their finite half-life.

Applying the conservation principle for TTR oligomers in cardiac tissue and scaling the equation by *V_heart_* yields the following equation:

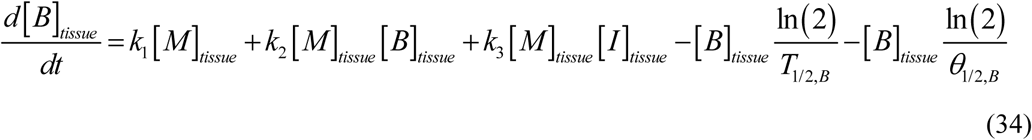

where the first and second terms on the right-hand side describe the generation of free TTR oligomers from monomers through nucleation and autocatalytic pathways, respectively. The third term describes the catalytic effect of the fibril surface on the conversion of monomers into oligomers. The fourth term represents the clearance of oligomers due to their finite lifespan, and the fifth captures their removal through incorporation into TTR fibrils.

Applying the conservation principle to TTR oligomers incorporated into fibrils and normalizing by *V_heart_* yields the following equation:

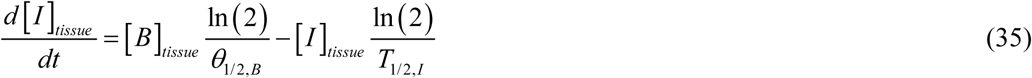

It should be noted that TTR fibrils are highly stable once deposited in cardiac tissue, and currently, heart transplantation is the only available treatment for patients with extensive fibril infiltration (Concetta Di Nora et al., 2022). Accordingly, the last term on the right-hand side of Eq. (35) is included solely to represent the potential impact of future therapies that may enable the clearance of TTR fibrils from the affected heart.

The system of Eqs. (31), (33)–(35) is solved subject to the initial conditions specified below:

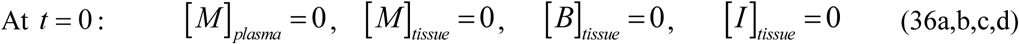

#### 2.1.5. Approximate analytical solution for the scenario in which amyloid fibrils form in cardiac muscle tissue

Similar to Section 2.1.3, approximate solutions are obtained under the assumption that fibrillar species cannot further catalyze monomer conversion to amyloid form (*k*_3_ = 0). In addition, all half-lives are assumed to be infinitely long, while half-deposition times remain finite.

Assuming that *T*_1/2,*M*, *plasma*_ → ∞, *k*_3_ = 0, and considering the steady-state conditions, Eq. (31) can be simplified as:

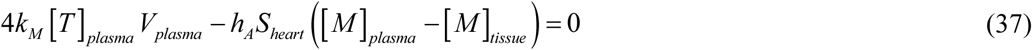

Assuming *T*_1/2,*M*, *tissue*_ →∞, *k*_3_ = 0, and applying steady-state conditions, Eq. (33) may be rewritten as:

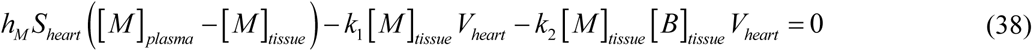

Combining Eqs. (37) and (38) yields the following equation:

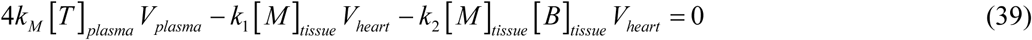

Dividing both sides of Eq. (39) by *V_heart_*, the equation can be rewritten as:

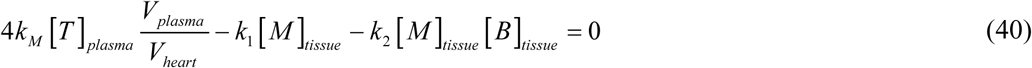

Assuming *T*_1/ 2,*B*_ →∞, Eq. (34) can be rewritten as:

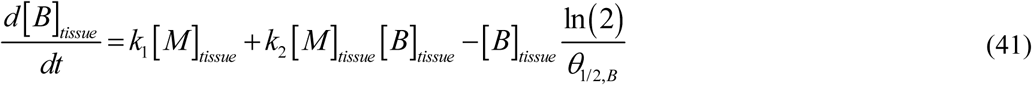

Adding Eqs. (40) and (41) yields the following result:

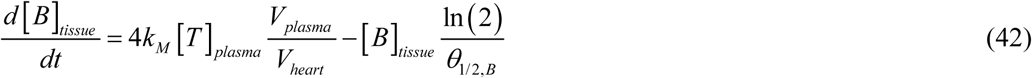

Solving Eq. (42) subject to initial condition (36c) yields:

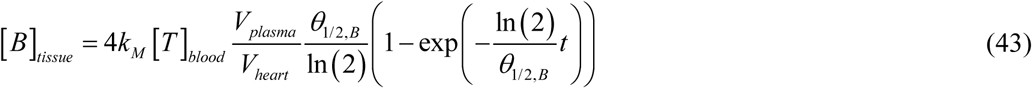

Substituting Eq. (43) into Eq. (40) and solving for [*M*]*_tissue_* yields:

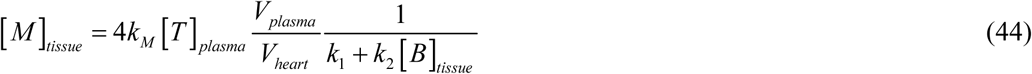

Rearranging Eq. (37) to solve [*M*]*_plasma_* yields the following equation:

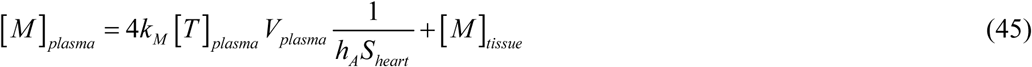

Assuming *T*_1/ 2,*I*_ →∞, Eq. (35) can be rewritten as:

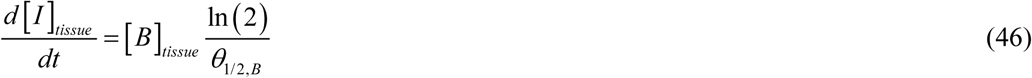

Substituting Eq. (43) for [*B*]*_tissue_* into Eq. (46) and solving it with initial condition (36d) yields the following result:

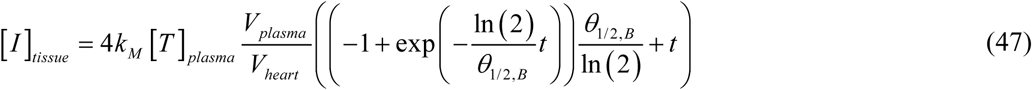

For large values of *t*

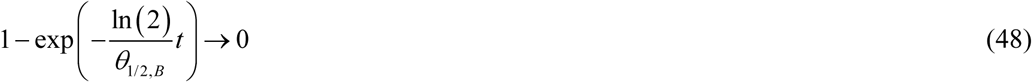

In the limit of large *t*, Eq. (47) converges to the following asymptotic expression:

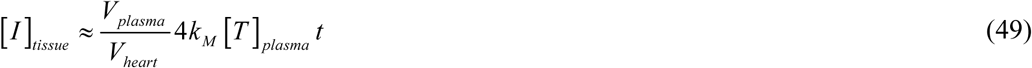

### 2.2. Calculating the volume occupied by TTR amyloid fibrils within the cardiac tissue

TTR amyloid fibrils are characterized by a twisted cross-β-helix structure (Miroy et al., 1996; Steinebrei et al., 2023; Dorbala et al., 2020; Schmidt et al., 2019).

The increase in the volume occupied by TTR amyloid fibril deposits in the cardiac tissue is determined by calculating the total number of TTR monomers incorporated into these fibril formations over a specified time *t*, *N_mon in fibrils_* (*t*). In the F-W model, monomers are assumed to have the same molecular weight as oligomers (see Eq. (6)); this consideration enables the number of incorporated monomers to directly represent the volume of the deposits. Applying the approach adapted from (Watzky et al., 2008) yields the following expression:

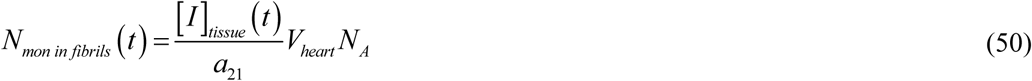

Here, *N_A_* denotes Avogadro’s number.

Another way to calculate *N_mon in fibrils_ (t)* is by employing the following equation, as outlined in (Watzky et al., 2008):

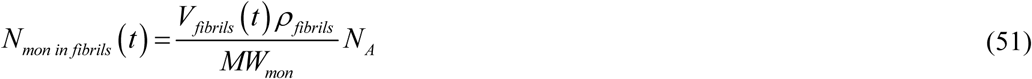

Here, *MW_mon_*refers to the molecular weight of a TTR monomer, *V_fibrils_* (*t*) denotes the volume occupied by TTR amyloid fibril deposits in the heart at time *t*, and *ρ _fibrils_* specifies their density.

Equating the expressions on the right side of Eqs. (50) and (51) and solving for the fraction of baseline heart volume occupied by TTR amyloid fibrils yields the following equation:

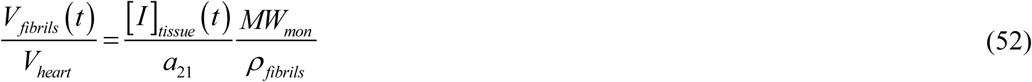

The total cardiac volume at time *t* can then be found as *V_fibrils_* (*t*) + *V_heart_*.

#### 2.2.1. Volume occupied by TTR amyloid fibrils for the scenario when amyloid fibrils form in blood plasma

In the scenario where amyloid fibrils form in the blood plasma, [*I*]*_tissue_* is given by Eq. (26), which leads to:

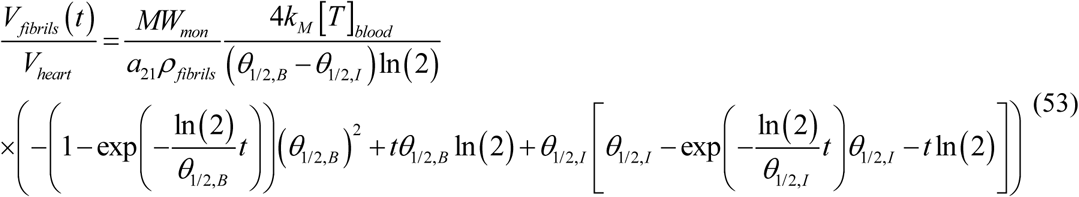

For the case where *θ*_1/2,*I*_ → *θ*_1/2,*B*_, Eq. (53) can be further simplified to:

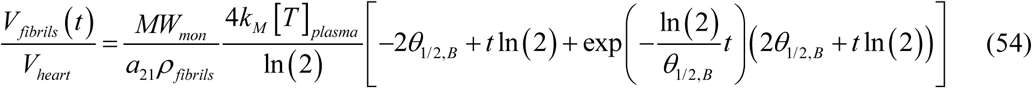

For large *t*, the asymptotic limit of [*I*]*_tissue_* is given by Eq. (30), which leads to the following asymptotic form of Eq. (53):

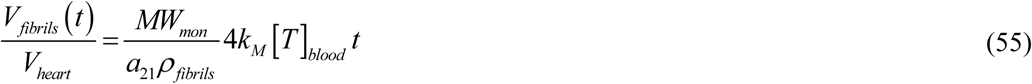

#### 2.2.2. Volume occupied by TTR amyloid fibrils for the scenario when amyloid fibrils form in cardiac muscle tissue

In the scenario where amyloid fibrils form in cardiac muscle tissue, [*I*]*_tissue_* is given by Eq. (47), which leads to:

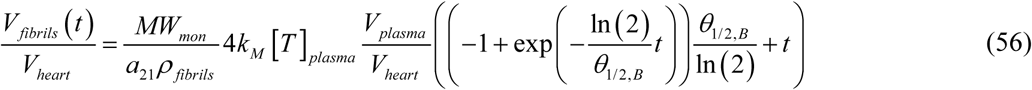

At large values of [*I*]*_tissue_*, the asymptotic behavior of [*I*]*_tissue_* is described by Eq. (49), resulting in the following simplified form of Eq. (56):

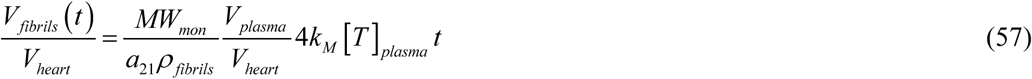

It is important to note that the main distinction between Eqs. (55) and (57) is the inclusion of the factor 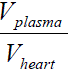 in Eq. (57). Using the parameter values listed in Table 2, this factor is estimated to be 5.21. This suggests that in scenario 2—where TTR monomers first deposit in the heart and aggregate locally within the cardiac tissue—the accumulation of TTR fibrils in the heart is approximately five times greater than in scenario 1, where amyloid fibrils form in the blood plasma and subsequently deposit into the cardiac tissue. This can be explained by the fact that in scenario 2, the concentration of TTR oligomers—which autocatalyze their formation from monomers—is higher than in scenario 1. This is due to the smaller volume of the heart compared to the volume of blood plasma, resulting in a higher local concentration of oligomers in scenario 2.

The biological age of the heart (referred to hereafter as biological age) is defined as follows:

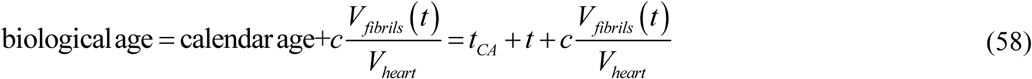

Simulations for the scenario in which amyloid fibril formation occurs in cardiac muscle tissue, using the following parameter values: 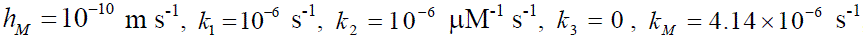, 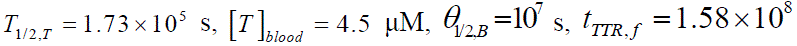 s(5 years), along with values of other parameters given in Table 2, show that 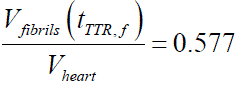. This result falls within the range reported by (Mehta et al., 2019), who classified patients into two groups based on their cardiac amyloidosis burden: moderate (0%–49%) and high (≥50%).

Assuming that TTR amyloidosis begins at age 60, progresses over 5 years, and that the patient’s biological age reaches 100 by age 65, the following estimate for parameter *c* in Eq. (58) is obtained:

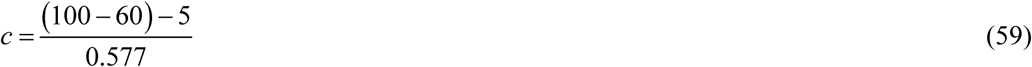

In this model, cardiac injury is quantified by the ratio of amyloid fibril volume to initial myocardial volume, 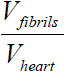. ‘Biological age’ is defined as a linear function of 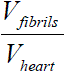, where the scaling constant *c* is calibrated such that terminal disease stages correspond to a biological age of 100 years. Maintaining a consistent *c* value for both plasma-aggregation and myocardial-aggregation models allows for a direct comparison of their respective physiological impacts, ensuring that higher TTR deposition consistently translates to an advanced biological age regardless of the site of fibrillogenesis.

In the scenario where amyloid fibrils form in the blood plasma, the volumetric ratio of deposited TTR fibrils to baseline myocardial volume at large *t* is given by Eq. (55), and the biological age approaches the following asymptotic expression:

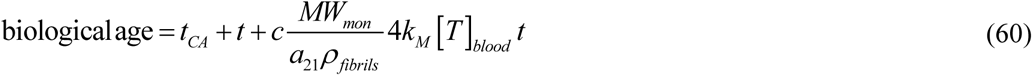

In the scenario where amyloid fibrils develop within the cardiac muscle tissue, Eq. (57) describes the volumetric ratio of deposited TTR fibrils to baseline myocardial volume as *t* becomes large, and the biological age tends toward the following asymptotic form:

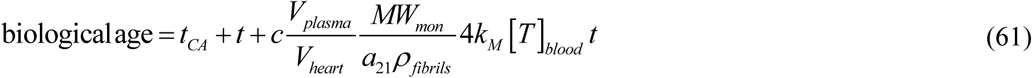

### 2.3. Sensitivity of the heart volume fraction occupied by TTR amyloid fibrils to various parameters

The sensitivity of the heart volume fraction occupied by TTR amyloid fibrils with respect to key model parameters was analyzed by calculating local sensitivity coefficients. These coefficients represent the first-order partial derivatives of the output variable— the volumetric ratio of deposited TTR fibrils to baseline myocardial volume —with respect to various parameters, such as the dissociation rate constant of TTR tetramers into monomers, *k_M_*. The methodology used follows established approaches described in (Beck and Arnold, 1977; Zadeh and Montas, 2010; Zi, 2011; Kuznetsov and Kuznetsov, 2019). For instance, the sensitivity coefficient of the volume fraction with respect to *k_M_* was computed using the following expression:

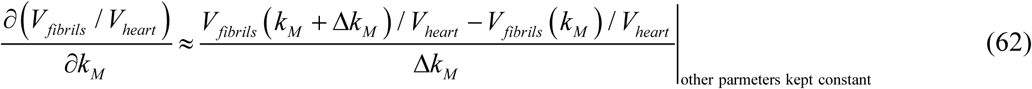

In this analysis, Δ*k* = 10^−3^ *k* represents the step size used for numerical differentiation. Multiple step sizes were tested to ensure that the computed sensitivity coefficients remained consistent and were not significantly affected by the choice of step size.

To facilitate comparison across parameters with different units and scales, dimensionless relative sensitivity coefficients were computed, following the approaches described in (Zadeh and Montas, 2010; Kuznetsov and Kuznetsov, 2019). For example:

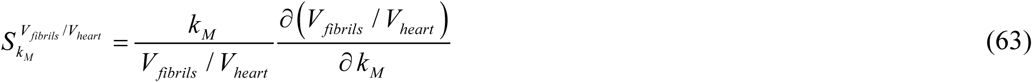

## 3. Results

The numerically and analytically computed dependencies of various TTR concentrations, the volumetric ratio of deposited TTR fibrils to baseline myocardial volume, and the predicted biological age are analyzed for both TTR aggregation scenarios—occurring in blood plasma and cardiac tissue—to determine which scenario yields more physiologically plausible predictions.

It is important to note that in all figures—except those illustrating sensitivity coefficients—the approximate analytical solutions show excellent agreement with the numerical results. Sensitivity coefficients, which represent partial derivatives of the solution with respect to model parameters, are highly sensitive to the accuracy of the underlying solution. As a result, some discrepancies between the analytically and numerically calculated sensitivity coefficients are observed at early time points.

The molar concentration of TTR tetramers, [*T*], increases rapidly until it reaches a steady-state value. This initial rapid increase occurs due to the assumed zero concentration of [*T*] at *t*=0. The steady-state concentration of TTR tetramers is governed by the rate constant for tetramer dissociation into monomers, *k_M_*. The value of *q_T_* is calibrated by setting *k_M_* in Eq. (5) to 4.14 × 10⁻⁶ s⁻¹, a value reported in (Nelson et al., 2021). At this value, the model yields a tetramer concentration of 4.5 µM, which lies at the midpoint of the physiological range reported in (Coelho et al., 2016). An order-of-magnitude increase in *k_M_* (to 4.14 × 10⁻^5^ s⁻¹) leads to a fivefold decrease in the steady-state concentration of TTR tetramers, whereas an order-of-magnitude decrease in *k_M_* (to 4.14 × 10⁻^7^ s⁻¹) results in a twofold increase in tetramer concentration (Fig. S1). Tetramer concentrations are identical in the scenarios of TTR accumulation in blood plasma and cardiac tissue (see Figs. S1a and S1b), as both are governed by the same Eq. (2) and initial condition (3).

Tafamidis is used in the treatment of TTR amyloidosis. Its effectiveness is attributed to its ability to bind to the native tetrameric form of TTR, thereby stabilizing the tetramer and preventing its dissociation into monomers, which are prone to aggregation (Maurer et al., 2018). The stabilizing effect of tafamidis on TTR tetramers is modeled by reducing the rate constant governing tetramer dissociation into monomers, *k_M_*. A lower *k_M_* leads to an increased concentration of tetramers (Fig. S1). This rise in native TTR (tetramer) concentration with decreasing *k_M_* is consistent with findings reported by (Monteiro et al., 2023).

The concentration of TTR monomers in blood plasma shows a sharp initial decline, caused by the assumed initial condition, and subsequently stabilizes at a low steady-state level (Fig. S2). This aligns with the observations of (Corino et al., 2025), who noted that TTR monomers are highly unstable *in vivo* and generally exist at very low plasma concentrations, [*M*]*_plasma_*. For the case of TTR aggregation in cardiac tissue the steady-state values of [*M*]*_plasma_* depend on the value of *k_M_*, with a larger value of *k_M_* corresponding to a larger value of steady-state concentration of [*M*]*_plasma_* (Fig. S2b).

In the scenario where TTR aggregates in the cardiac tissue, the concentration of TTR monomers in the tissue quickly reaches a low steady-state value after a brief transient period. This steady-state concentration is independent of the value of *k_M_* (Fig. S3).

The concentration of TTR oligomers, [*B*]*_plasma_* for scenario 1 and [*B*]*_tissue_* for scenario 2, increases over time until it reaches a steady-state level, which is higher for larger values of the tetramer dissociation constant, *k_M_* (Fig. S4). This can be explained by the fact that higher values of *k_M_* lead to increased production of monomers, which subsequently undergo conversion into oligomers.

Similarly, when TTR aggregates in plasma, the concentration of TTR oligomers incorporated into circulating amyloid protofibrils increases over time and eventually reaches a steady-state level, which is higher for larger values of *k_M_* (Fig. S5).

In the scenario where TTR monomers aggregate in plasma, the total concentration of TTR in blood plasma rises over time and eventually reaches very high levels. For *k_M_* = 4.14×10^−6^ s^-1^, the steady-state concentration of [*TTR*]*_total_* is approximately 3000 mg/dL (Fig. 2a), suggesting that this scenario is likely unrealistic. In this scenario, increasing *k_M_* results in a higher steady-state concentration of [*TTR*]*_total_*. This is because in this scenario the main components contributing to [*TTR*]*_total_* are TTR oligomers and fibrils circulating in the blood plasma. The concentrations of both components increase with increasing *k_M_* (see Figs. S4a and S5), as TTR monomers are progressively converted into oligomers, which then aggregate into fibrils.

**Fig. 2.**
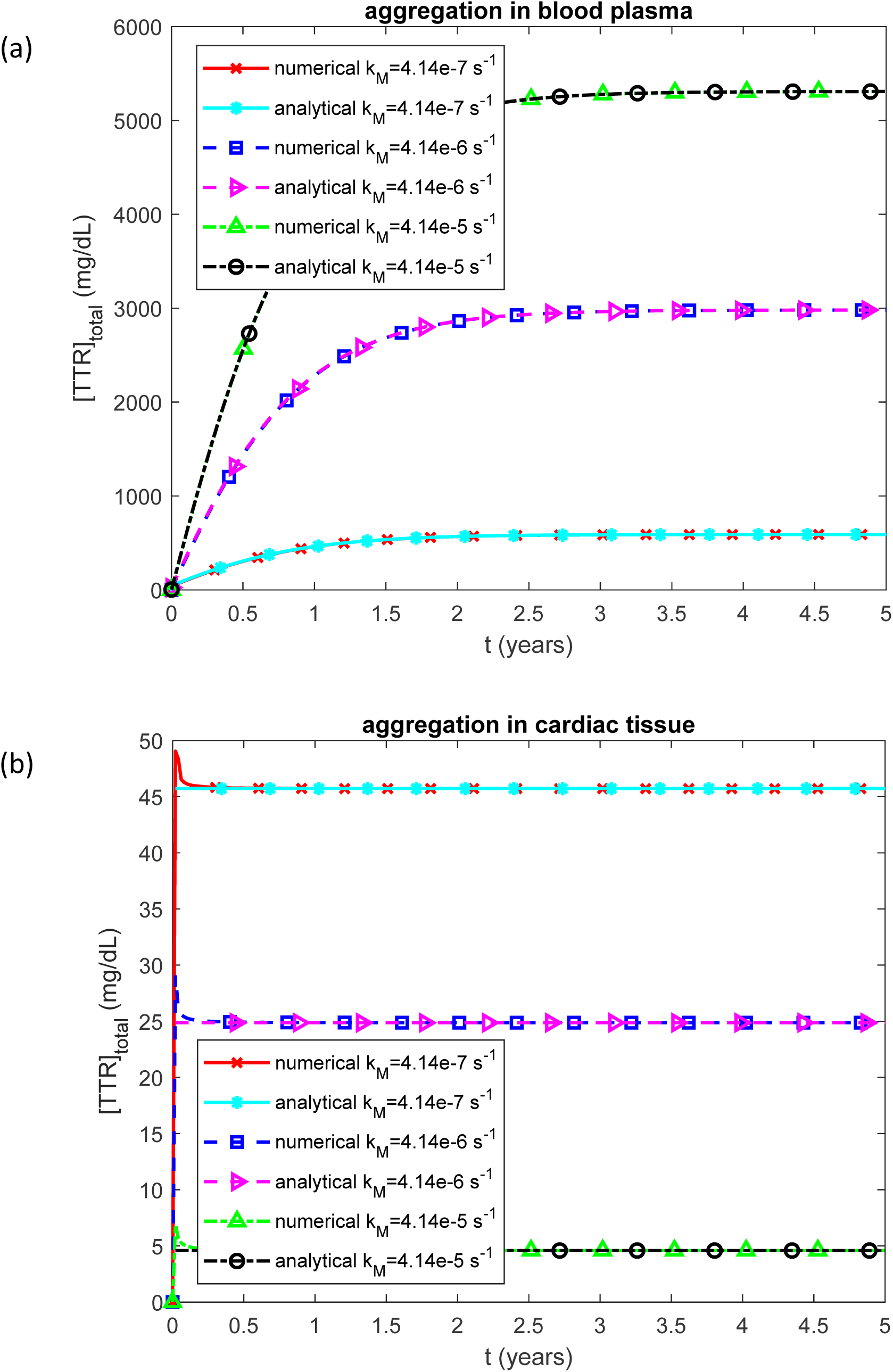
Total concentration of TTR (in any state) in the blood plasma vs time for various values of the rate constant describing the dissociation of TTR tetramers into monomers. (a) Scenario when TTR monomers misfold and assemble into protofibrillar formations within the bloodstream before depositing in the heart. (b) Scenario when TTR monomers remain in circulation until they directly attach to sites within the heart, where they form amyloid deposits *in situ*.

In contrast, when TTR aggregates in cardiac tissue, the total TTR concentration exhibits an initial jump due to the imposed initial condition and then rapidly declines to a steady-state value. At *k_M_* = 4.14×10^−6^ s^-1^, the predicted steady-state concentration of [*TTR*]*_total_* is approximately 25 mg/dL (Fig. 2b), falling within the reported physiological range of 18–45 mg/dL (Hood et al., 2022; Nativi-Nicolau, 2018). In this scenario, increasing *k_M_* leads to a decrease in the steady-state concentration of [*TTR*]*_total_*. This is because in this scenario the TTR tetramer concentration constitutes the main component of the total TTR in blood plasma (see Figs. S1b and 2b).

The concentration of TTR oligomers incorporated into amyloid fibrils within cardiac tissue initially rises slowly, then transitions to a linear increase over time (Fig. 3). The rate of this increase, characterized by the slope of the curve, becomes greater with higher values of *k_M_*. Similarly, the volumetric ratio of deposited TTR fibrils to baseline myocardial volume increases linearly following an initial period, suggesting that fibril accumulation in the heart is primarily governed by the rate at which TTR monomers are supplied through tetramer dissociation (Fig. 4).

**Fig. 3.**
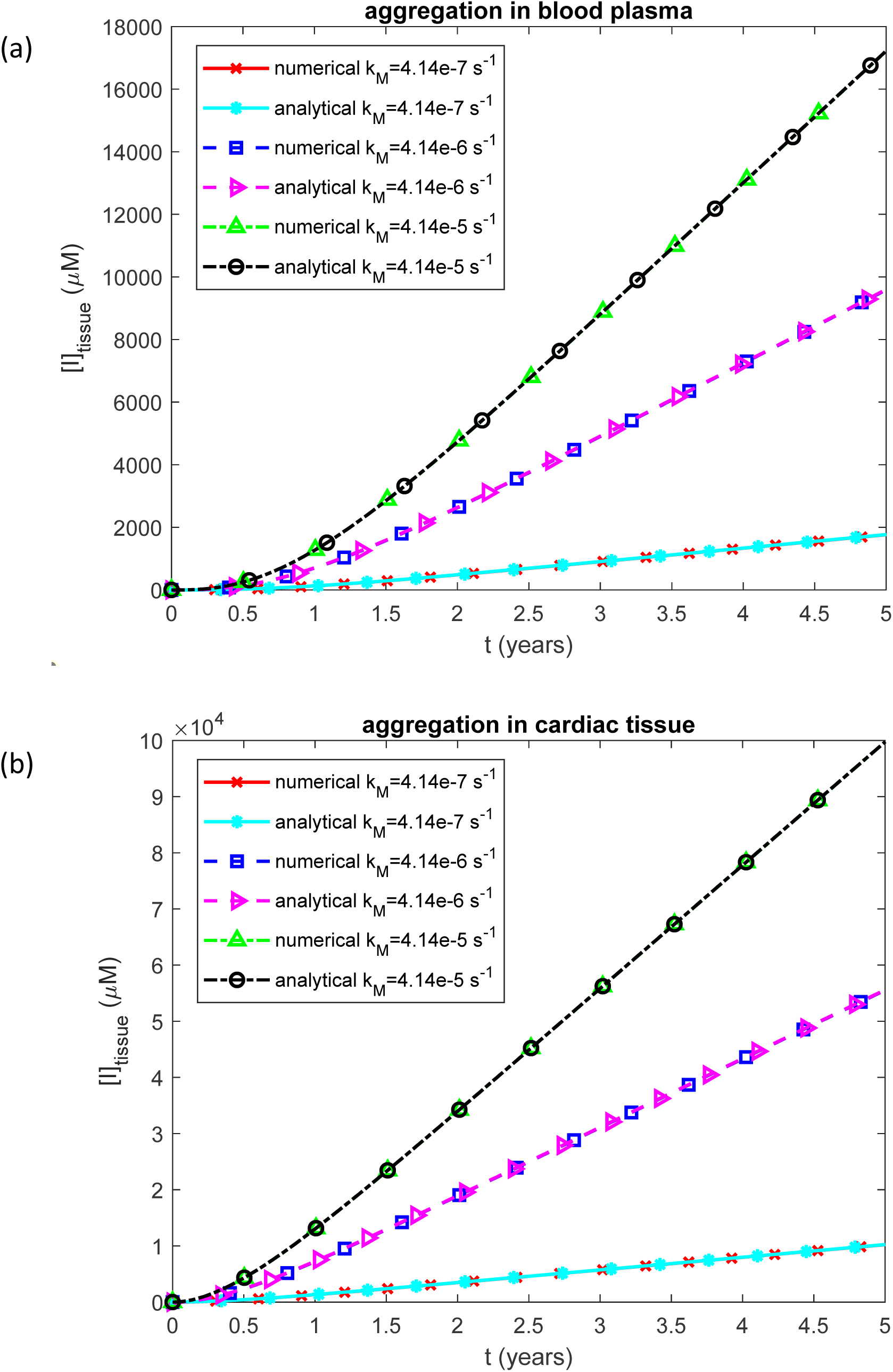
Molar concentration of TTR oligomers deposited into amyloid fibrils that infiltrated the cardiac tissue vs time for various values of the rate constant describing the dissociation of TTR tetramers into monomers. (a) Scenario when TTR monomers misfold and assemble into protofibrillar formations within the bloodstream before depositing in the heart. (b) Scenario when TTR monomers remain in circulation until they directly attach to sites within the heart, where they form amyloid deposits *in situ*.

**Fig. 4.**
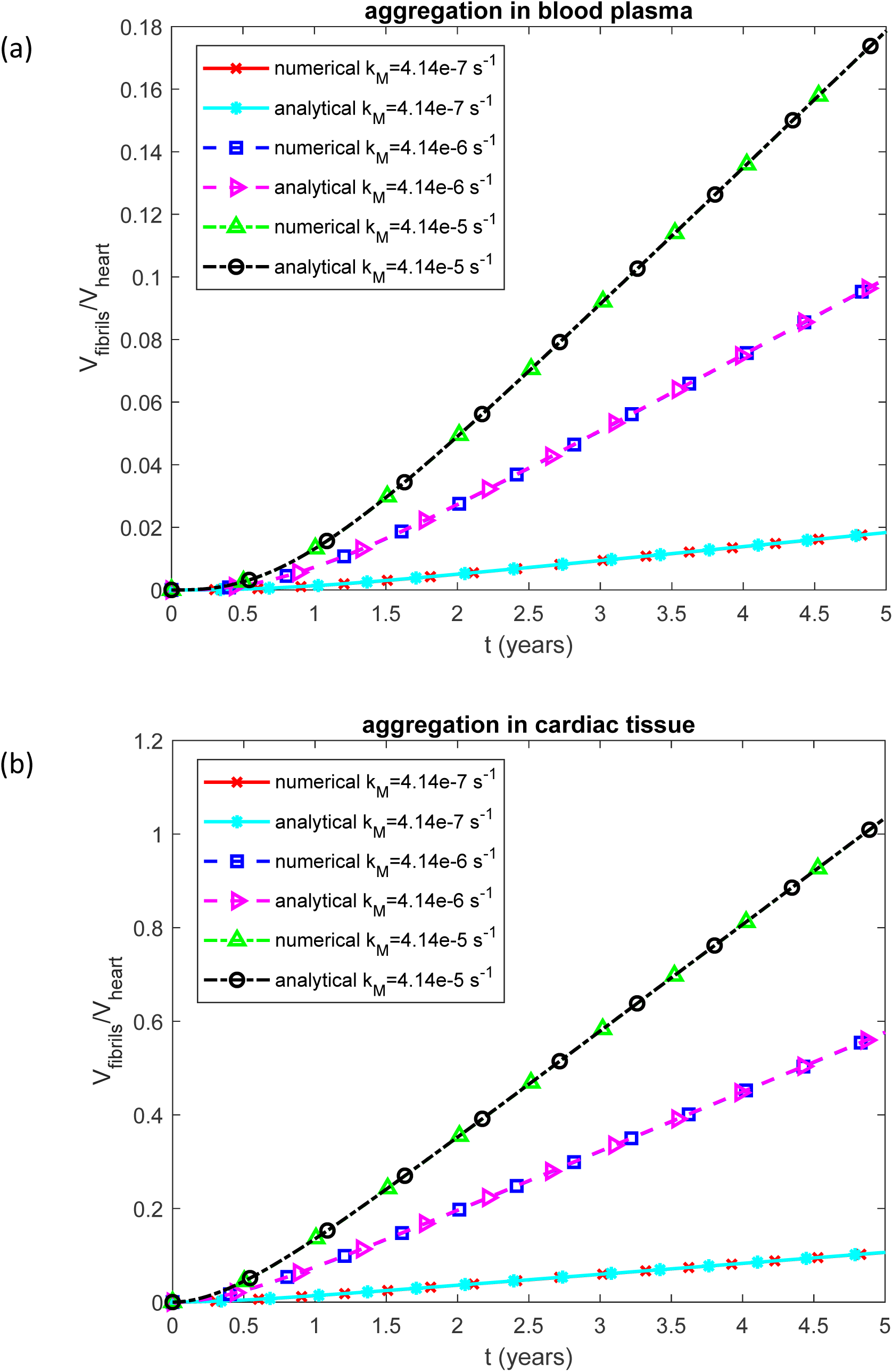
Volumetric ratio of deposited TTR fibrils to baseline myocardial volume vs time for various values of the rate constant describing the dissociation of TTR tetramers into monomers. (a) Scenario when TTR monomers misfold and assemble into protofibrillar formations within the bloodstream before depositing in the heart. (b) Scenario when TTR monomers remain in circulation until they directly attach to sites within the heart, where they form amyloid deposits *in situ*.

It is important to note that, as shown by the black curve in Fig. 4b (marked by empty circles), the predicted volume of TTR fibrils can, in some cases, exceed the initial (before TTR accumulation) volume of the heart. This observation aligns with Eq. (52), which indicates that the volume of amyloid fibrils incorporated into the heart is not constrained by the initial heart volume. Clinically, progressive accumulation of TTR amyloid fibrils within the myocardium leads to thickening of the heart walls and severe stiffening of the cardiac muscle, and patients typically die from heart failure once amyloid infiltration becomes advanced (Kocher et al., 2020; Bukhari et al., 2025).

Biological age increases more rapidly in the scenario where TTR fibrils aggregate within cardiac tissue compared to when fibrils aggregate in the blood plasma (compare Figs. 5b and 5a). In both scenarios, the rate of biological aging also rises with increasing values of *k_M_* (Fig. 5).

**Fig. 5.**
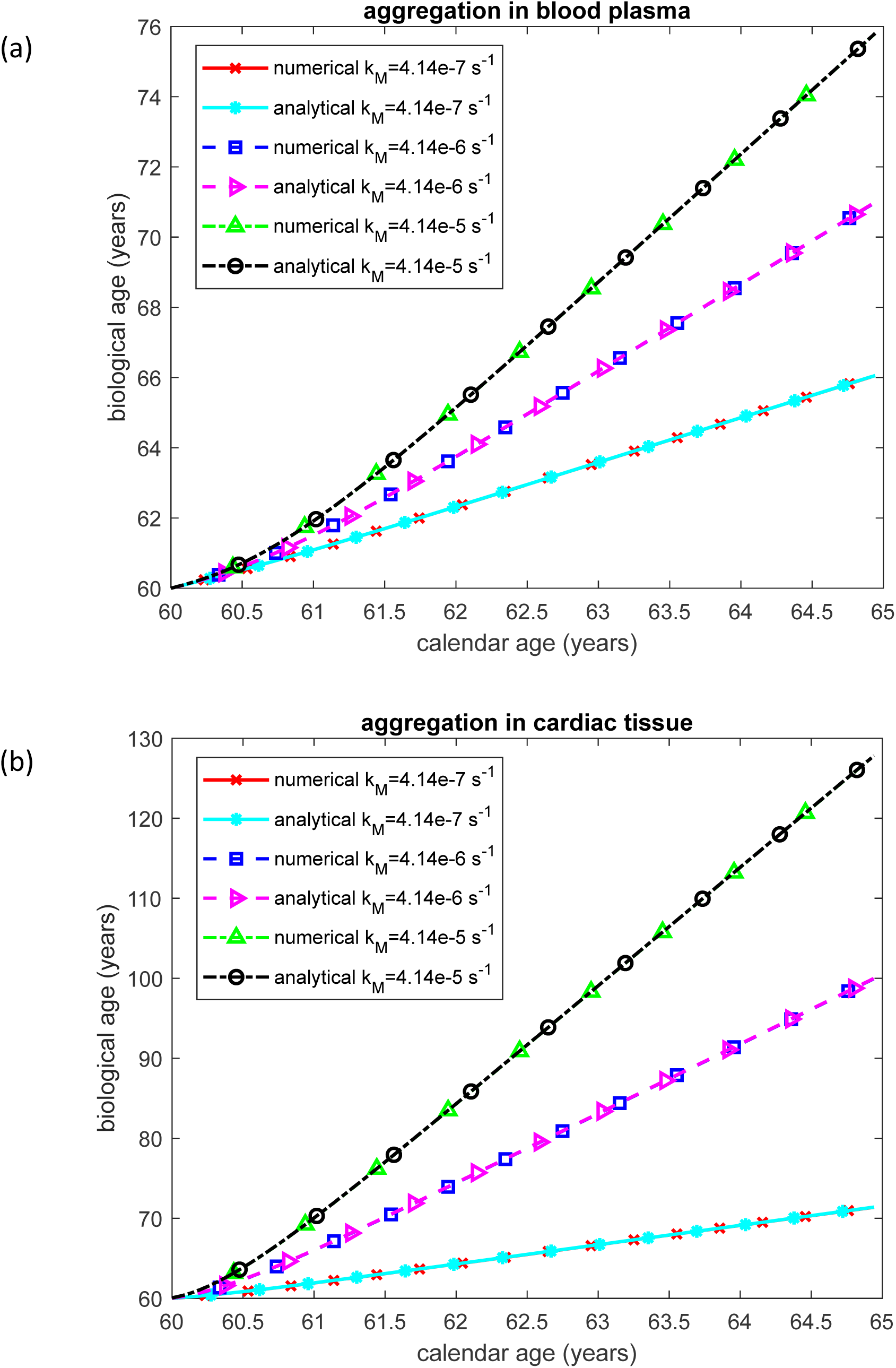
Biological age vs calendar age for various values of the rate constant describing the dissociation of TTR tetramers into monomers. (a) Scenario when TTR monomers misfold and assemble into protofibrillar formations within the bloodstream before depositing in the heart. (b) Scenario when TTR monomers remain in circulation until they directly attach to sites within the heart, where they form amyloid deposits *in situ*.

A decrease of *k_M_* thus leads to a reduction in the molar concentration of TTR oligomers deposited into amyloid fibrils within the cardiac tissue, [*I*]*_tissue_* (Fig. 3), the volumetric ratio of deposited TTR fibrils to baseline myocardial volume, *V_fibrils_* / *V_heart_* (Fig. 4), and the biological age (Fig. 5).

The molar concentration of TTR tetramers is independent of the half-deposition time of free TTR oligomers into fibrils. As in Fig. S1, the tetramer concentrations shown in Fig. S6 are identical for the scenarios of TTR aggregation in plasma (Fig. S6a) and cardiac tissue (Fig. S6b).

In both scenarios, the molar concentration of TTR monomers in blood plasma reaches a steady-state value that depends on the half-deposition time of free TTR oligomers into fibrils, *θ*_1/2,*B*_ (Fig. S7). A smaller value of *θ*_1/2,*B*_ corresponds to a higher steady-state concentration of monomers, a relationship that warrants a more detailed explanation.

In the scenario where TTR monomers aggregate into free oligomers in the bloodstream, this relationship arises because monomers convert into oligomers via two pathways: primary nucleation, which is slow, and secondary nucleation, which is faster and autocatalyzed by existing free oligomers. A decrease in *θ*_1/2,*B*_ lowers the concentration of free oligomers (Fig. S9a), slowing the autocatalytic conversion of monomers and thereby increasing their steady-state concentration.

In the alternative scenario—where TTR monomers form oligomers within the cardiac tissue—the effect is explained by a compensatory mechanism. At steady state, plasma monomers are deposited into cardiac tissue to replenish the local monomer pool, which is being depleted through conversion into oligomers. A smaller value of *θ*_1/2,*B*_ implies fewer oligomers in the tissue, leading to a lower conversion rate of monomers. Consequently, fewer monomers are drawn from the plasma into the tissue, resulting in a higher plasma monomer concentration.

In the scenario of TTR accumulation in cardiac tissue, the molar concentration of TTR monomers is also higher when the half-deposition time of free TTR oligomers into fibrils, *θ*_1/2,*B*_, is shorter (Fig. S8). A smaller *θ*_1/2,*B*_ results in fewer oligomers accumulating in the tissue (Fig. S9b), as they are more rapidly deposited into fibrils. This reduces the rate of autocatalytic conversion of monomers into oligomers, thereby leading to a higher concentration of monomers in the tissue.

The molar concentrations of TTR oligomers deposited into amyloid protofibrils circulating in the blood plasma reach the same steady-state value for different values of *θ*_1/2,*B*_ (compare the curves for *θ*_1/2,*B*_ = 10^5^ s and *θ*_1/2,*B*_ = 10^7^ s in Fig. S10). However, a smaller value of *θ*_1/2,*B*_ leads to a faster approach to steady state, as it corresponds to more rapid deposition of free oligomers into fibrils.

When TTR aggregation is assumed to occur in plasma, the total TTR concentration rises continuously and eventually reaches unrealistically elevated levels. For example, at *θ*_1/2,*B*_ =10^7^ s, the model predicts a steady-state concentration of [*TTR*]*_tissue_* to be approximately 3000 mg/dL (Fig. 6a), which far exceeds values observed *in vivo*, suggesting that this scenario is unlikely to reflect physiological reality.

**Fig. 6.**
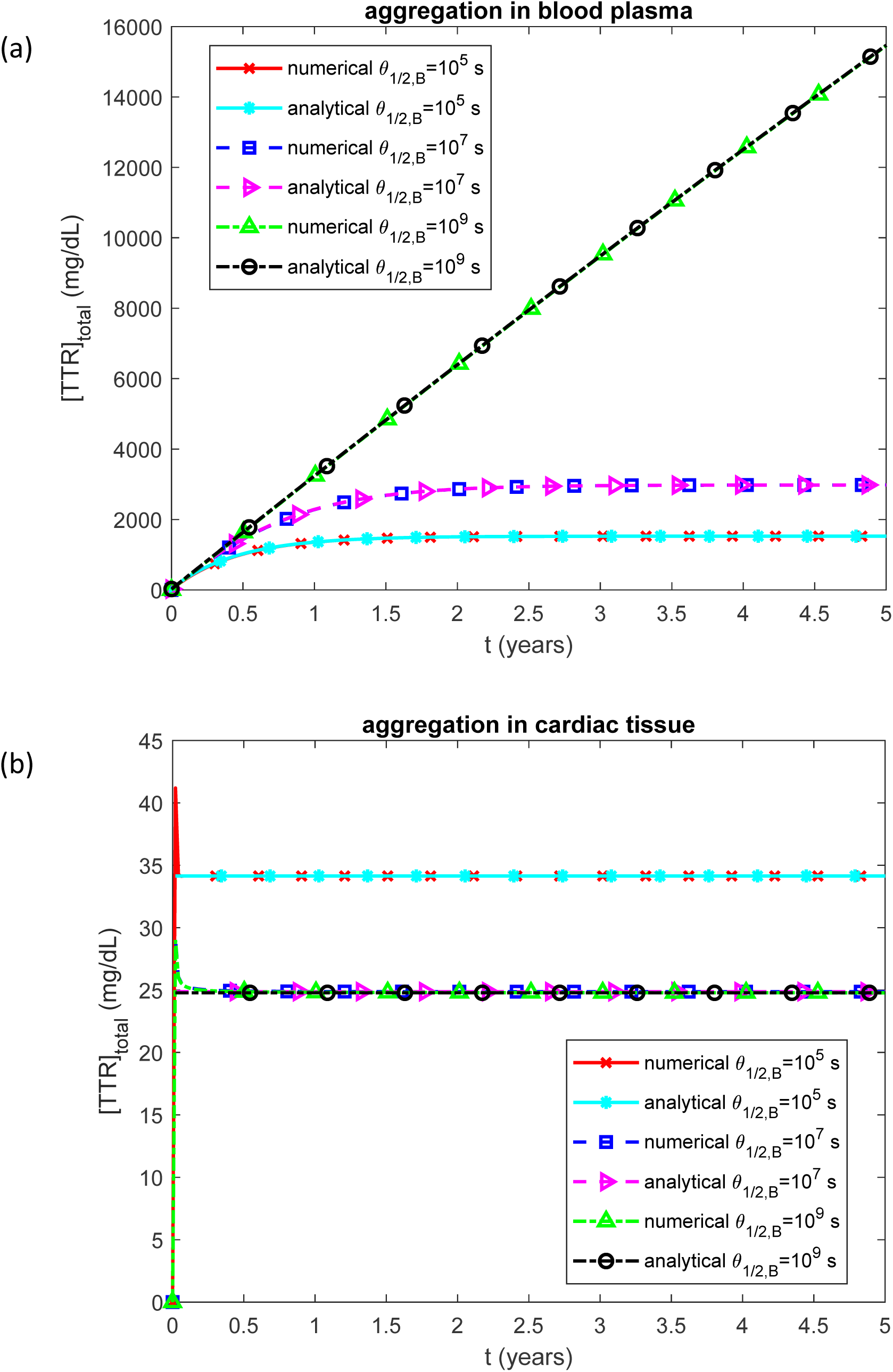
Total concentration of TTR (in any state) in the blood plasma vs time for various values of the half-deposition time for free TTR oligomers to become incorporated into TTR fibrils. (a) Scenario when TTR monomers misfold and assemble into protofibrillar formations within the bloodstream before depositing in the heart. (b) Scenario when TTR monomers remain in circulation until they directly attach to sites within the heart, where they form amyloid deposits *in situ*.

The rise of steady-state total TTR concentration with increasing *θ*_1/2,*B*_ is primarily driven by the accumulation of TTR oligomers, which make up the dominant fraction of total TTR in the blood plasma in this scenario (see Figs. 6a, S9a, and S10a). As *θ*_1/2,*B*_ increases, oligomer concentrations also rise (Fig. S9a), since the slower deposition of oligomers into fibrils allows them to persist longer in the plasma.

In the scenario where TTR aggregates in cardiac tissue, the total TTR concentration, [*TTR*]*_total_*, shows an initial spike due to the imposed initial condition and then quickly declines to a steady-state level. As shown in Fig. 6b, steady-state TTR concentrations are in excellent agreement with the 18–45 mg/dL range reported in the literature (Hood et al., 2022; Nativi-Nicolau, 2018). In this scenario, increasing *θ*_1/2,*B*_ leads to a decrease in the steady-state concentration of total TTR. This is because in this scenario total TTR consists of the concentrations of circulating TTR tetramers and monomers (see Figs. S6b and S7b). While the tetramer concentration remains unaffected by changes in *θ*_1/2,*B*_ (Fig. S6b), the monomer concentration decreases as *θ*_1/2,*B*_ increases (Fig. S7b). This occurs because monomer-to-oligomer conversion is autocatalyzed by oligomers; thus, increasing the oligomer half-life raises their steady-state levels, enhancing the catalytic conversion of monomers (Fig. S9b).

The molar concentration of TTR oligomers deposited into amyloid fibrils that have infiltrated the cardiac tissue, [*I*]*_tissue_*, is approximately five times higher in the scenario where TTR aggregation occurs within the cardiac tissue compared to when aggregation takes place in the plasma (Fig. 7). A smaller value of *θ*_1/ 2,*B*_ leads to a higher [*I*]*_tissue_* because in this case the deposition of free oligomers into fibrils occurs faster.

**Fig. 7.**
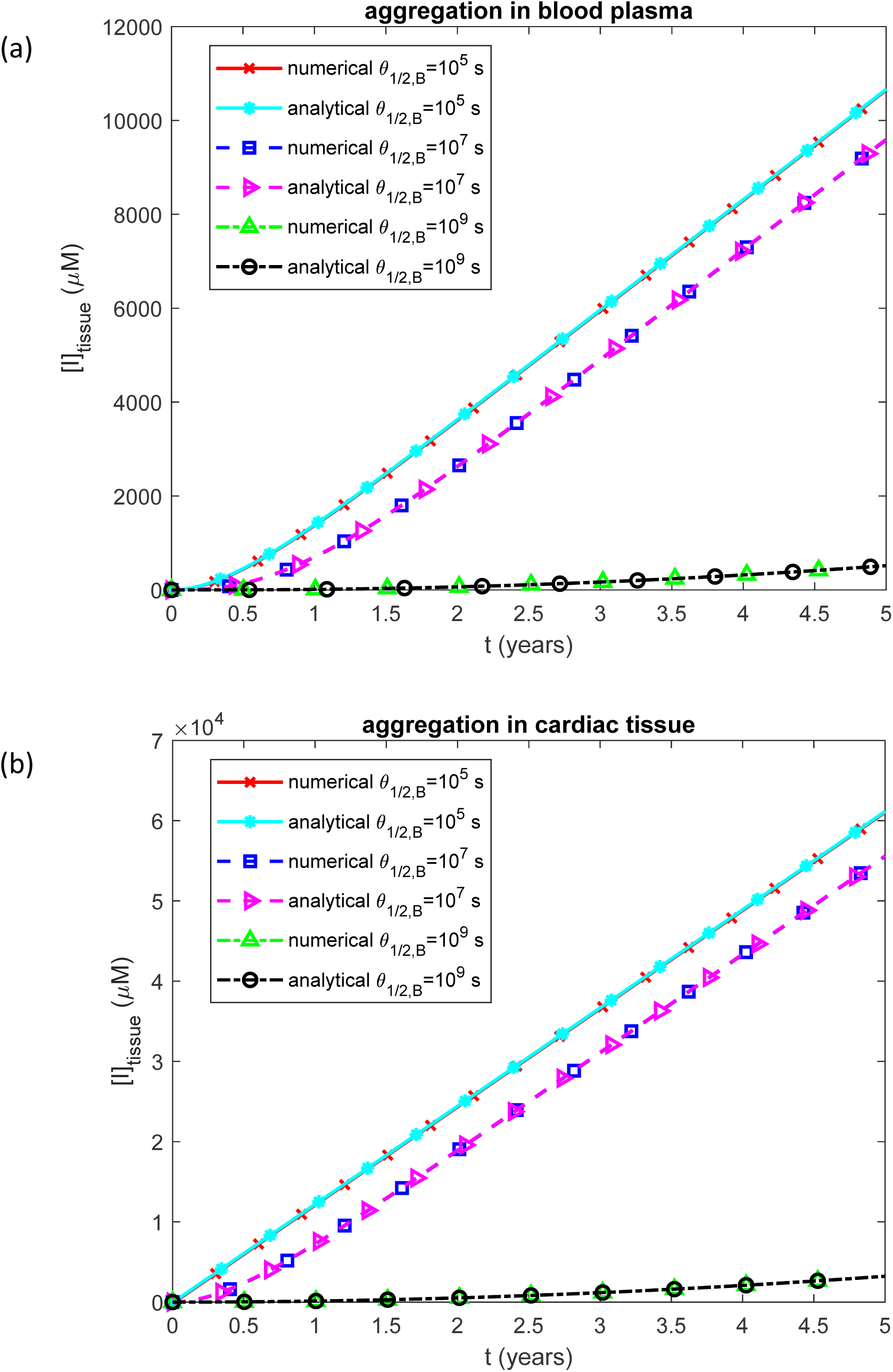
Molar concentration of TTR oligomers deposited into amyloid fibrils that infiltrated the cardiac tissue vs time for various values of the half-deposition time for free TTR oligomers to become incorporated into TTR fibrils. (a) Scenario when TTR monomers misfold and assemble into protofibrillar formations within the bloodstream before depositing in the heart. (b) Scenario when TTR monomers remain in circulation until they directly attach to sites within the heart, where they form amyloid deposits *in situ*.

The volumetric ratio of deposited TTR fibrils to baseline myocardial volume follows a similar trend (Fig. 8). Notably, for *θ*_1/2,*B*_ = 10^7^ s, after five years of disease progression the volumetric ratio of deposited TTR fibrils to baseline myocardial volume reaches 58% in the tissue aggregation scenario, whereas it reaches only 10% when aggregation occurs in the plasma. This difference arises because the plasma volume is significantly larger than the volume of the heart. As a result, the local concentration of free oligomers—the drivers of autocatalysis—is much higher in the heart in the scenario when aggregation takes place there, compared to when aggregation occurs in the plasma.

**Fig. 8.**
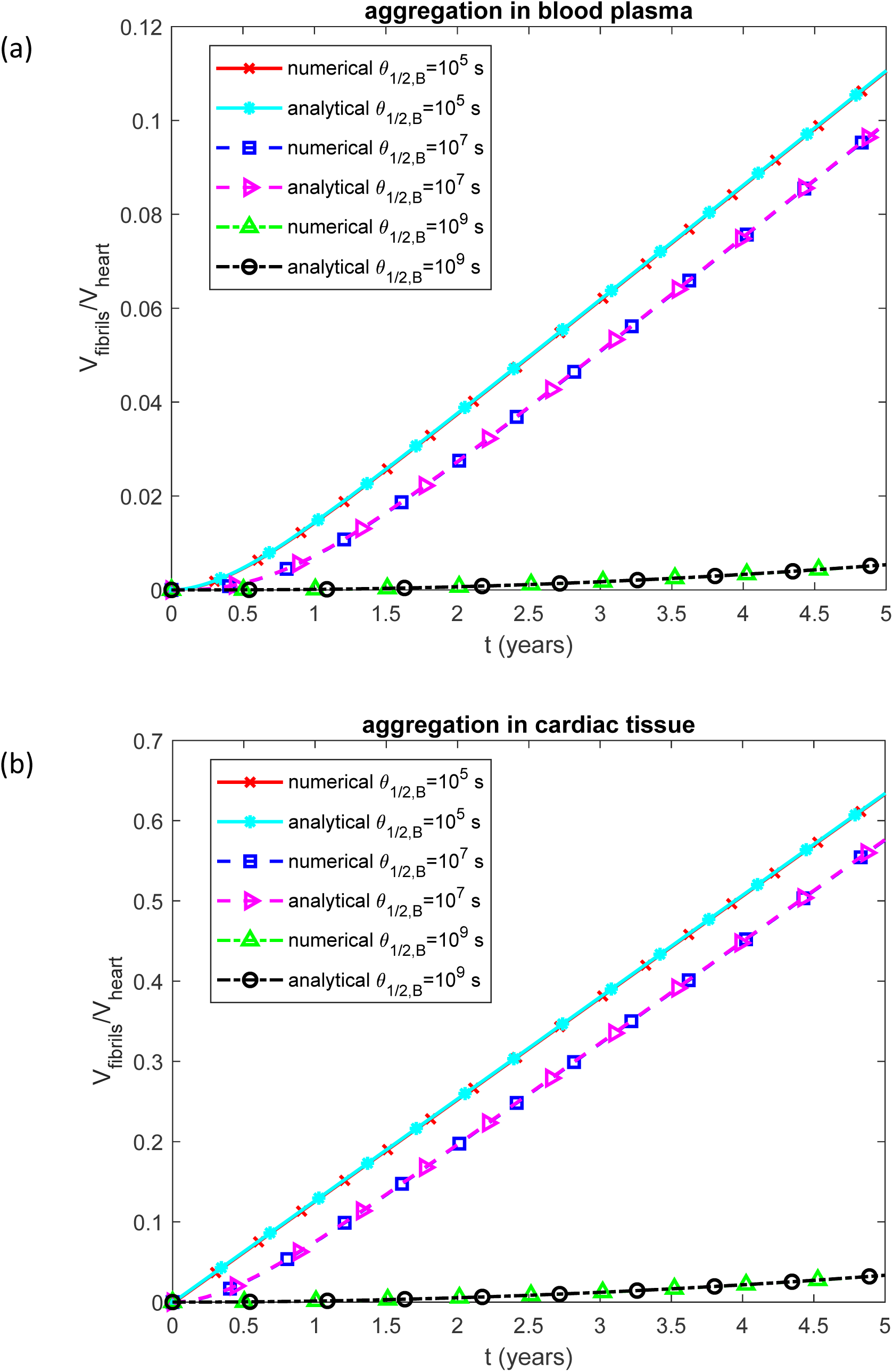
Volumetric ratio of deposited TTR fibrils to baseline myocardial volume vs time for various values of the half-deposition time for free TTR oligomers to become incorporated into TTR fibrils. (a) Scenario when TTR monomers misfold and assemble into protofibrillar formations within the bloodstream before depositing in the heart. (b) Scenario when TTR monomers remain in circulation until they directly attach to sites within the heart, where they form amyloid deposits *in situ*.

Biological age is calculated using Eq. (58) (Fig. 9). Prior to the onset of TTR amyloidosis—assumed to begin at age 60—biological and calendar ages are identical. However, following the disease onset, biological age advances more rapidly than calendar age due to the accelerated aging effect caused by amyloid fibril deposition in the heart.

**Fig. 9.**
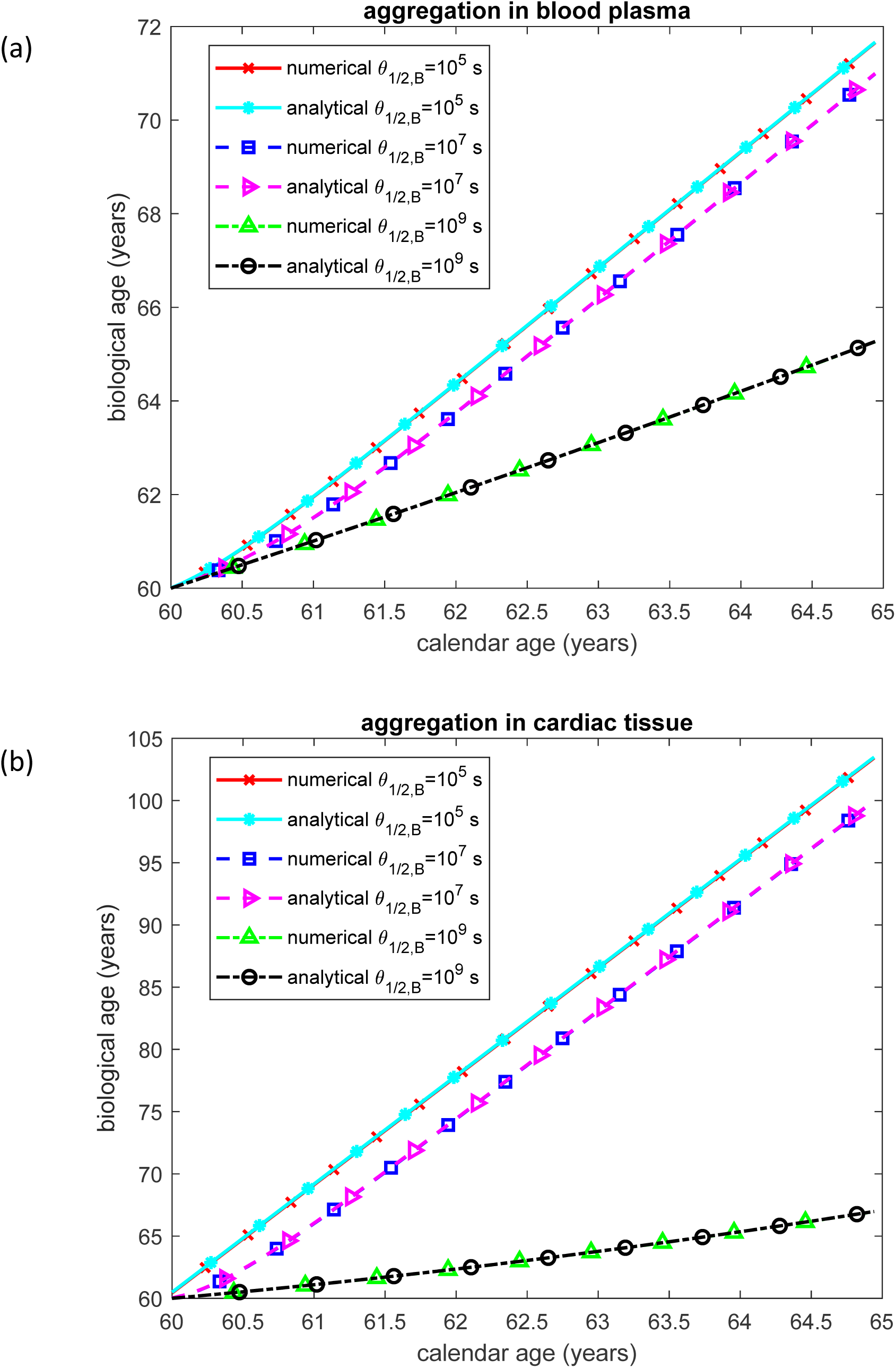
Biological age vs calendar age for various values of the half-deposition time for free TTR oligomers to become incorporated into TTR fibrils. (a) Scenario when TTR monomers misfold and assemble into protofibrillar formations within the bloodstream before depositing in the heart. (b) Scenario when TTR monomers remain in circulation until they directly attach to sites within the heart, where they form amyloid deposits *in situ*.

In the scenario where TTR aggregation occurs within cardiac tissue, and for *θ*_1/2,*B*_ = 10^7^ s, the biological age reaches 100 years after five years of disease progression—a value approaching the maximum human lifespan (Fig. 9b). In contrast, when TTR aggregates in the plasma, and for *θ*_1/2,*B*_ = 10^7^ s, biological age increases to only 71 years after the same disease duration (Fig. 9a). Given that the average survival time for patients with wild-type TTR amyloidosis is approximately five years, the model in which TTR monomers first deposit in cardiac tissue and subsequently aggregate into fibrils appears to more accurately reflect the clinical course of the disease.

The molar concentration of TTR monomers in blood plasma, [*M*]*_plasma_*, remains within a reasonable range (not exceeding 0.3 µM at steady state) when *θ*_1/2,*I*_ is varied in the scenario where TTR aggregates in the plasma (Fig. S11a), as well as when *h_M_* is varied in the scenario where TTR aggregation occurs in cardiac tissue (Fig. S11b). In both cases, [*M*]*_plasma_* gradually decreases over time and eventually stabilizes at low steady-state levels.

Assuming that TTR monomers aggregate in plasma leads to a gradual but continuous increase in total TTR concentration, ultimately reaching levels that are not observed under physiological conditions. At *θ*_1/2,*B*_ =10^7^ s, the model predicts a steady-state concentration nearing 3000 mg/dL (Fig. S12a), indicating that this aggregation pathway is unlikely to occur *in vivo*. The steady-state concentration of [*TTR*]*_total_* increases with larger values of *θ*_1/2,*I*_.

When TTR aggregation occurs in cardiac tissue, the total TTR concentration initially spikes—due to the prescribed initial condition—and then rapidly decreases to a steady-state value. As illustrated in Fig. S12b, the steady-state TTR concentration remains between 24.9 and 25.2 mg/dL, falling well within the 18–45 mg/dL reference range reported in the literature (Hood et al., 2022; Nativi-Nicolau, 2018). Increasing *h_M_* leads to a reduction in the steady-state total TTR concentration.

The molar concentration of TTR oligomers deposited into amyloid fibrils that infiltrate the cardiac tissue [*I*]*_tissue_*, increases over time, consistent with the progressive accumulation of TTR fibrils in the heart as the disease advances (Fig. S13). In the scenario where TTR aggregates in the blood plasma [*I*]*_tissue_* increases as *θ*_1/2,*I*_ decreases (Fig. S13a). This is because lower deposition time promotes more rapid deposition of TTR fibrils into cardiac tissue. In contrast, when TTR aggregates directly within the cardiac tissue, [*I*]*_tissue_* is largely insensitive to changes in *h_M_* (Fig. S13b).

A similar trend is observed for the volumetric ratio of deposited TTR fibrils to baseline myocardial volume (Fig. S14). As in Fig. 8, after 5 years of the disease this ratio reaches approximately 10% when aggregation occurs in plasma for *θ*_1/2,*I*_ = 10^7^ s, and increases to 58% when aggregation takes place in cardiac tissue for *h_M_* = 10^-6^ m s^-1^.

As in Fig. 9a, in the scenario where TTR aggregates in the plasma, the biological age reaches 71 years after 5 years of disease progression for *θ*_1/2,*I*_ = 10^7^ s (Fig. S15a). In contrast, when TTR aggregates in cardiac tissue, the biological age exceeds 100 years for *h_M_* = 10^-6^ m s^-1^ (see Fig. S15b, also compare with Fig. 9b).

The volumetric ratio of deposited TTR fibrils to baseline myocardial volume after 5 years of the disease increases with higher dissociation rates of TTR tetramers into monomers, *k_M_*, and with shorter half-deposition times for free TTR oligomers to be incorporated into fibrils, *θ*_1/2,*B*_ (Fig. 10). It is worth noting that for large values of *k_M_* the predicted volumetric ratio of deposited TTR fibrils to baseline myocardial volume can exceed one (Fig. 10b), similar to the behavior observed in Fig. 4b. Biological age after 5 years of the disease follows similar trends: it increases with an increase in *k_M_* and also increases as *θ*_1/2,*B*_ decreases (Fig. 11).

**Fig. 10.**
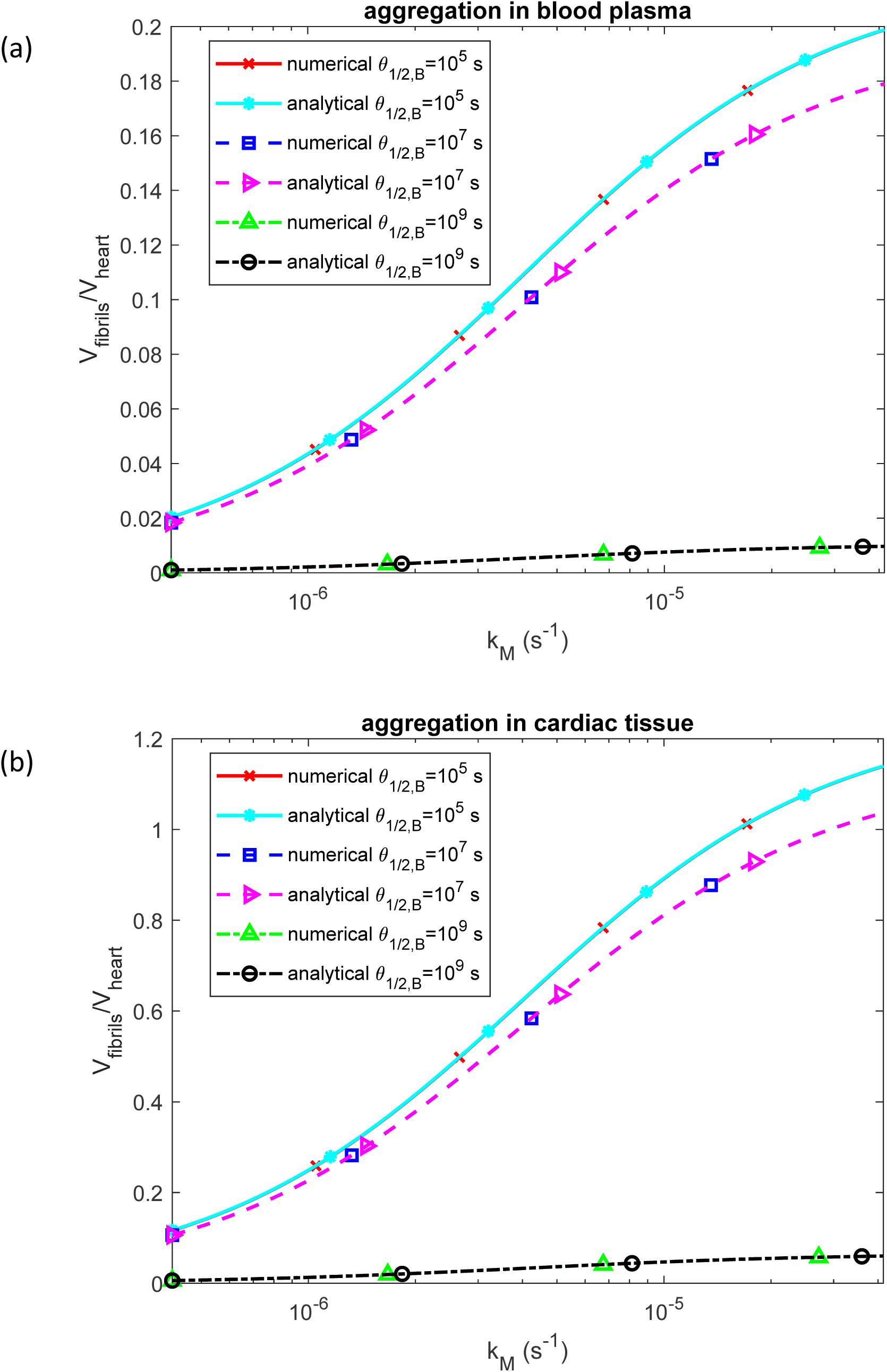
Volumetric ratio of deposited TTR fibrils to baseline myocardial volume as a function of the rate constant describing the dissociation of TTR tetramers into monomers, *k_M_*. (a) Scenario in which TTR monomers misfold and assemble into protofibrils within the bloodstream before depositing in the heart, shown for various values of *θ*_1/2,*B*_. (b) Scenario in which TTR monomers circulate until they directly bind to sites in the heart and form amyloid deposits *in situ*, shown for various values of *θ*_1/2,*B*_.

**Fig. 11.**
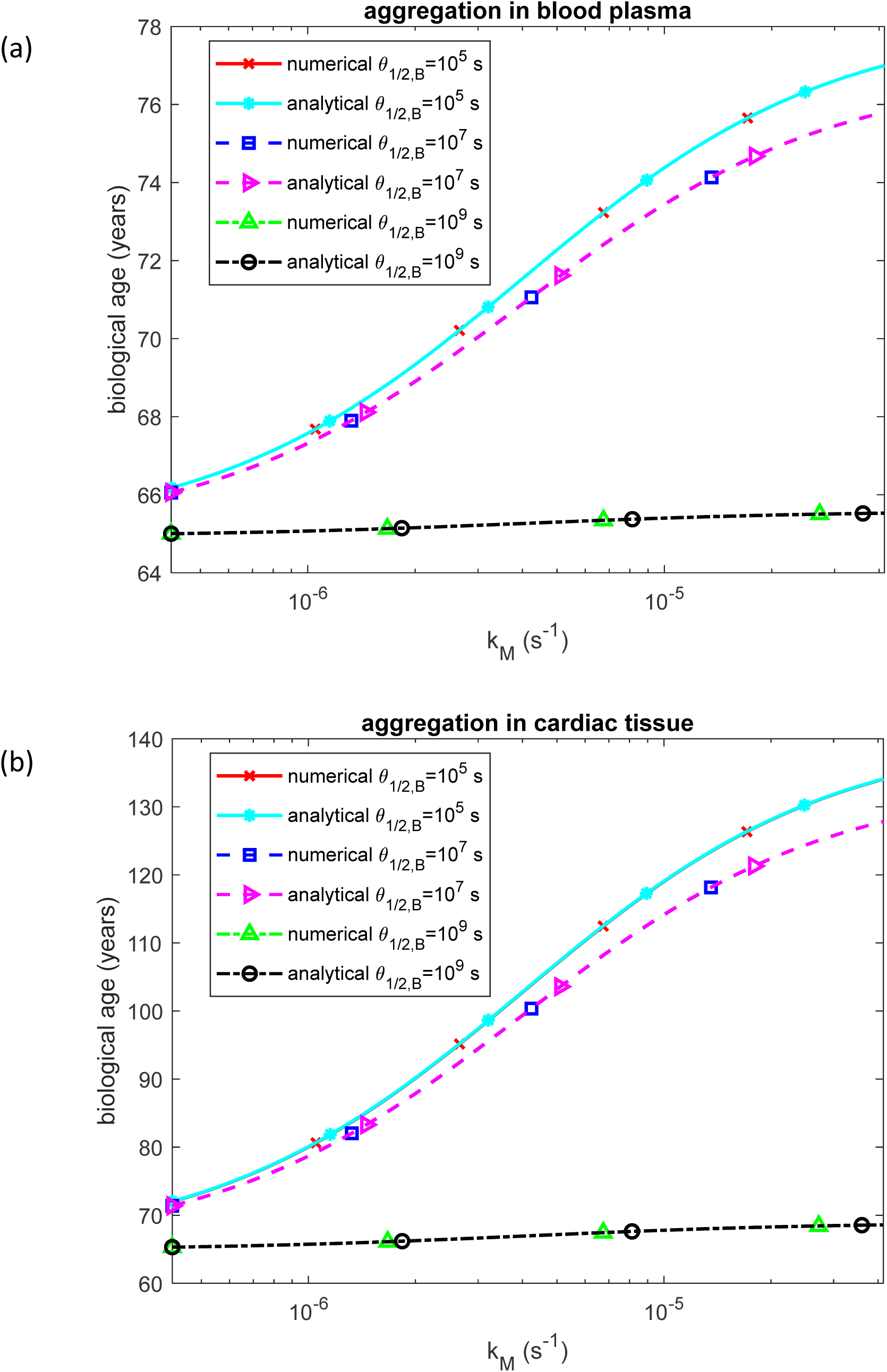
Biological age as a function of the rate constant describing the dissociation of TTR tetramers into monomers, *k_M_*. (a) Scenario in which TTR monomers misfold and assemble into protofibrils within the bloodstream before depositing in the heart, shown for various values of *θ*_1/2,*B*_. (b) Scenario in which TTR monomers circulate until they directly bind to sites in the heart and form amyloid deposits *in situ*, shown for various values of *θ*_1/2,*B*_.

The volumetric ratio of deposited TTR fibrils to baseline myocardial volume after five years of disease progression decreases as the half-deposition time for free TTR oligomers to be incorporated into fibrils, *θ*_1/2,*B*_, increases (Fig. 12). A longer *θ*_1/2,*B*_ indicates a slower rate of oligomer incorporation into fibrils, resulting in reduced fibril accumulation. Likewise, an increase in the half-deposition time for TTR-derived fibrils circulating in the blood plasma to deposit into cardiac tissue, *θ*_1/2,*I*_, also leads to a lower fibril burden in the heart, as fewer circulating fibrils are deposited (Fig. 12a). In contrast, the mass transfer coefficient governing the deposition rate of TTR monomers into the endocardium, *h_M_*, shows no significant effect on fibril accumulation (Fig. 12b).

**Fig. 12.**
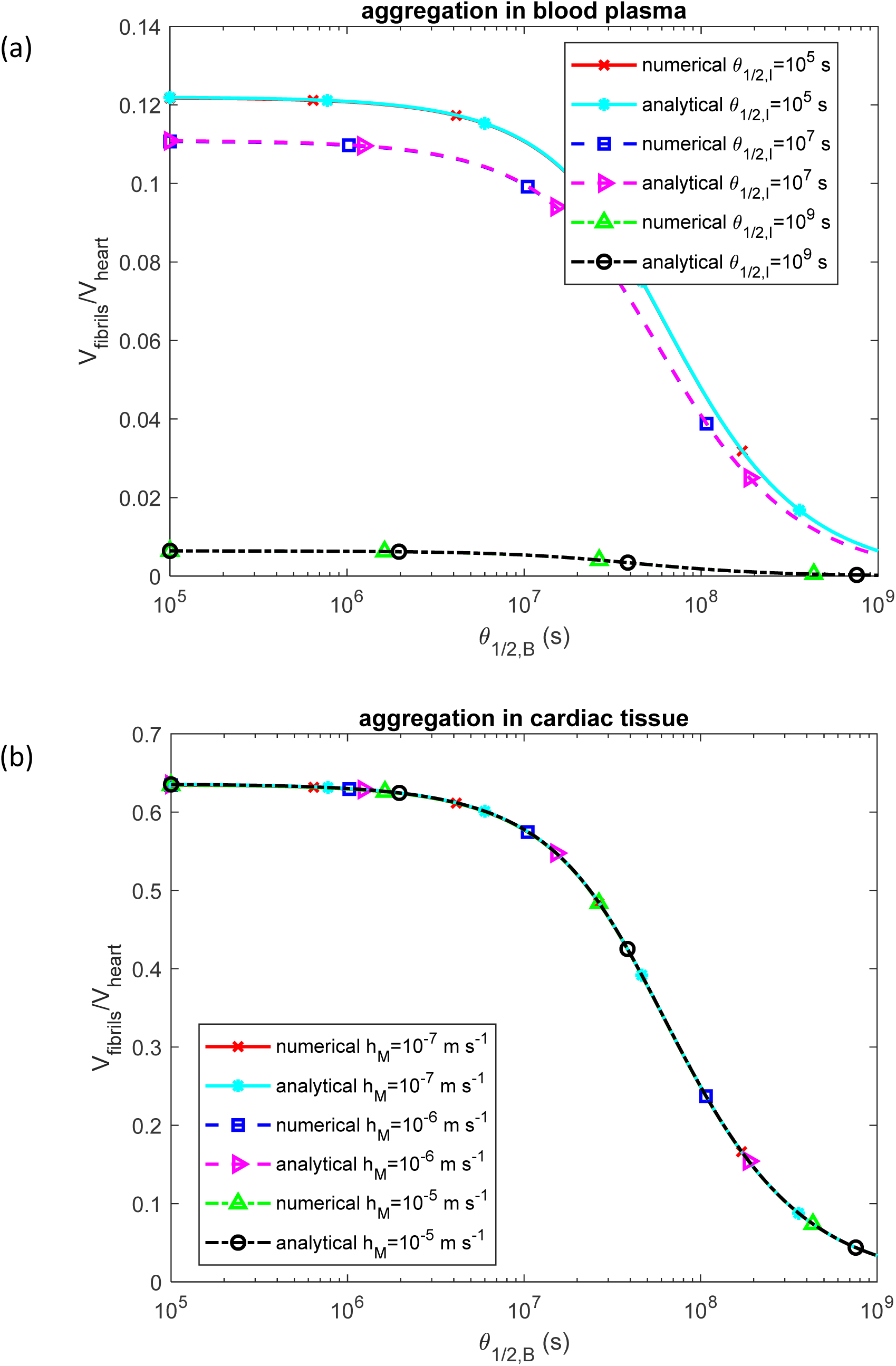
Volumetric ratio of deposited TTR fibrils to baseline myocardial volume as a function of the half-deposition time required for free TTR oligomers to become incorporated into fibrils, *θ*_1/2,*B*_. (a) Scenario in which TTR monomers misfold and assemble into protofibrils within the bloodstream before depositing in the heart, shown for various values of *θ*_1/2,*I*_. (b) Scenario in which TTR monomers circulate until they directly bind to sites in the heart and form amyloid deposits *in situ*, shown for various values of *h_M_*.

Similarly, biological age after five years of the disease decreases with increasing *θ*_1/2,*B*_ (Fig. 13) and also decreases with increasing fibril deposition half-time from plasma to cardiac tissue, *θ*_1/2,*I*_ (Fig. 13a). In contrast, variations in the mass transfer coefficient for monomer deposition, *h_M_*, have no noticeable effect on biological age (Fig. 13b).

**Fig. 13.**
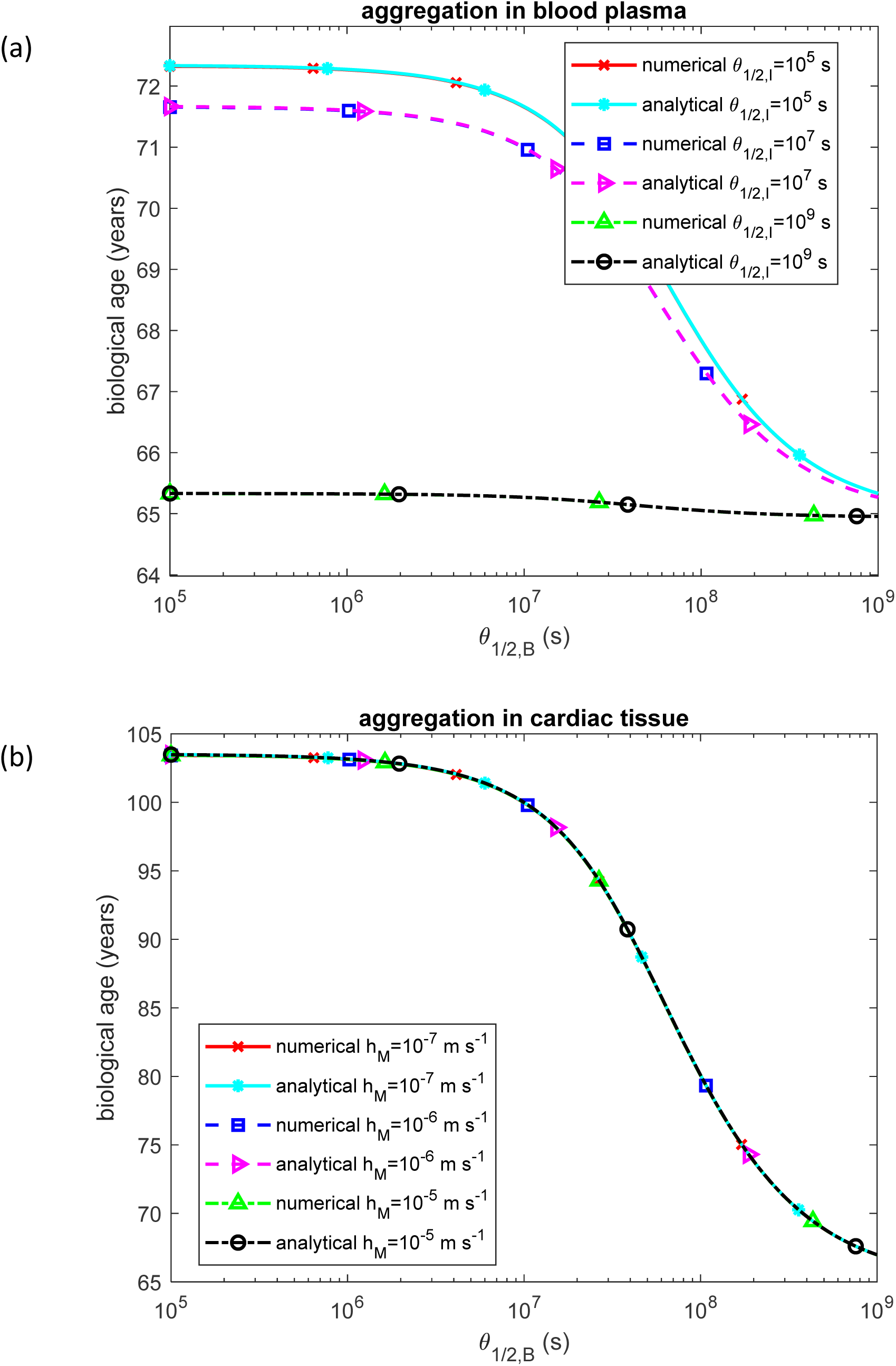
Biological age as a function of the half-deposition time required for free TTR oligomers to become incorporated into fibrils, *θ*_1/2,*B*_. (a) Scenario in which TTR monomers misfold and assemble into protofibrils within the bloodstream before depositing in the heart, shown for various values of *θ*_1/2,*I*_. (b) Scenario in which TTR monomers circulate until they directly bind to sites in the heart and form amyloid deposits *in situ*, shown for various values of *h_M_*.

The volumetric ratio of deposited TTR fibrils to baseline myocardial volume after 5 years of the disease decreases as the half-deposition time for TTR-derived fibrils circulating in the blood plasma to deposit into cardiac tissue *θ*_1/2,*I*_ increases (Fig. S16a), and remains unaffected by changes in the mass transfer coefficient characterizing the rate of TTR monomer deposition into the endocardium, *h_M_* (Fig. S16b). Overall, an increase in *θ*_1/2,*B*_ leads to a reduction in the heart volume occupied by TTR amyloid fibrils (Fig. S16). This is consistent with the findings presented in Fig. 12.

Similarly, biological age after 5 years of the disease decreases as *θ*_1/2,*I*_ increases (Fig. S17a) and remains unaffected by changes in *h_M_* (Fig. S17b). Overall, an increase in *θ*_1/2,*B*_ results in a reduction in biological age. These observations align with the trends shown in Fig. 13.

The sensitivity of the fraction of heart volume occupied by TTR amyloid fibrils to the dissociation rate constant of TTR tetramers into monomers, *k_M_*, is positive, indicating that a higher dissociation rate leads to increased fibril accumulation in the heart. This is consistent with the trend observed in Fig. 11. The sensitivity declines rapidly over time before stabilizing at a positive steady-state value (Fig. 14).

**Fig. 14.**
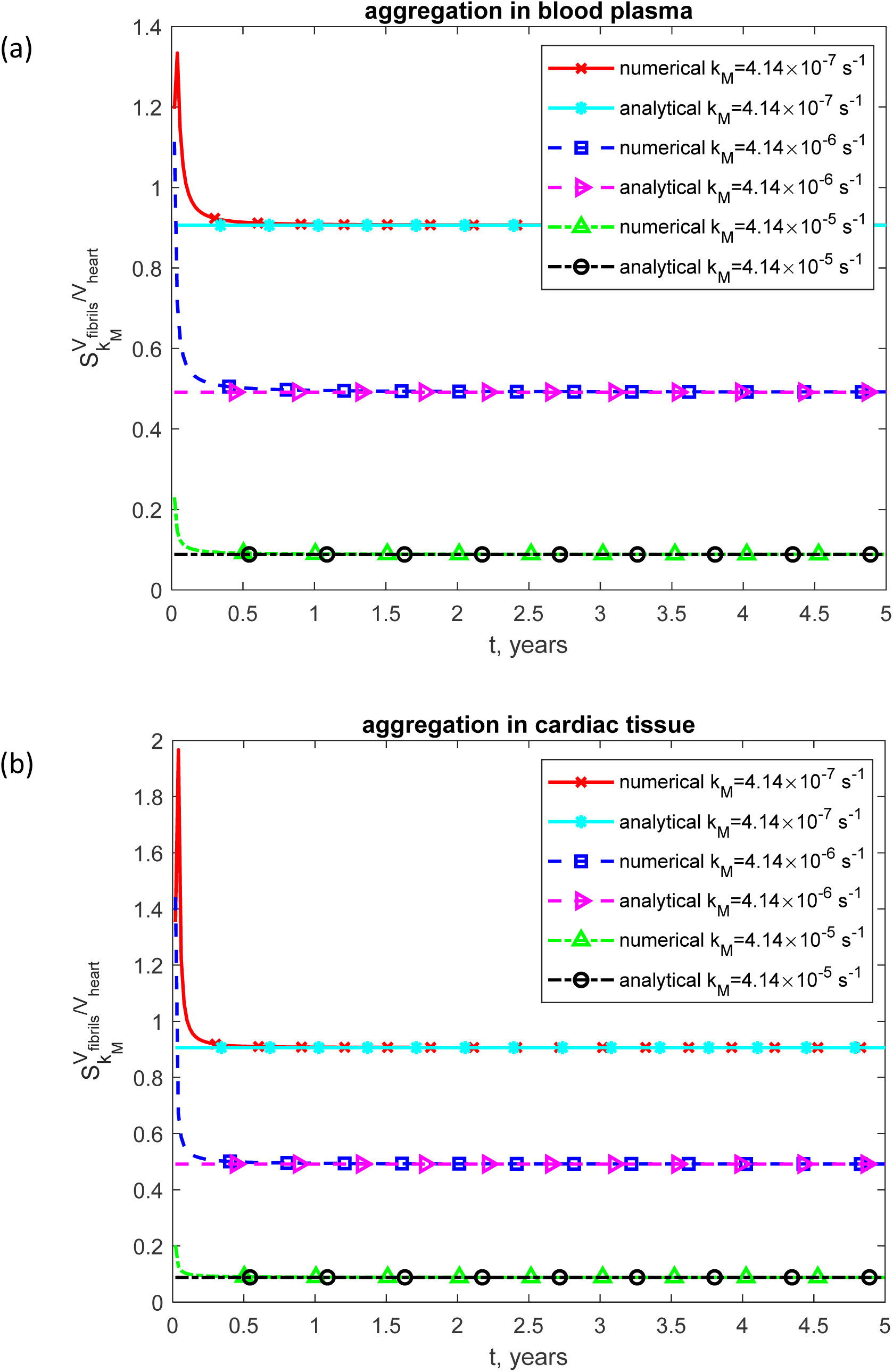
Volumetric ratio of deposited TTR fibrils to baseline myocardial volume occupied by TTR amyloid fibrils to the rate constant describing the dissociation of TTR tetramers into monomers, *k_M_*, shown as a function of time. (a) Scenario in which TTR monomers misfold and assemble into protofibrillar structures within the bloodstream before depositing in the heart. (b) Scenario in which TTR monomers remain in circulation until they directly bind to sites within the heart, where they form amyloid deposits *in situ*.

The sensitivity of the fraction of heart volume occupied by TTR amyloid fibrils to the half-deposition time required for free TTR oligomers to be incorporated into fibrils, *θ*_1/2,*B*_, is negative (Fig. 15). This indicates that as *θ*_1/2,*B*_ increases, the volume occupied by TTR fibrils decreases, consistent with the trend observed in Fig. 12. Additionally, increasing *θ*_1/2,*B*_ reduces the magnitude of the sensitivity (Fig. 15).

**Fig. 15.**
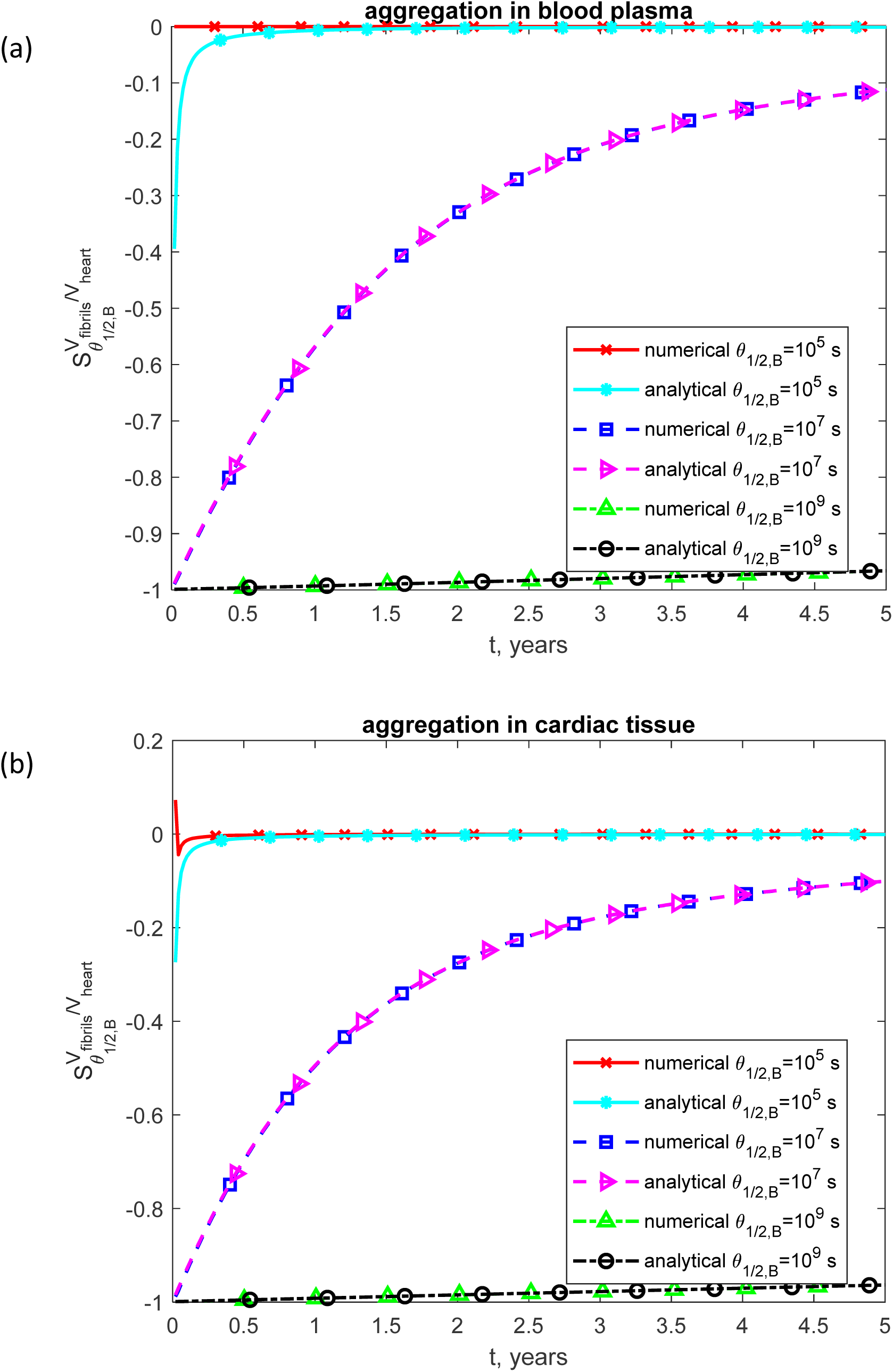
Dimensionless sensitivity of the fraction of heart volume occupied by TTR amyloid fibrils to the half-deposition time required for free TTR oligomers to be incorporated into fibrils, *θ*_1/2,*B*_, shown as a function of time. (a) Scenario in which TTR monomers misfold and assemble into protofibrillar structures within the bloodstream before depositing in the heart. (b) Scenario in which TTR monomers remain in circulation until they directly bind to sites within the heart, where they form amyloid deposits *in situ*.

The sensitivity of the fraction of heart volume occupied by TTR amyloid fibrils to the half-deposition time for TTR-derived fibrils circulating in the blood plasma to deposit into cardiac tissue, *θ*_1/2,*I*_, is negative (Fig. S18a), indicating that an increase in *θ*_1/2,*I*_ leads to a decrease in fibril accumulation. This is consistent with the trend shown in Fig. S16a. In contrast, the volume occupied by TTR fibrils is largely insensitive to *h_M_*, except at early time points (Fig. S18b). The analytical solution does not depend on *h_M_* (see Eqs. (54) and (55)). Therefore, its sensitivity to *h_M_* is zero, as shown in Fig. S18b.

In the simulations described thus far in this paper, TTR fibrils were modeled as inert sinks for oligomeric species, assuming they lack the catalytic capacity to facilitate monomer-to-oligomer conversion. This baseline scenario is characterized by *k*_3_ = 0. Next a scenario where both oligomers and fibrils facilitate monomer-to-oligomer conversion via surface-mediated catalysis (secondary nucleation) is simulated. An increase in *k*_3_ results in a reduction of monomer concentrations within the blood plasma (for the scenario when fibril assembly occurs in the plasma, Fig. S19a) and in the cardiac tissue (for the scenario when fibril assembly occurs in the cardiac tissue, Fig. S20). This reduction is attributed to the fact that when *k*_3_ > 0, monomer conversion is driven by both autocatalysis (via existing oligomers) and surface-mediated catalysis (via the fibrillar lattice). Furthermore, for *k*_3_ > 0 plasma monomer concentrations are attenuated even when fibril assembly is localized to the cardiac tissue (Fig. 19b), a phenomenon attributed to the enhanced flux of monomers from the plasma into the myocardium (Eq. (33)).

However, due to the high baseline concentration of oligomers, the additional conversion occurring on the fibril surface does not significantly elevate total oligomer levels (Fig. S21). Similarly, there is no significant increase in the concentration of oligomers incorporated into the fibrillar mass (Figs. S22 and S23a,b), and the total volume of cardiac amyloid deposition remains largely unaffected by the *k*_3_ value (Fig. 24a,b). These observations extend to the predicted biological age of the heart (Fig. S25a,b). These findings suggest that the rate of myocardial fibril deposition is primarily dictated by upstream supply kinetics—specifically the rate-limiting dissociation of TTR tetramers into unstable monomers—rather than the downstream kinetics of oligomerization or fibrillar growth. This conclusion is further evidenced by Fig. 16, which demonstrates that the sensitivity of cardiac fibril volume to *k*_3_ is negligible.

**Fig. 16.**
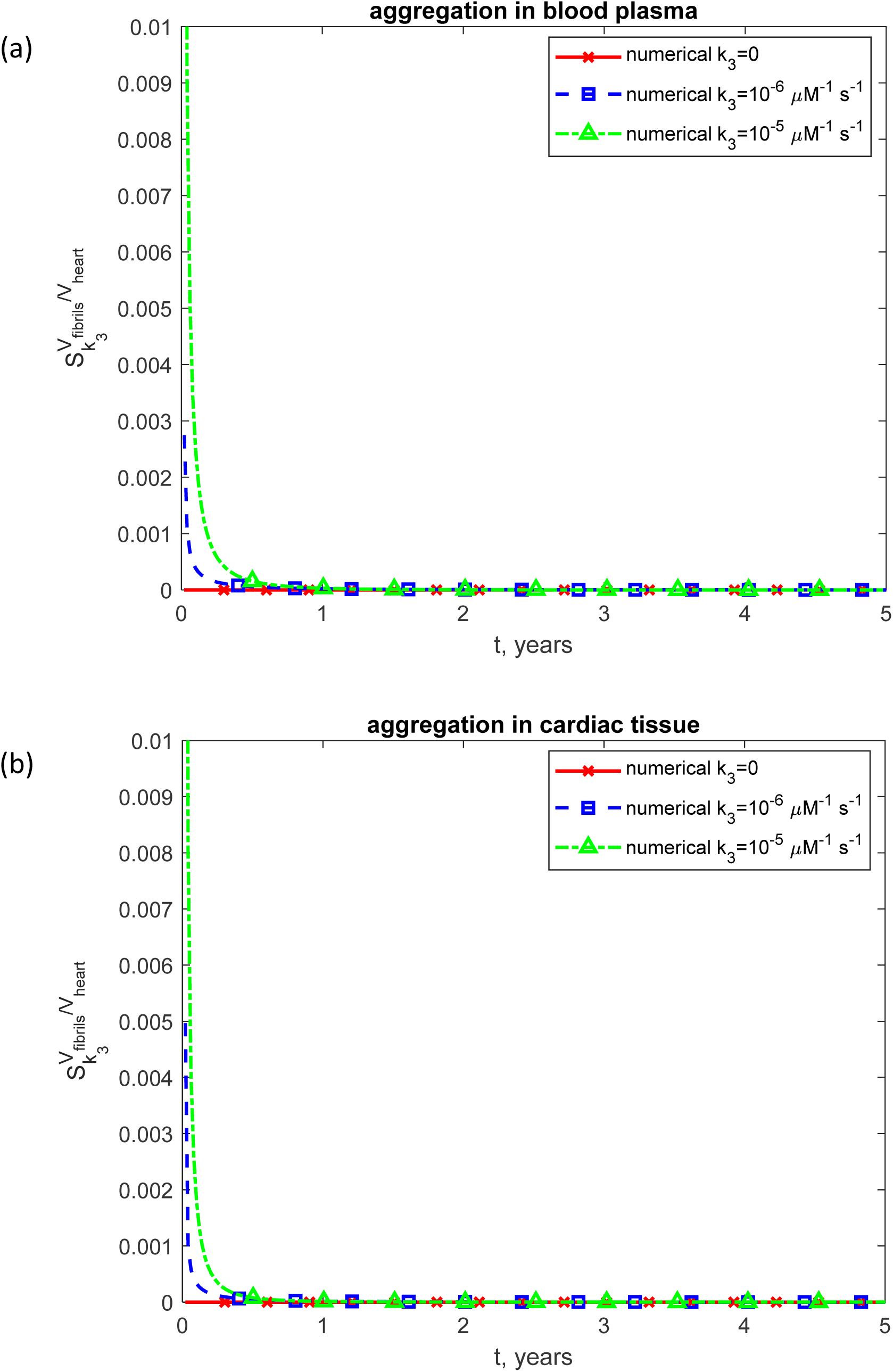
Dimensionless sensitivity of the fraction of heart volume occupied by TTR amyloid fibrils to the kinetic rate constant for surface-catalyzed oligomerization, *k*_3_, shown as a function of time. (a) Scenario in which TTR monomers misfold and assemble into protofibrillar structures within the bloodstream before depositing in the heart. (b) Scenario in which TTR monomers remain in circulation until they directly bind to sites within the heart, where they form amyloid deposits *in situ*.

## 4. Discussion, limitations of the model, and future directions

This study presents a mathematical model of TTR aggregation dynamics, capturing key features of amyloid deposition in TTR amyloidosis. The obtained analytical solutions closely match numerical results for all quantities except sensitivity coefficients, which diverge at early times due to their strong dependence on solution accuracy; however, at later times, the analytically predicted sensitivity coefficients align very well with the numerical results.

The model simulates two aggregation scenarios, suggested in (Criddle et al., 2024): (1) TTR fibril formation in the bloodstream and (2) TTR aggregation within cardiac tissue. Results demonstrate that fibril accumulation in cardiac tissue leads to physiologically realistic TTR concentrations and disease progression consistent with clinical observations. In contrast, the scenario where fibril formation occurs in plasma predicts unrealistically high total TTR concentrations, which may exceed 3000 mg/dL. Such high concentrations are not observed in patients.

The obtained results corroborate the established view that tetramer dissociation is the rate-limiting step in amyloidogenesis (Esperante et al., 2021; Bezerra et al., 2020). Furthermore, the simulations suggest that the cardiac tissue aggregation pathway provides a more plausible mechanistic basis for the clinical progression of wild-type ATTR cardiomyopathy. Specifically, this pathway predicts levels of fibril accumulation that are consistent with the high cardiac amyloid burdens clinically associated with increased mortality (Mehta et al., 2019), thereby aligning the model’s kinetic outputs with observed patient outcomes.

The model also captures the therapeutic mechanism of TTR stabilizers such as tafamidis. By reducing the tetramer dissociation rate, tafamidis effectively raises the concentration of native TTR tetramers while limiting the formation of non-native aggregation-prone species (Monteiro et al., 2023; Nelson et al., 2021). Simulations reported in this paper agree with this experimental observation, showing that a decrease in the tetramer dissociation constant results in higher tetramer concentrations and reduced fibril burden. The model predicts that the molar concentration of TTR tetramers remains unaffected by changes in the fibril deposition kinetics of oligomers, reinforcing the idea that tetramer dissociation is upstream and largely independent of later aggregation stages. Moreover, the model highlights the role of autocatalytic oligomer formation: longer oligomer half-lives result in higher oligomer concentrations, which in turn accelerate the conversion of monomers into oligomers, particularly when aggregation occurs in plasma.

Our results demonstrate a pattern of TTR deposition that aligns with clinical observations of progressive tissue dysfunction and age-related decline (Almeida et al., 2024). While the model primarily simulates the kinetics of fibril accumulation over time, the predicted amyloid burden reaches thresholds that carry significant prognostic weight; specifically, (Mehta et al., 2019) have linked higher levels of cardiac amyloidosis burden to increased mortality rates. The model underscores the therapeutic importance of stabilizing tetramers early in the aggregation cascade to mitigate long-term myocardial burden.

To quantify systemic effect, the model defines biological age as a linear function of the volumetric ratio of deposited TTR fibrils to baseline myocardial volume. Biological age can be used as a surrogate marker to evaluate how different aggregation pathways impact disease progression. The scaling coefficient *c* was calibrated to ensure the tissue-deposition trajectory reaches a terminal threshold of 100 years exactly five years after the onset of myocardial fibril deposition. In contrast, using the same kinetic framework for the plasma aggregation based model yields a significantly more modest increase in biological age (reaching only 71 years), failing to replicate the rapid decline observed in patients. These results suggest that the kinetics of tissue-localized aggregation, rather than plasma-derived infiltration, are more consistent with the accelerated pathophysiology of the disease.

Simulation results also indicate that the rate-limiting step in the amyloidogenic pathway is the dissociation of the native TTR tetramer into its constituent monomers. Consequently, it is this upstream kinetic event—rather than downstream oligomer production and fibrillar assembly—that primarily dictates the total myocardial amyloid burden. From a clinical perspective, these results highlight the therapeutic necessity of reducing the systemic concentration of amyloidogenic monomers. This observation aligns with the pharmacological profile of TTR kinetic stabilizers, such as tafamidis; by inhibiting the initial dissociation of tetramers into aggregation-prone monomers, these agents target the primary bottleneck in the amyloidogenic cascade, representing an effective strategy for mitigating cardiac amyloid deposition.

The proposed model is a minimal representation of TTR amyloidosis, built on conservation principles for different TTR assembly states, including monomers, oligomers, and fibrils. It simplifies several biophysical processes—for example, TTR aggregation is modeled using the F-W framework, which does not distinguish between different oligomer sizes—highlighting the need for comparison with experimental data for further model development. The model does not resolve the size distribution of individual fibrils; instead, it focuses on the total volume of deposited fibrils.

Another limitation of the current model is the assumption of a uniform spatial distribution for all TTR species within the cardiac compartment. While this simplification is appropriate for the circulating blood plasma, myocardial amyloidosis is characterized by significant spatial heterogeneity. Clinical and histological observations often reveal non-uniform, patchy deposition patterns, where amyloid fibrils accumulate preferentially within the extracellular matrix of specific regions, such as the subendocardium (Cianci et al., 2024).

Because the model employs a compartmental mass-balance framework, it calculates a volume-averaged amyloid density across the entire heart volume. Consequently, it may underestimate local pathological effects, such as localized tissue stiffening or high-density catalytic zones where secondary nucleation catalyzed by fibrils (characterized by *k*_3_) may be accelerated. Future modeling efforts incorporating spatially resolved techniques, such as reaction-diffusion equations or finite element analysis, would be required to capture the anisotropic nature of TTR deposition and its localized impact on cardiac mechanics.

While defining biological age as a linear function of myocardial TTR deposition does not account for the non-linear physiological decline or homeostatic compensatory mechanisms typical of the aging myocardium, it serves as a robust comparative surrogate biomarker for assessing cumulative amyloid burden across various modeling scenarios.

Another limitation of the developed model is the very limited data regarding the duration of the subclinical phase in ATTR. Future research should consider a range for the temporal gap between the onset of fibril deposition and clinical detection and investigate the sensitivity of model predictions to this parameter.

## Data accessibility

This article has no additional data.

## Authors’ contributions

AVK is the sole author of this paper.

## Abbreviations

ATTR: transthyretin amyloidosis
F-W: Finke-Watzky
IDP: intrinsically disordered protein
TTR: transthyretin

## Supplemental Materials

### S1. Numerical solution

The systems of differential equations were solved numerically using MATLAB’s ODE45 solver (R2024a, MathWorks, Natick, MA, USA). For scenario 1, Eqs. (2), (9)–(11), and (14) with initial conditions (3) and (15) were used, while scenario 2 employed Eqs. (2), (31), and (33)–(35) with initial conditions (3) and (36). ODE45 solver, which uses an adaptive step-size Runge-Kutta method, was selected for its balance of accuracy and computational efficiency. To achieve high numerical precision, both the relative and absolute tolerances were set to 1e-10.

### S2. Supplementary figures

**Fig. S1.**
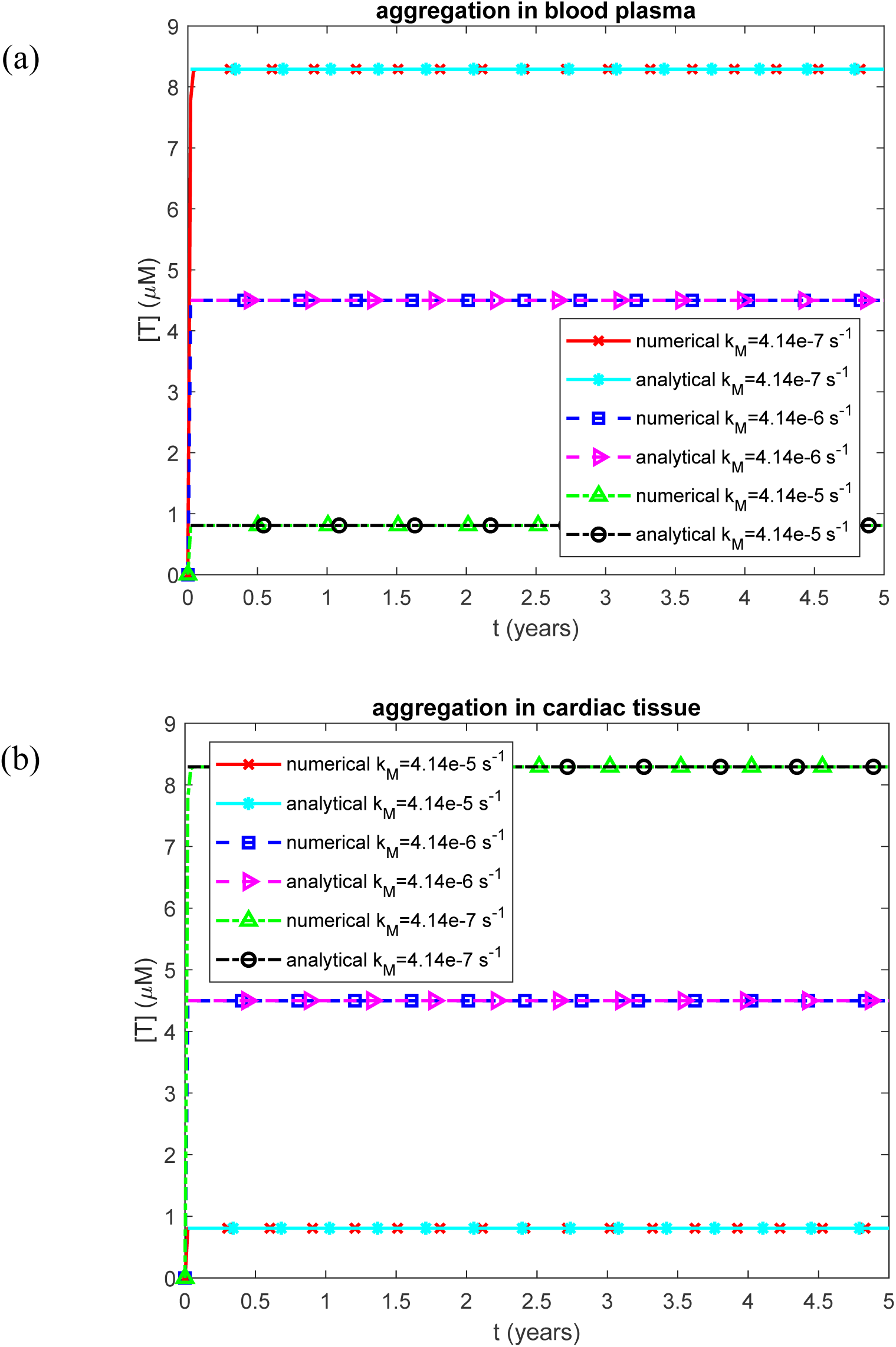
Molar concentration of TTR tetramers, representing the native form of TTR, in the blood plasma vs time for various values of *k_M_*. (a) Scenario when TTR monomers misfold and assemble into protofibrillar formations within the bloodstream before depositing in the heart. (b) Scenario when TTR monomers remain in circulation until they directly attach to sites within the heart, where they form amyloid deposits *in situ*.

**Fig. S2.**
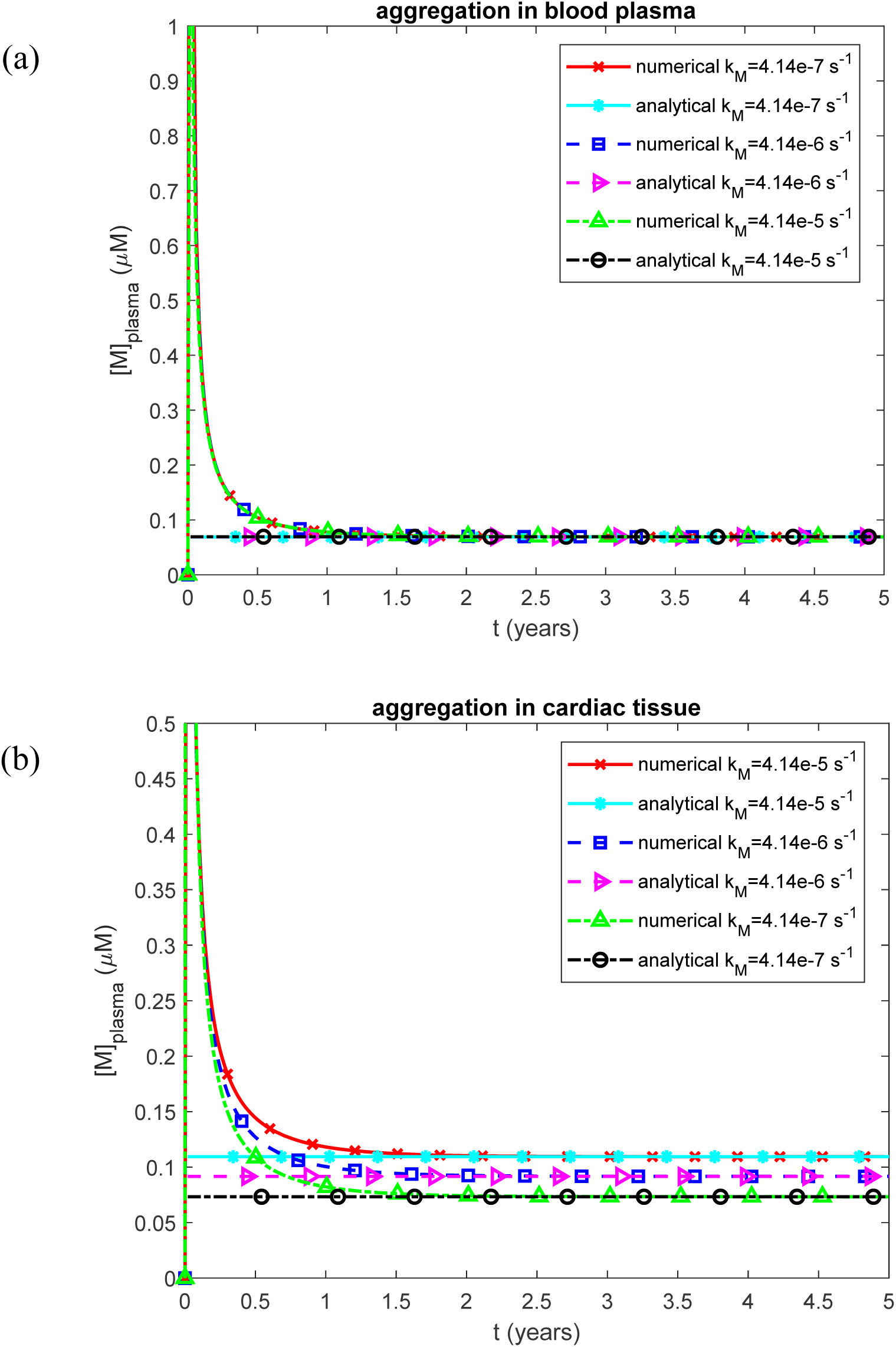
Molar concentration of TTR monomers in the blood plasma vs time for various values of *k_M_*. (a) Scenario when TTR monomers misfold and assemble into protofibrillar formations within the bloodstream before depositing in the heart. (b) Scenario when TTR monomers remain in circulation until they directly attach to sites within the heart, where they form amyloid deposits *in situ*.

**Fig. S3.**
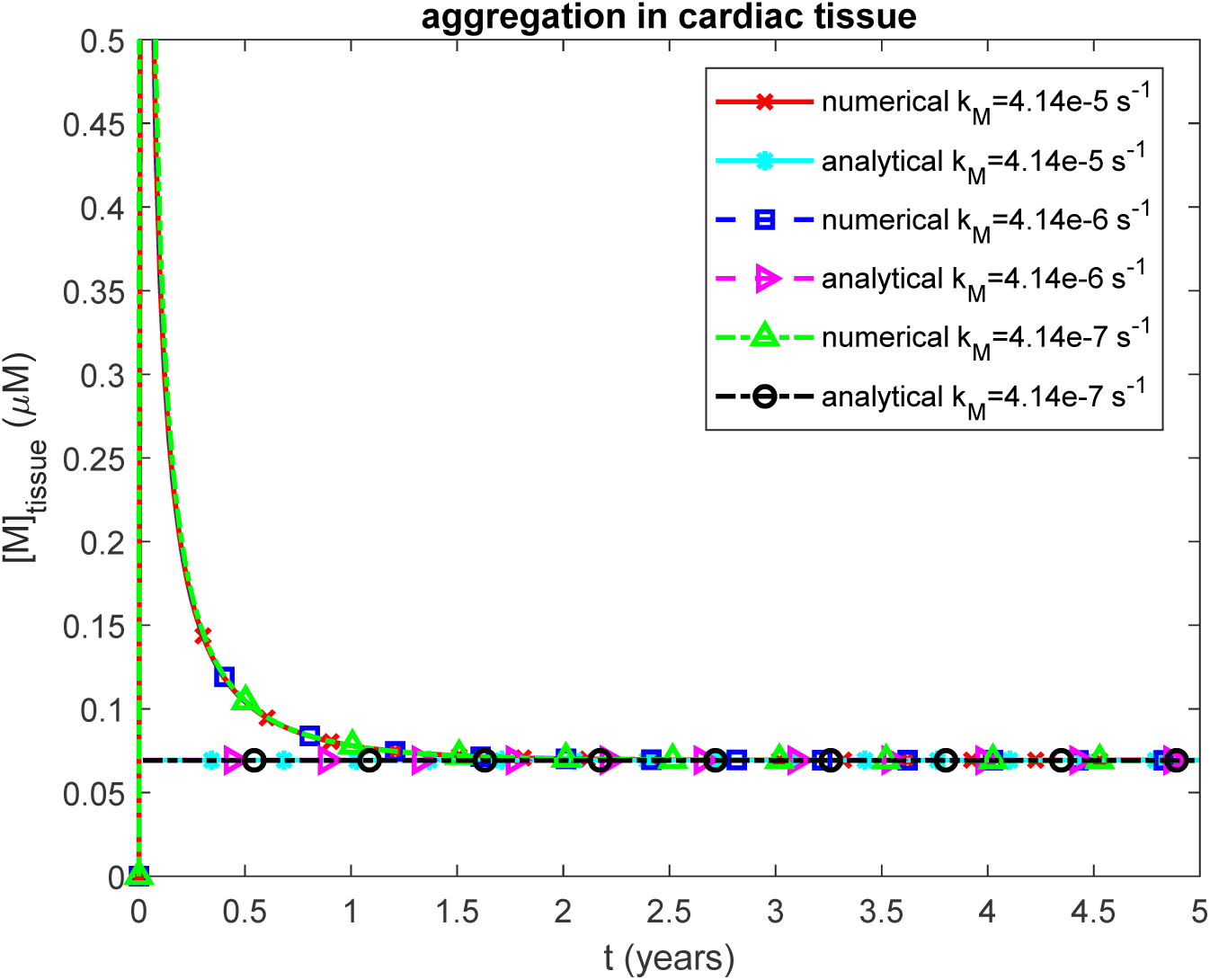
Molar concentration of TTR monomers in the cardiac tissue vs time for various values of *k_M_*. Scenario when TTR monomers remain in circulation until they directly attach to sites within the heart, where they form amyloid deposits *in situ*.

**Fig. S4.**
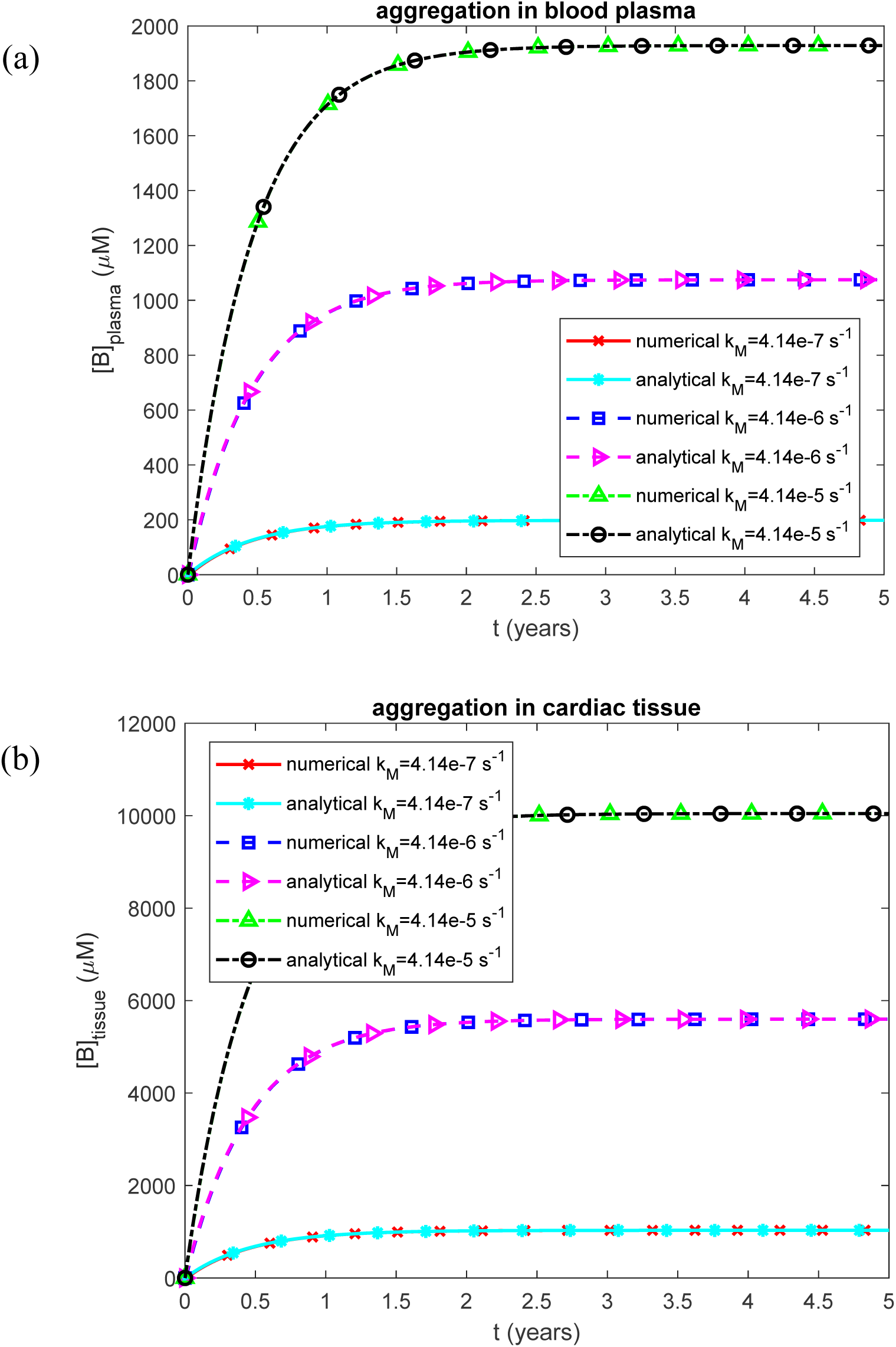
(a) Molar concentration of free TTR oligomers in the blood plasma vs time for various values of *k_M_*. Scenario when TTR accumulates in blood plasma. (b) Molar concentration of free TTR oligomers in the cardiac tissue vs time for various values of *k_M_*. Scenario when TTR accumulates in cardiac tissue.

**Fig. S5.**
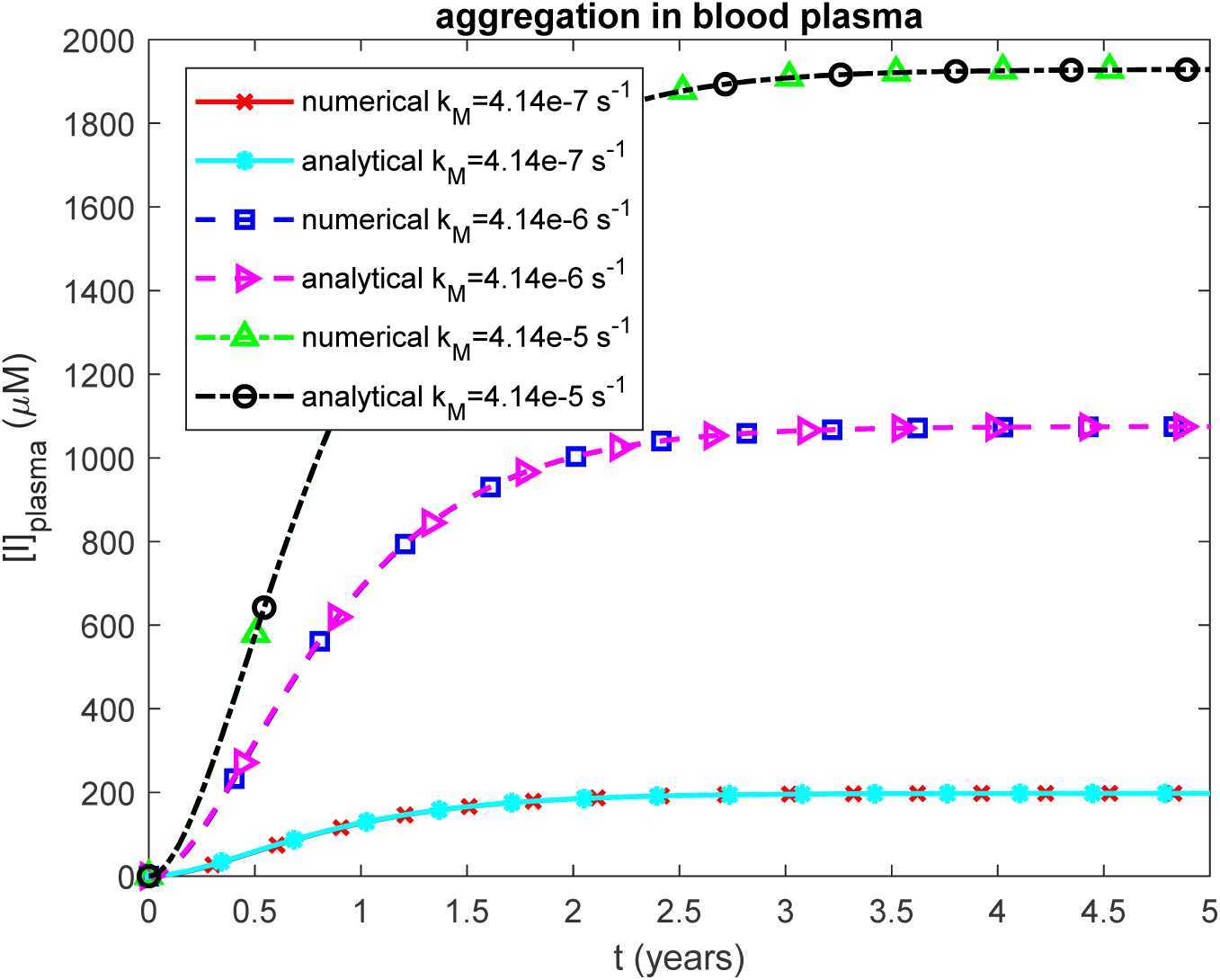
Molar concentration of TTR oligomers deposited into amyloid protofibrils circulating in the blood plasma vs time for various values of *k_M_*. Scenario when TTR monomers misfold and assemble into protofibrillar formations within the bloodstream before depositing in the heart.

**Fig. S6.**
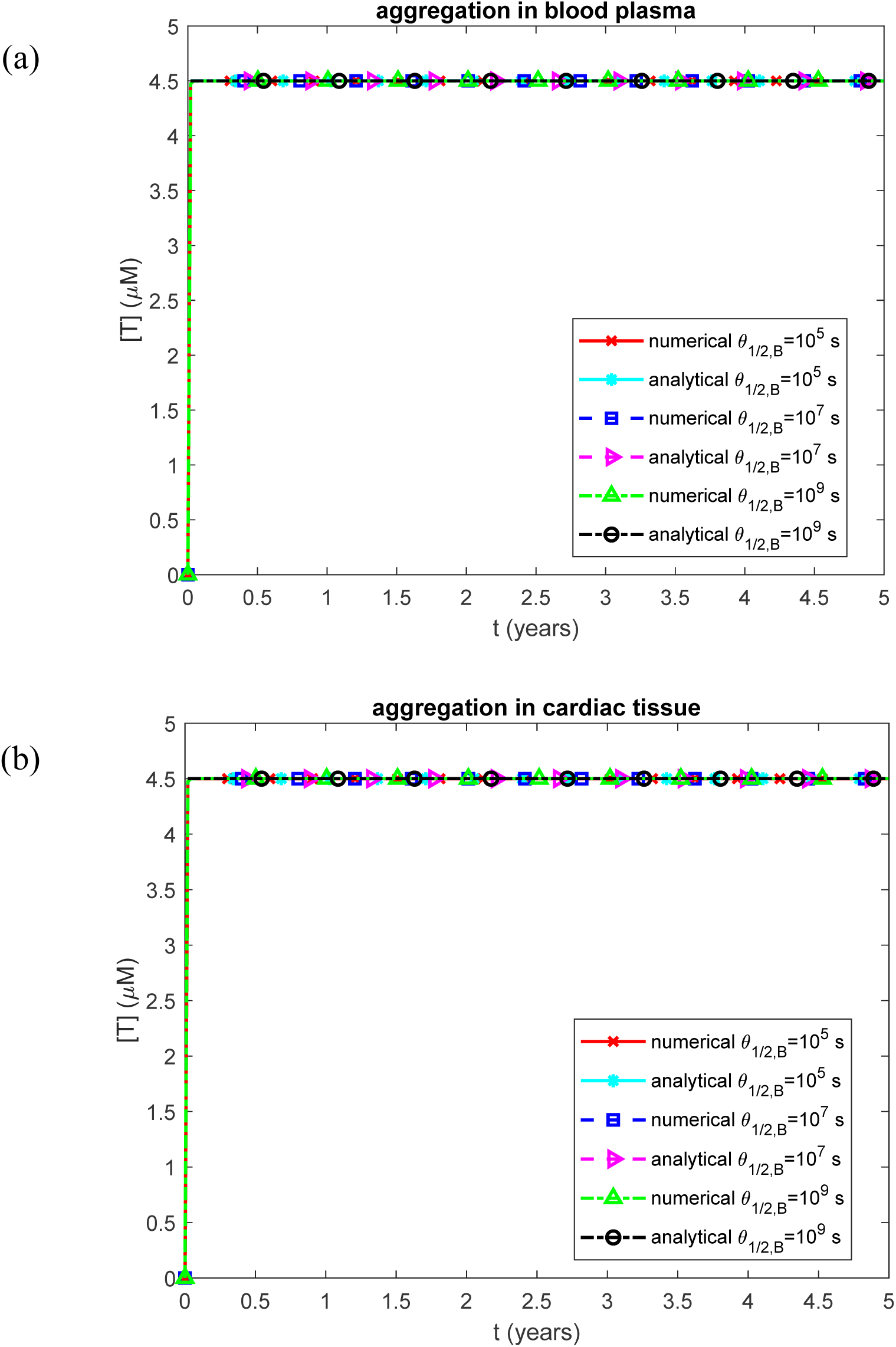
Molar concentration of TTR tetramers, representing the native form of TTR, in the blood plasma vs time for various values of *θ*_1/2,*B*_. (a) Scenario when TTR monomers misfold and assemble into protofibrillar formations within the bloodstream before depositing in the heart. (b) Scenario when TTR monomers remain in circulation until they directly attach to sites within the heart, where they form amyloid deposits *in situ*.

**Fig. S7.**
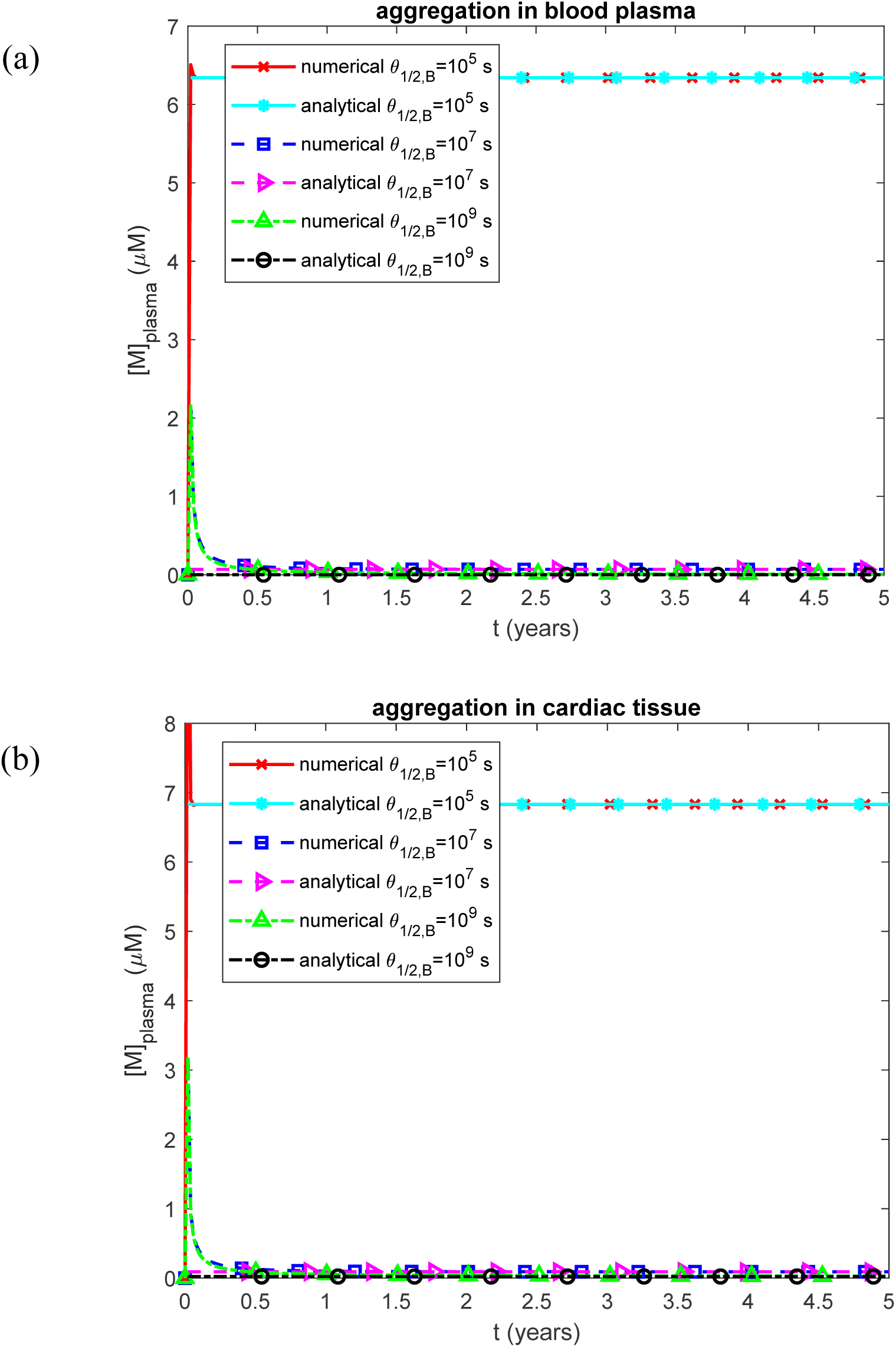
Molar concentration of TTR monomers in the blood plasma vs time for various values of *θ*_1/2,*B*_. (a) Scenario when TTR monomers misfold and assemble into protofibrillar formations within the bloodstream before depositing in the heart. (b) Scenario when TTR monomers remain in circulation until they directly attach to sites within the heart, where they form amyloid deposits *in situ*.

**Fig. S8.**
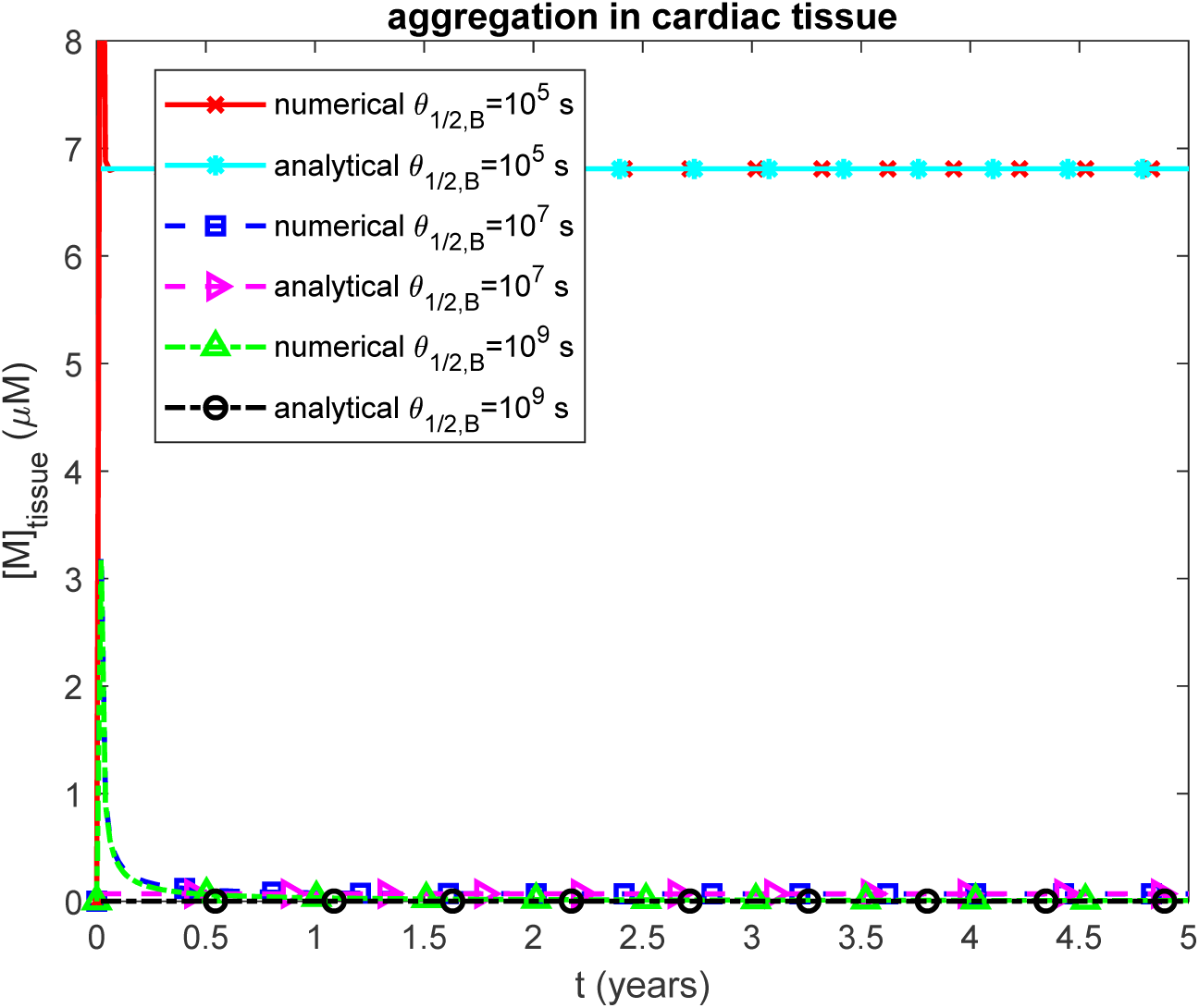
Molar concentration of TTR monomers in the cardiac tissue vs time for various values of *θ*_1/2,*B*_. Scenario when TTR monomers remain in circulation until they directly attach to sites within the heart, where they form amyloid deposits *in situ*.

**Fig. S9.**
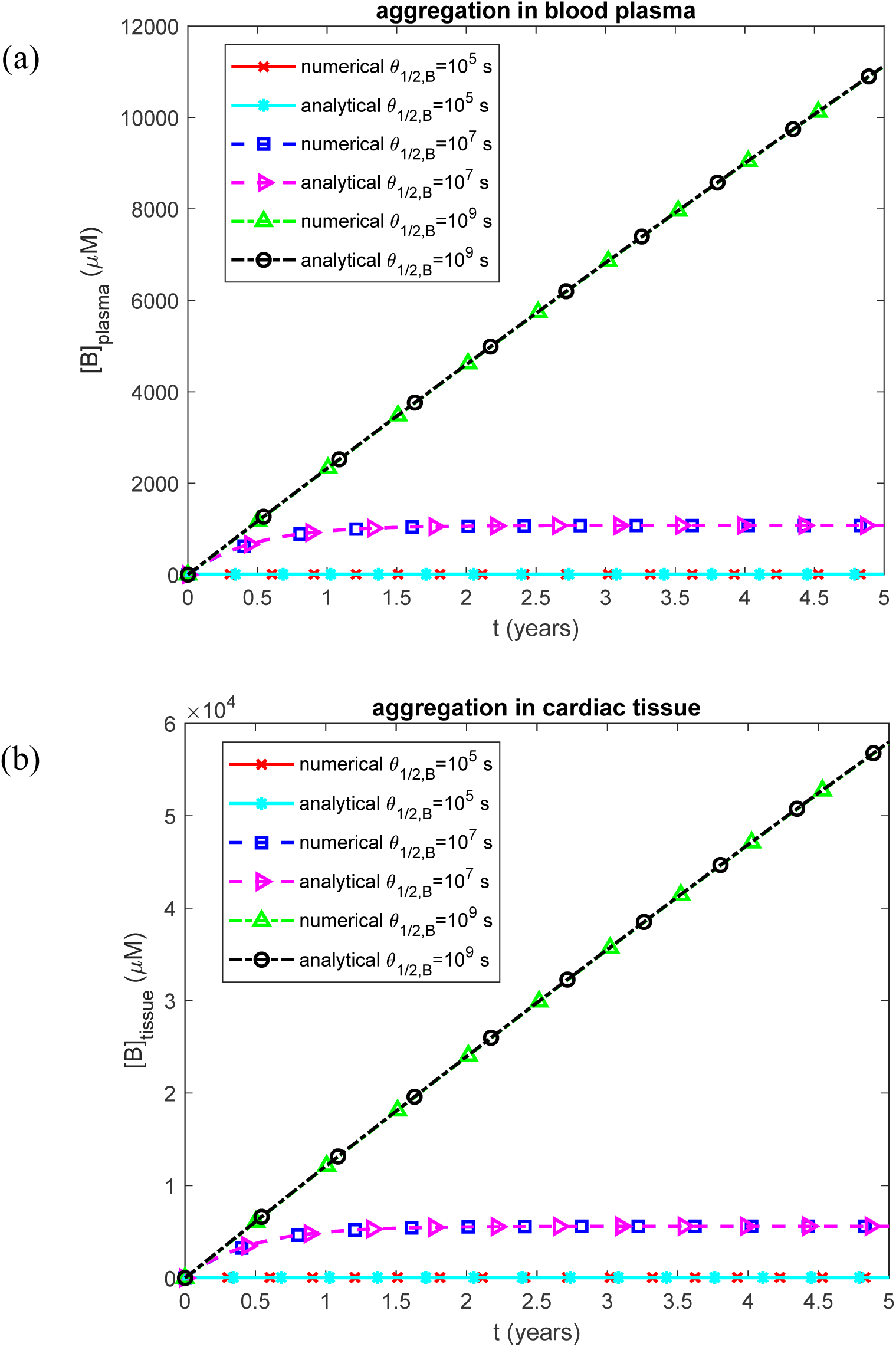
(a) Molar concentration of free TTR oligomers in the blood plasma vs time for various values of *θ*_1/2,*B*_. Scenario when TTR accumulates in blood plasma. (b) Molar concentration of free TTR oligomers in cardiac tissue. Scenario when TTR accumulates in cardiac tissue.

**Fig. S10.**
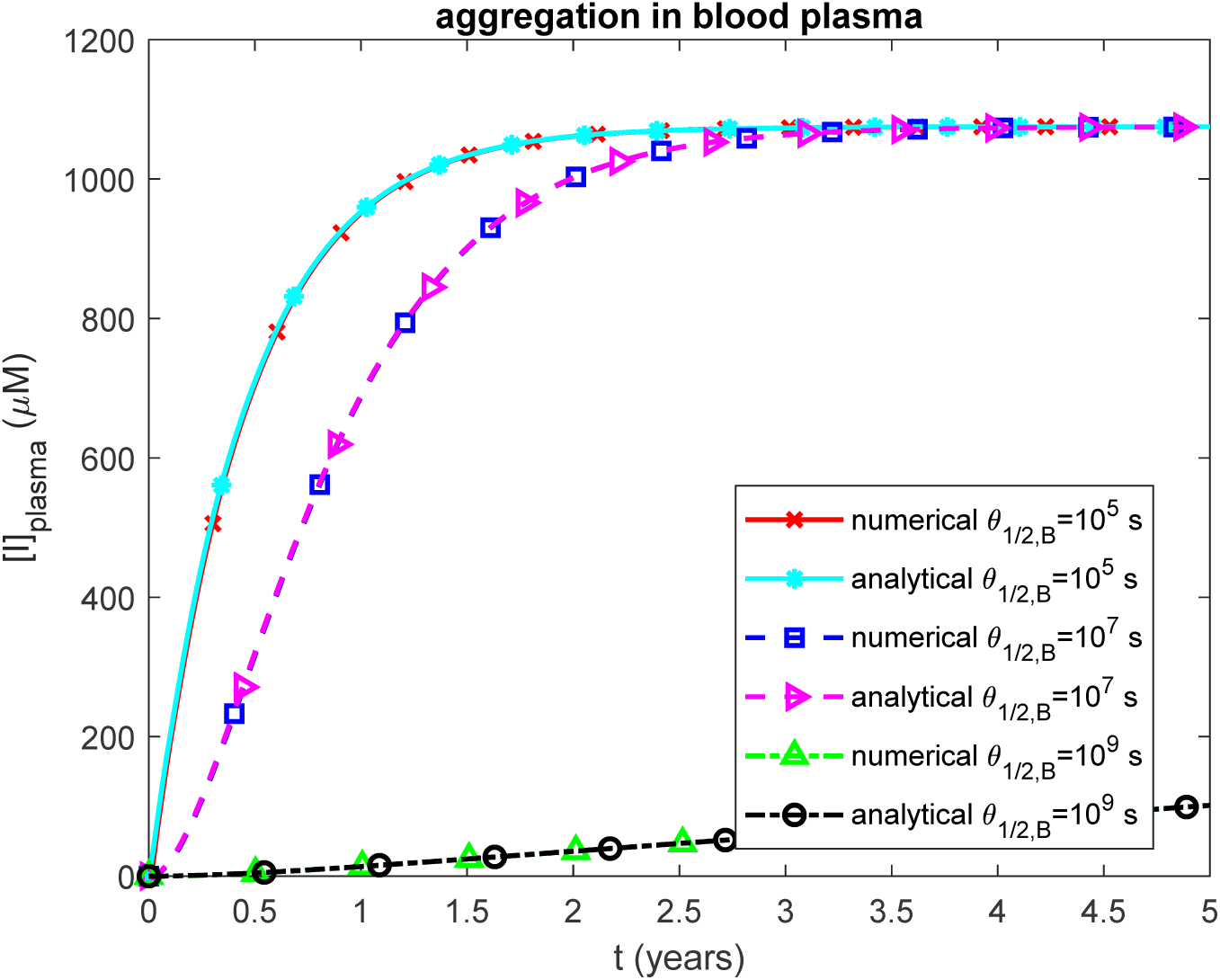
Molar concentration of TTR oligomers deposited into amyloid protofibrils circulating in the blood plasma vs time for various values of *θ*_1/2,*B*_. Scenario when TTR monomers misfold and assemble into protofibrillar formations within the bloodstream before depositing in the heart.

**Fig. S11.**
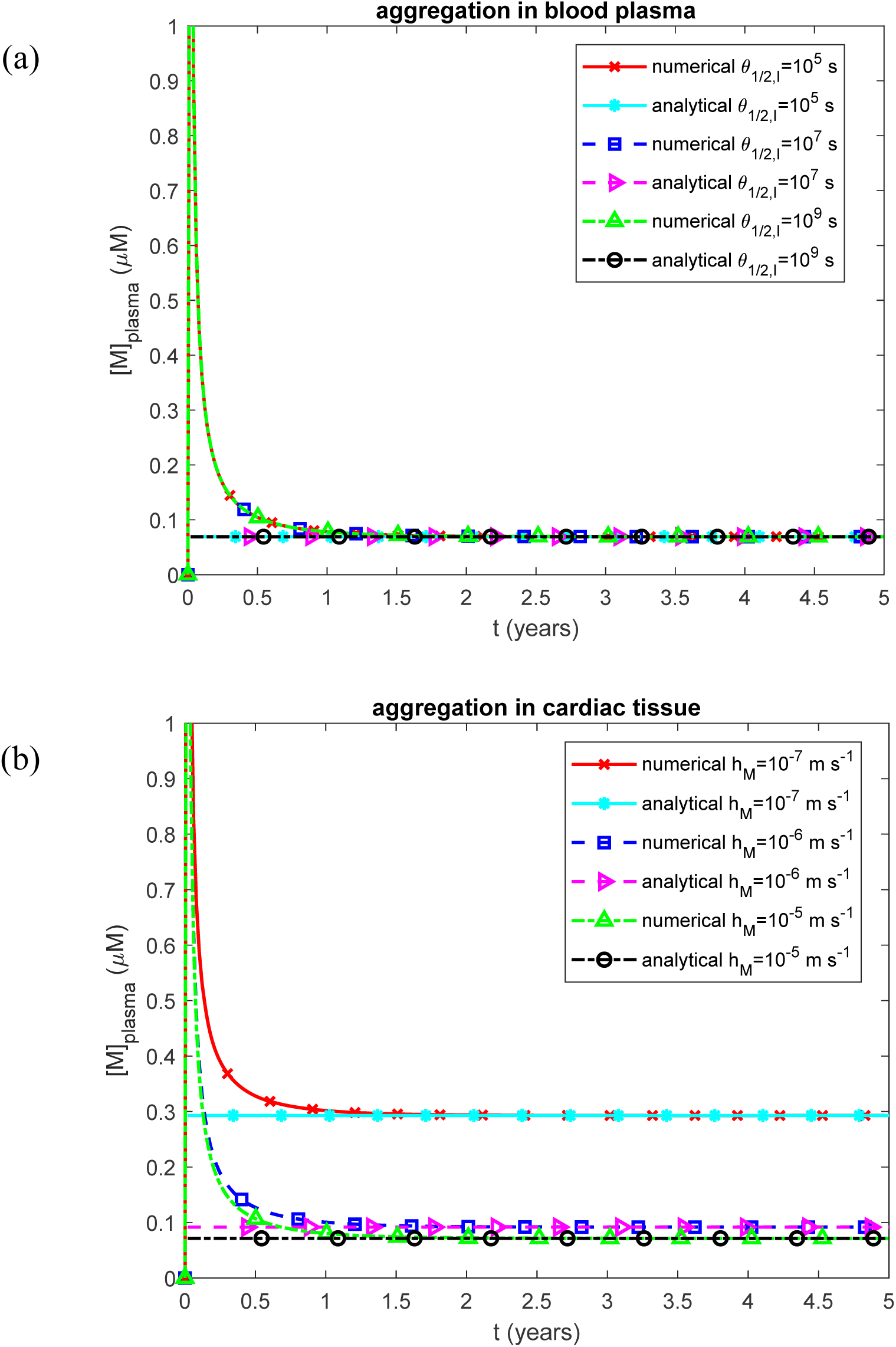
Molar concentration of TTR monomers in the blood plasma vs time. (a) Scenario when TTR monomers misfold and assemble into protofibrillar formations within the bloodstream before depositing in the heart for various values of *θ*_1/2,*I*_. (b) Scenario when TTR monomers remain in circulation until they directly attach to sites within the heart, where they form amyloid deposits *in situ* for various values of *h_M_*.

**Fig. S12.**
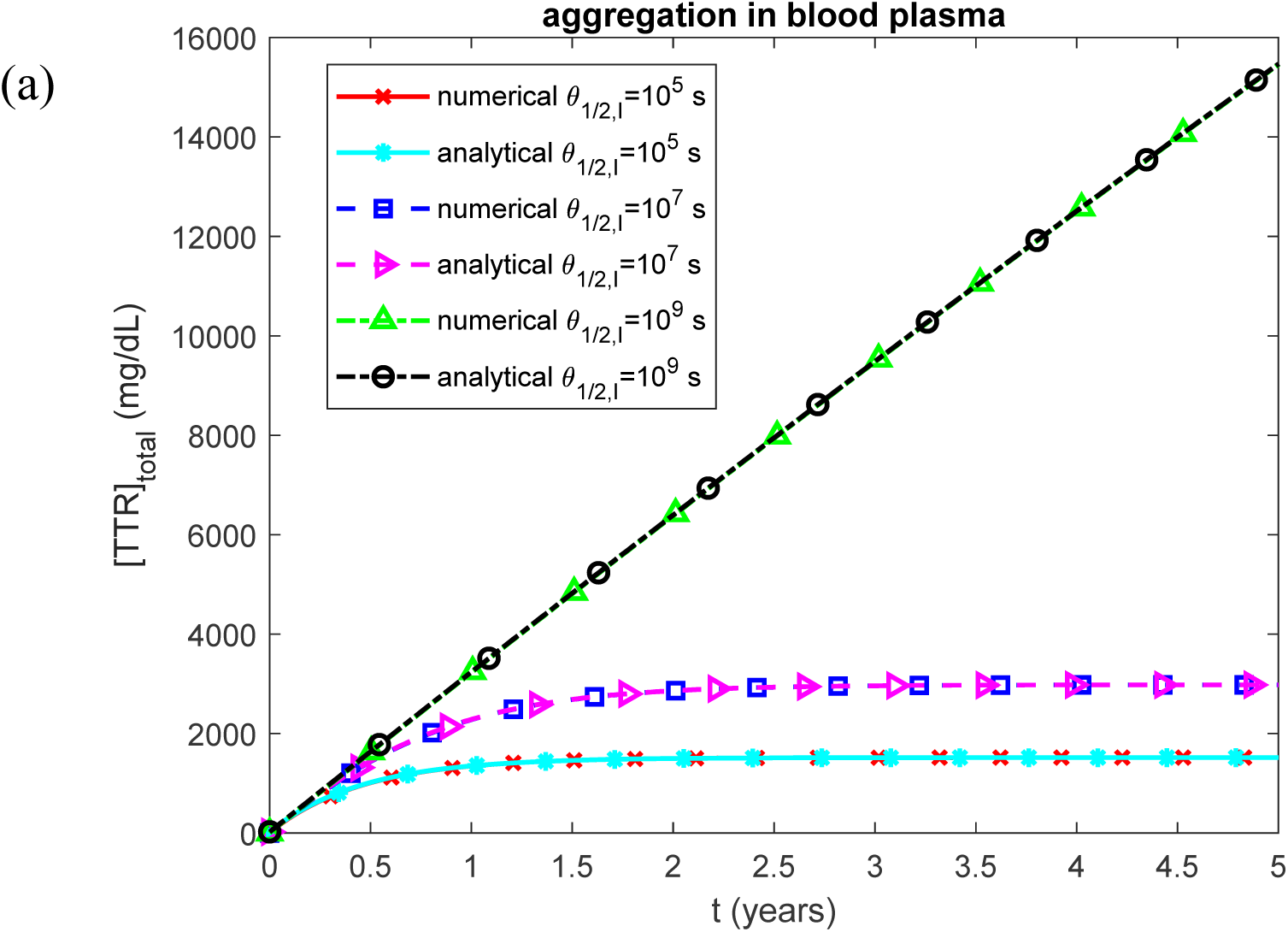
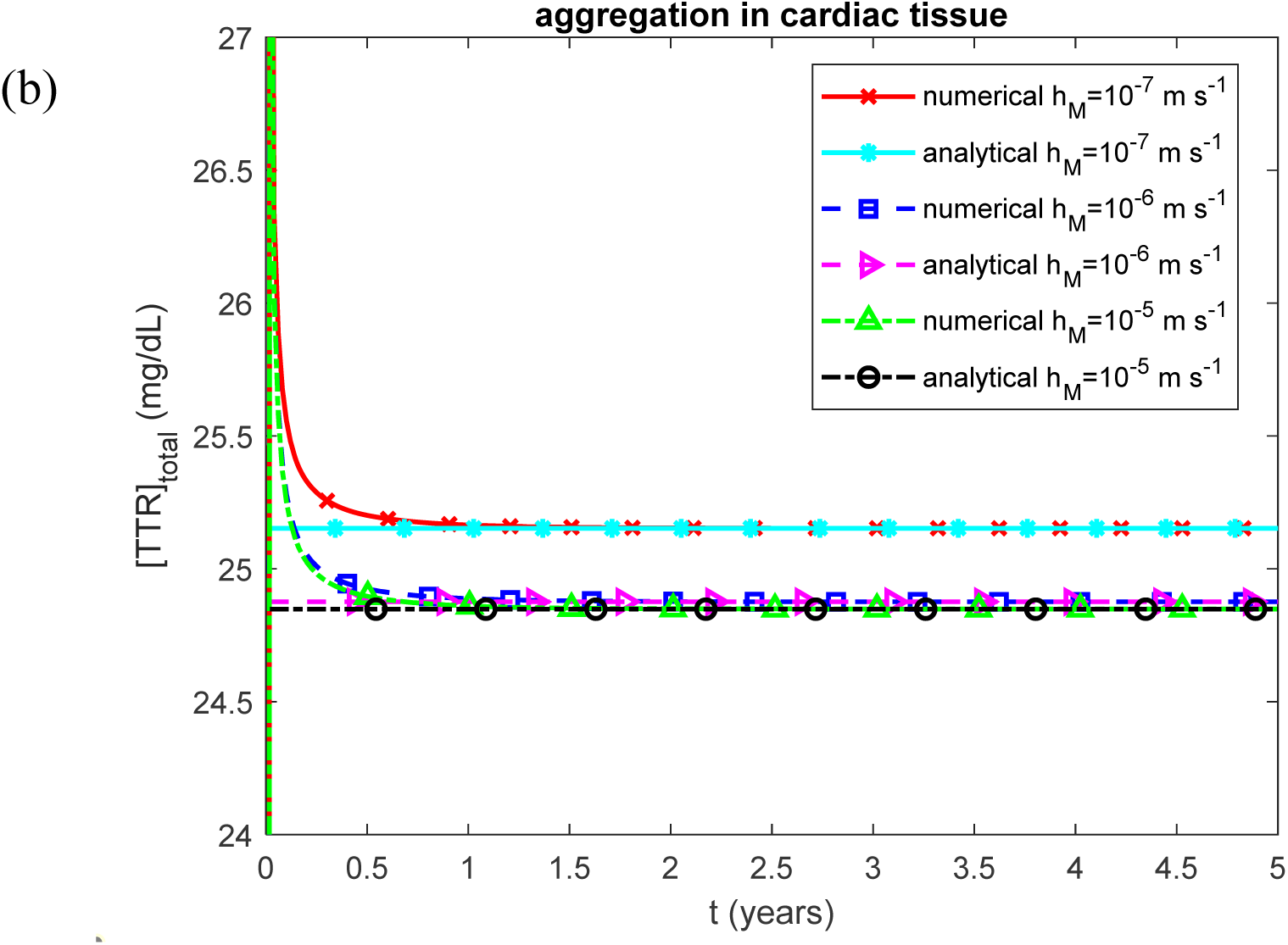
Total concentration of TTR (in any state) in the blood plasma vs time. (a) Scenario when TTR monomers misfold and assemble into protofibrillar formations within the bloodstream before depositing in the heart for various values of *θ*_1/2,*I*_. (b) Scenario when TTR monomers remain in circulation until they directly attach to sites within the heart, where they form amyloid deposits *in situ* for various values of *h_M_*.

**Fig. S13.**
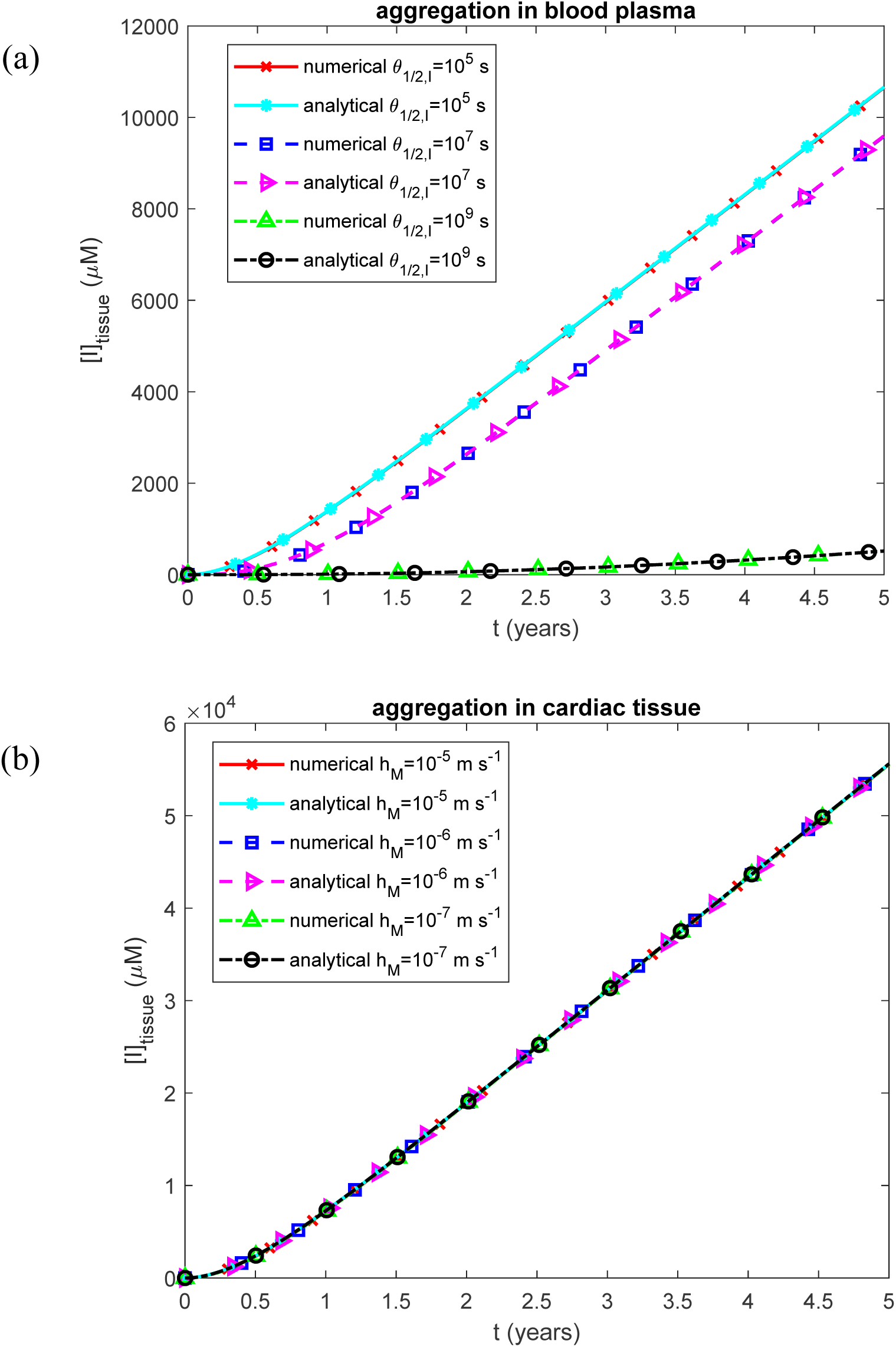
Molar concentration of TTR oligomers deposited into amyloid fibrils that infiltrated the cardiac tissue vs time. (a) Scenario when TTR monomers misfold and assemble into protofibrillar formations within the bloodstream before depositing in the heart for various values of *θ*_1/2,*I*_. (b) Scenario when TTR monomers remain in circulation until they directly attach to sites within the heart, where they form amyloid deposits *in situ* for various values of *h_M_*.

**Fig. S14.**
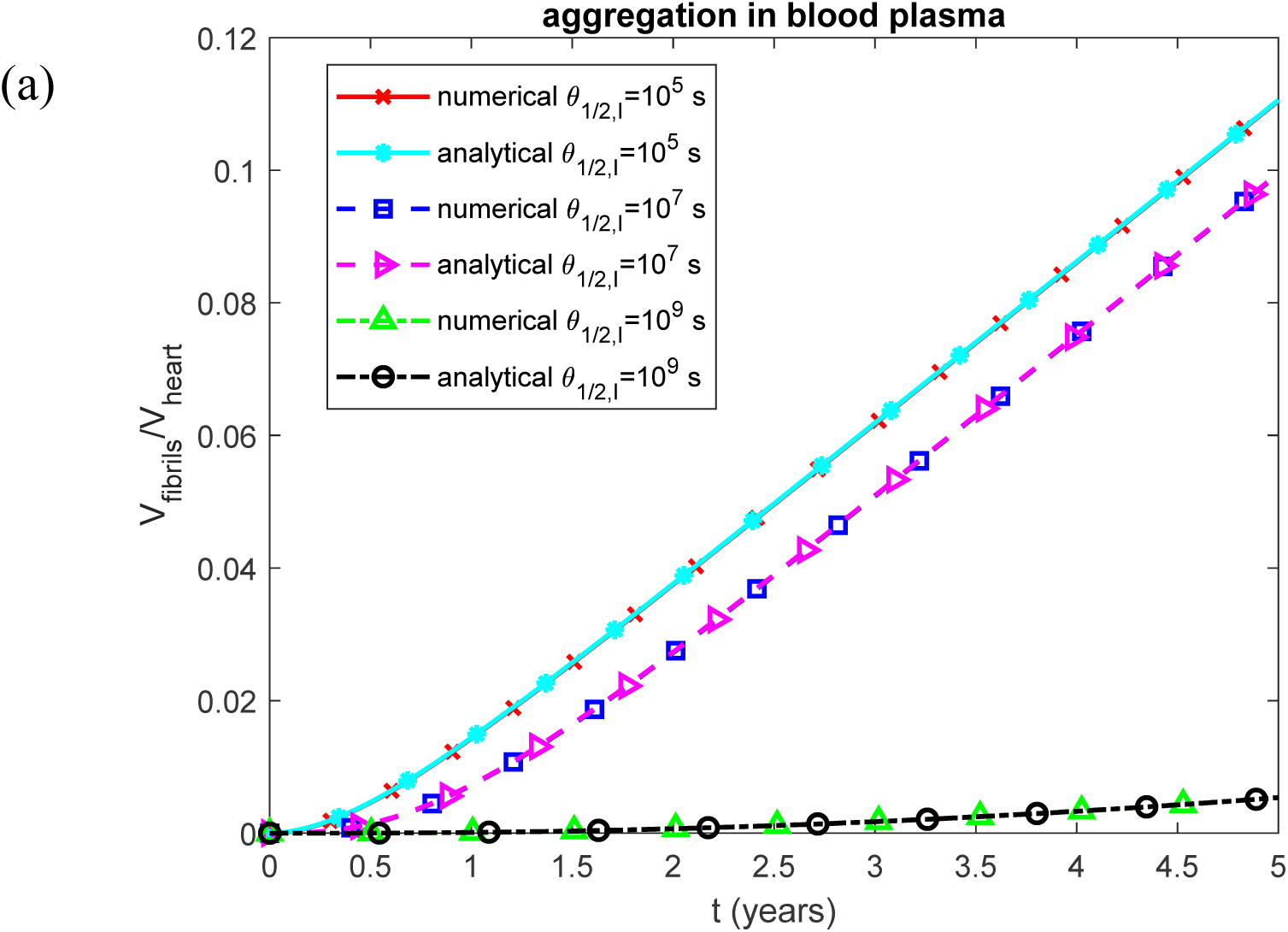
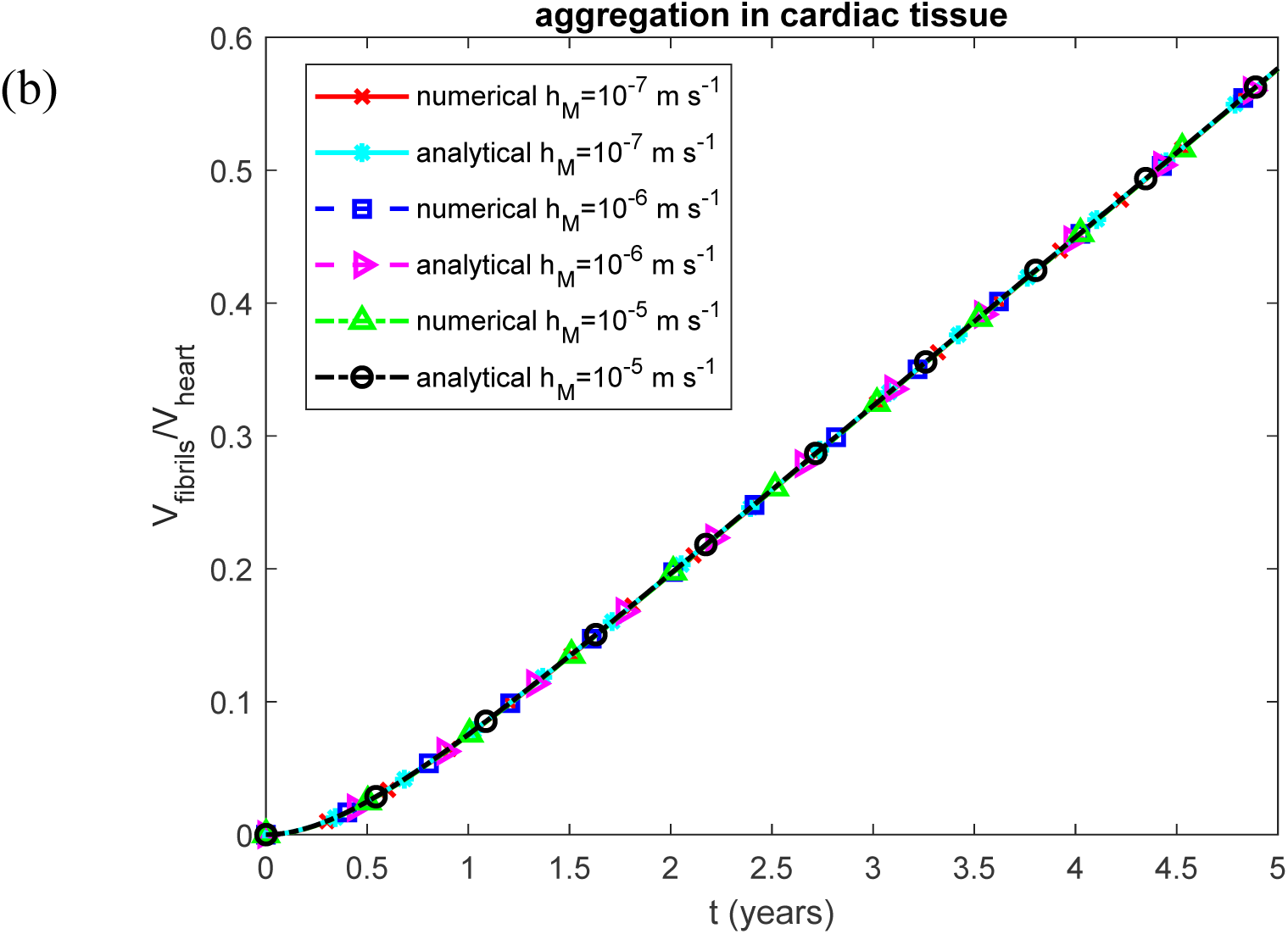
Volumetric ratio of deposited TTR fibrils to baseline myocardial volume vs time. (a) Scenario when TTR monomers misfold and assemble into protofibrillar formations within the bloodstream before depositing in the heart for various values of *θ*_1/2,*I*_. (b) Scenario when TTR monomers remain in circulation until they directly attach to sites within the heart, where they form amyloid deposits *in situ* for various values of *h_M_*.

**Fig. S15.**
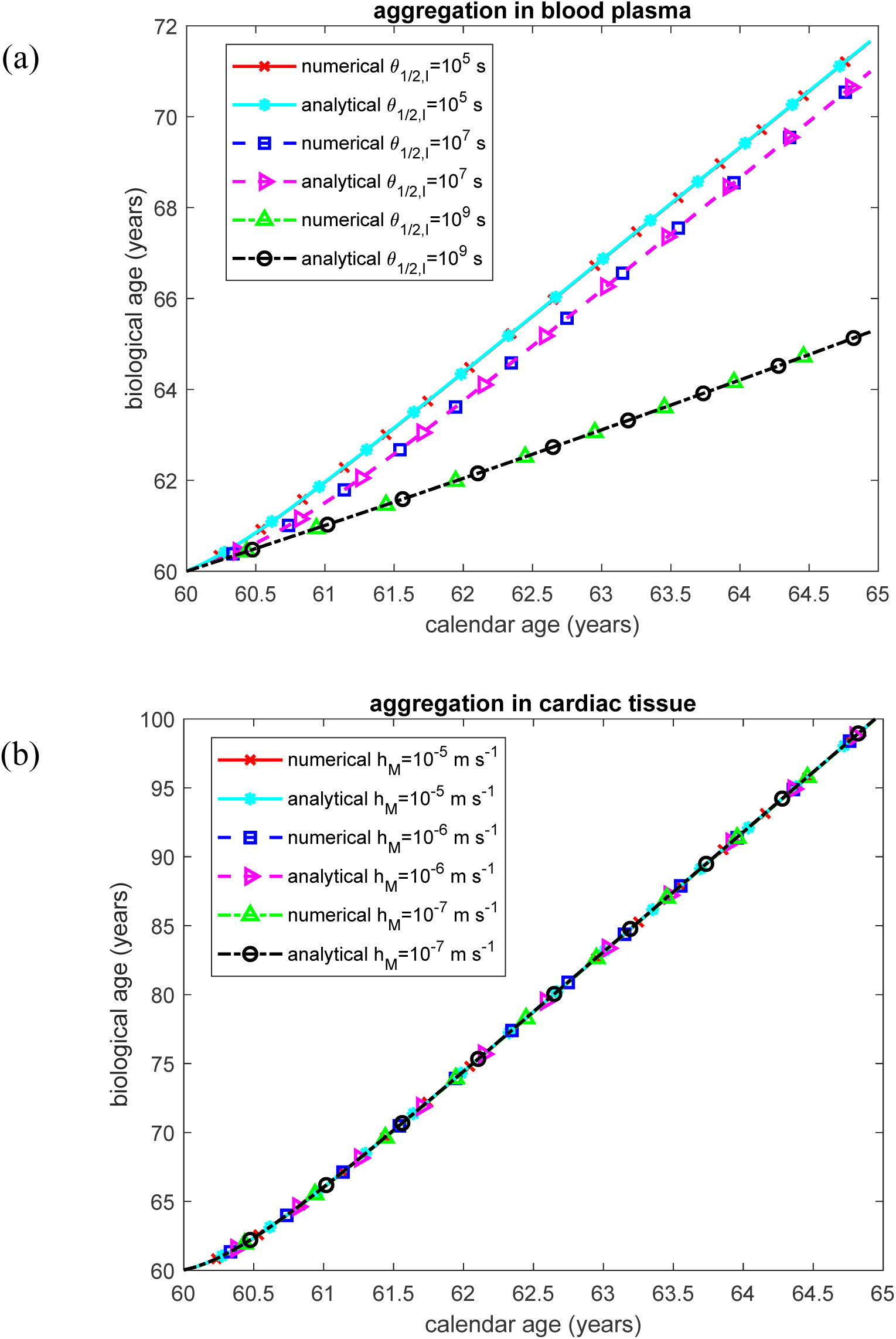
Biological age vs calendar age. (a) Scenario when TTR monomers misfold and assemble into protofibrillar formations within the bloodstream before depositing in the heart for various values of *θ*_1/2,*I*_. (b) Scenario when TTR monomers remain in circulation until they directly attach to sites within the heart, where they form amyloid deposits *in situ* for various values of *h_M_*.

**Fig. S16.**
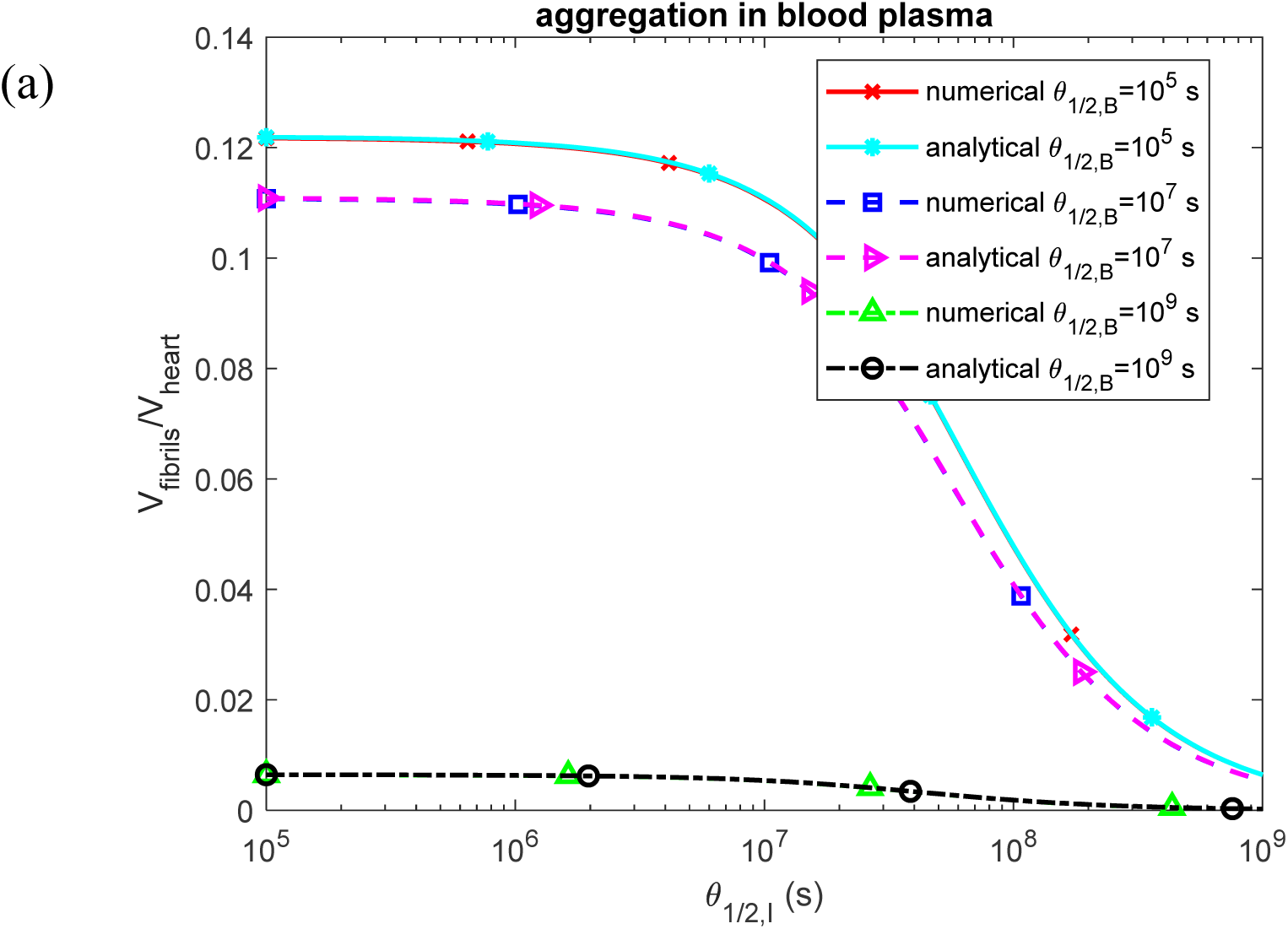
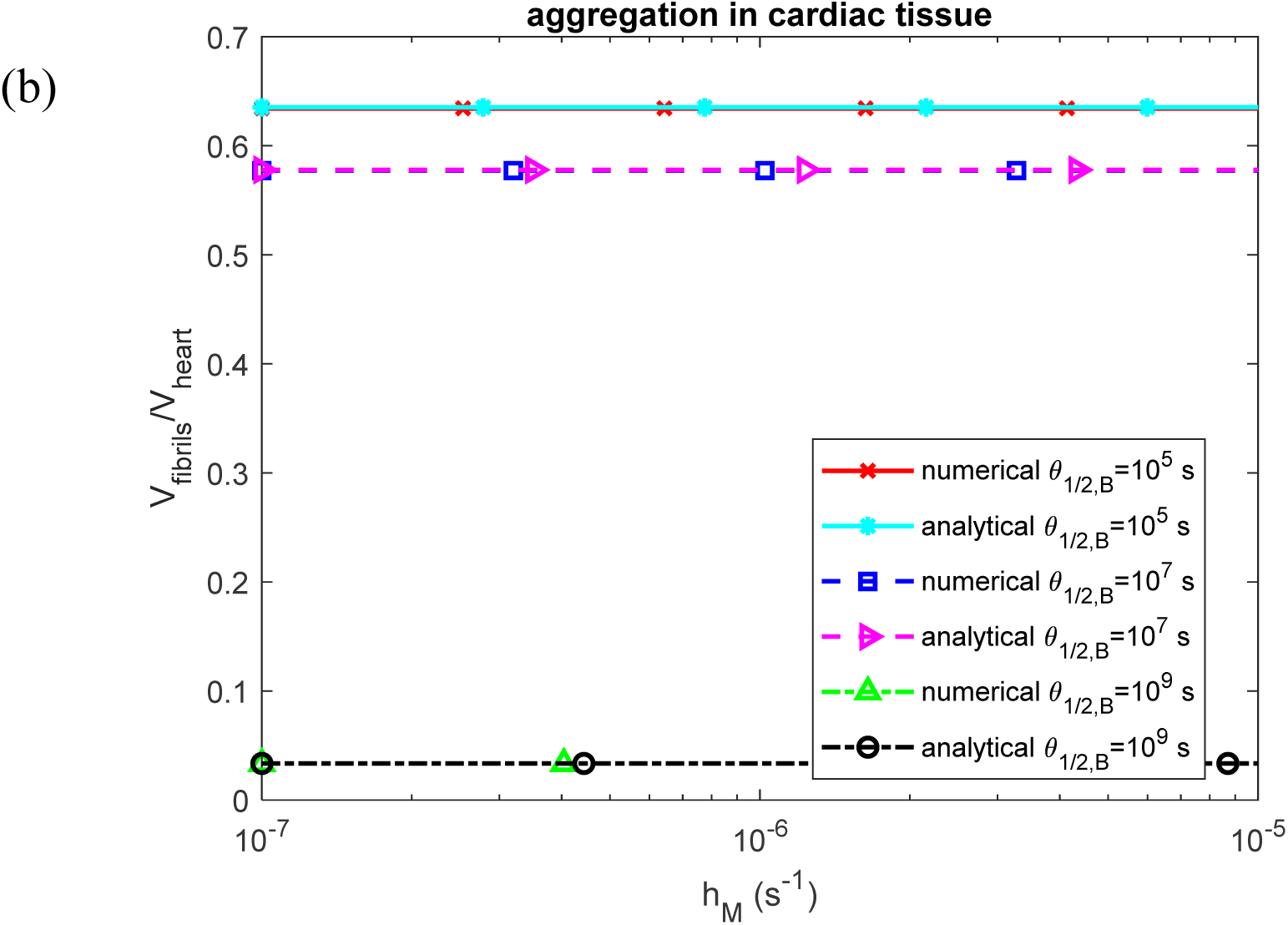
Volumetric ratio of deposited TTR fibrils to baseline myocardial volume. (a) As a function of the half-deposition time for TTR-derived fibrils circulating in the blood plasma to deposit into cardiac tissue. This corresponds to the scenario in which TTR monomers misfold and assemble into protofibrils within the bloodstream before depositing in the heart, shown for various values of *θ*_1/2,*B*_. (b) As a function of the mass transfer coefficient characterizing the rate at which circulating TTR monomers deposit into the endocardium. This represents the scenario in which TTR monomers remain in circulation until they bind directly to cardiac sites and form amyloid deposits *in situ*, shown for various values of *θ*_1/ 2,*B*_.

**Fig. S17.**
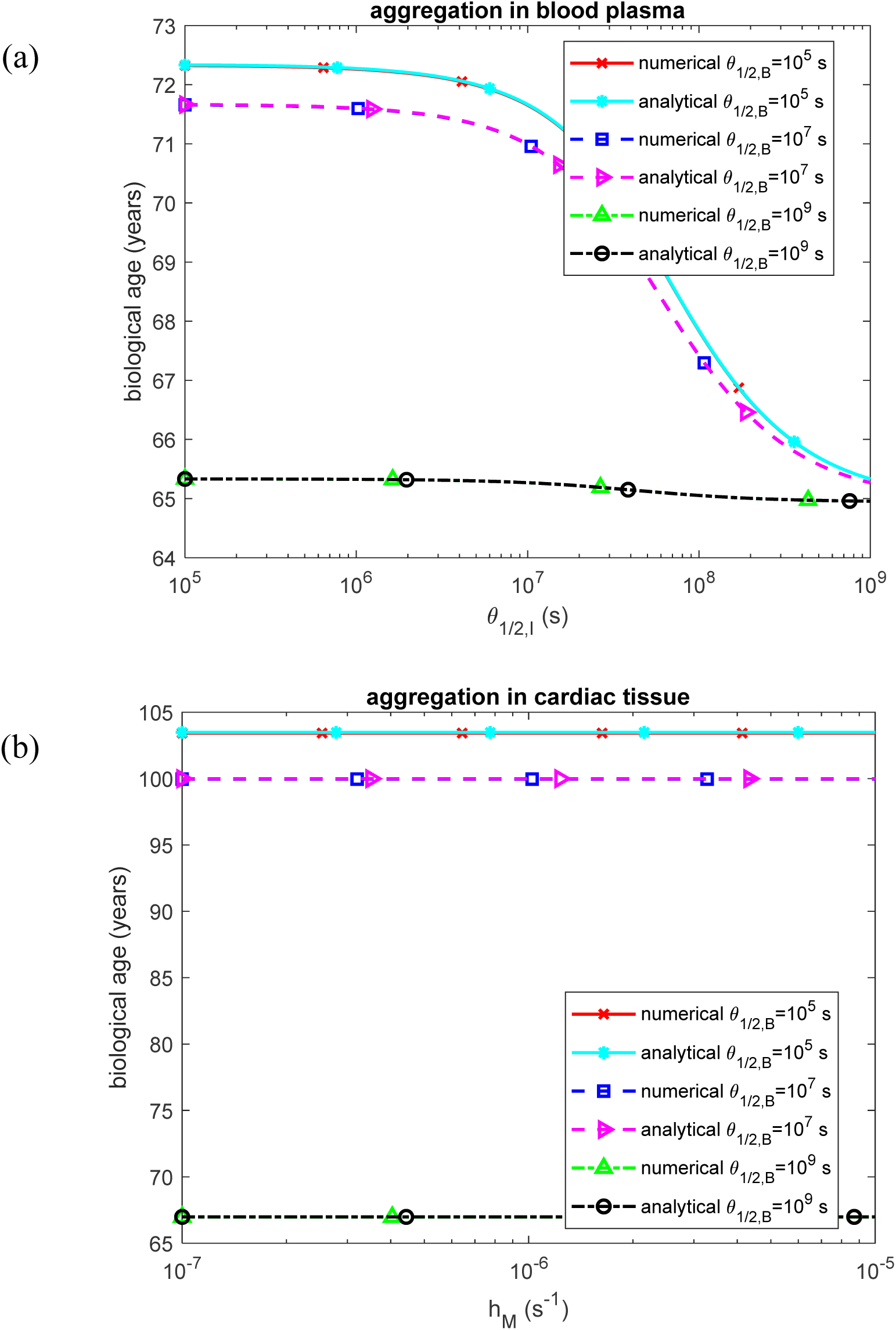
Biological age. (a) As a function of the half-deposition time for TTR-derived fibrils circulating in the blood plasma to deposit into cardiac tissue. This corresponds to the scenario in which TTR monomers misfold and assemble into protofibrils within the bloodstream before depositing in the heart, shown for various values of *θ*_1/2,*B*_. (b) As a function of the mass transfer coefficient characterizing the rate at which circulating TTR monomers deposit into the endocardium. This represents the scenario in which TTR monomers remain in circulation until they bind directly to cardiac sites and form amyloid deposits *in situ*, shown for various values of *θ*_1/2,*B*_.

**Fig. S18.**
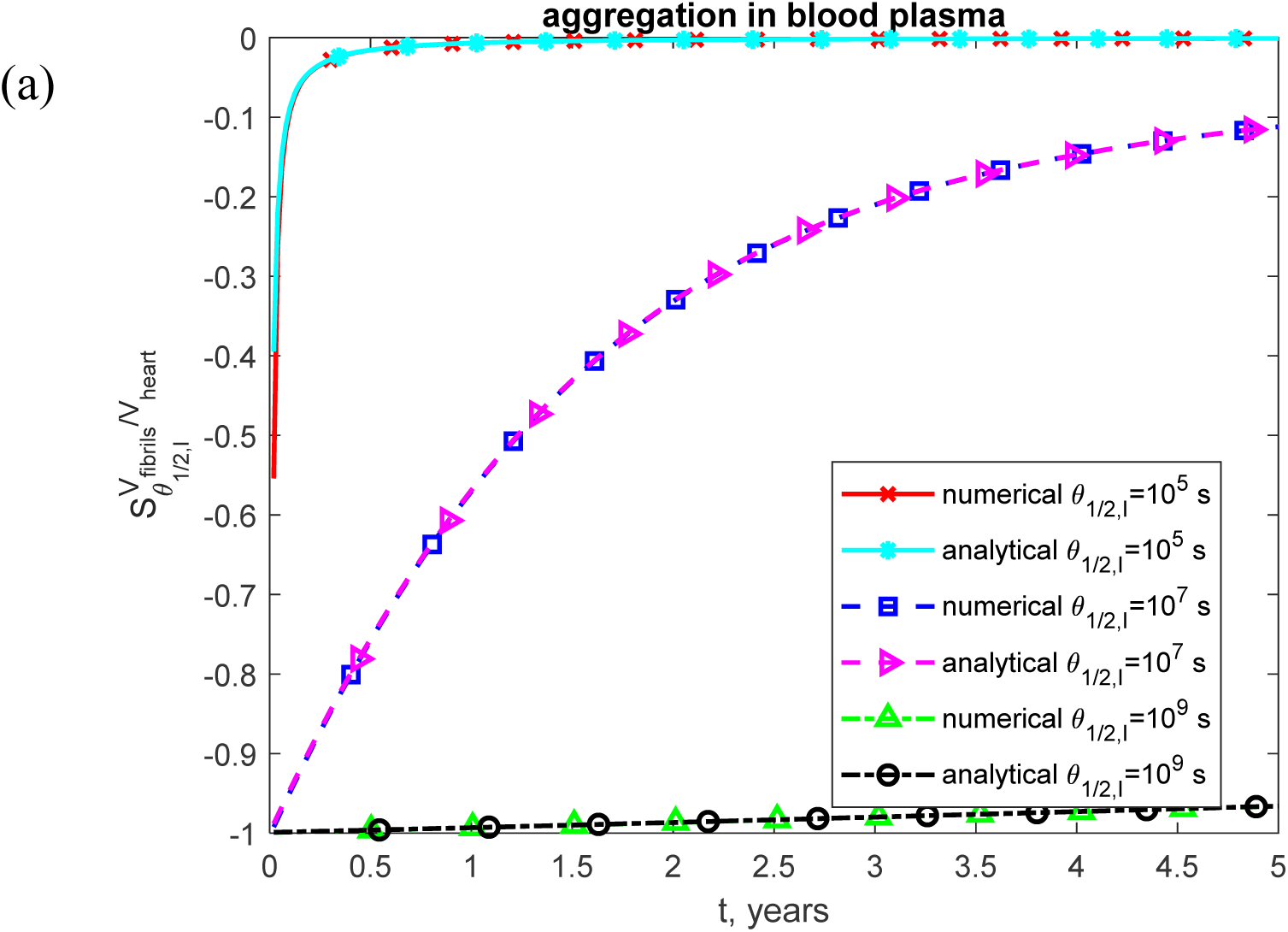
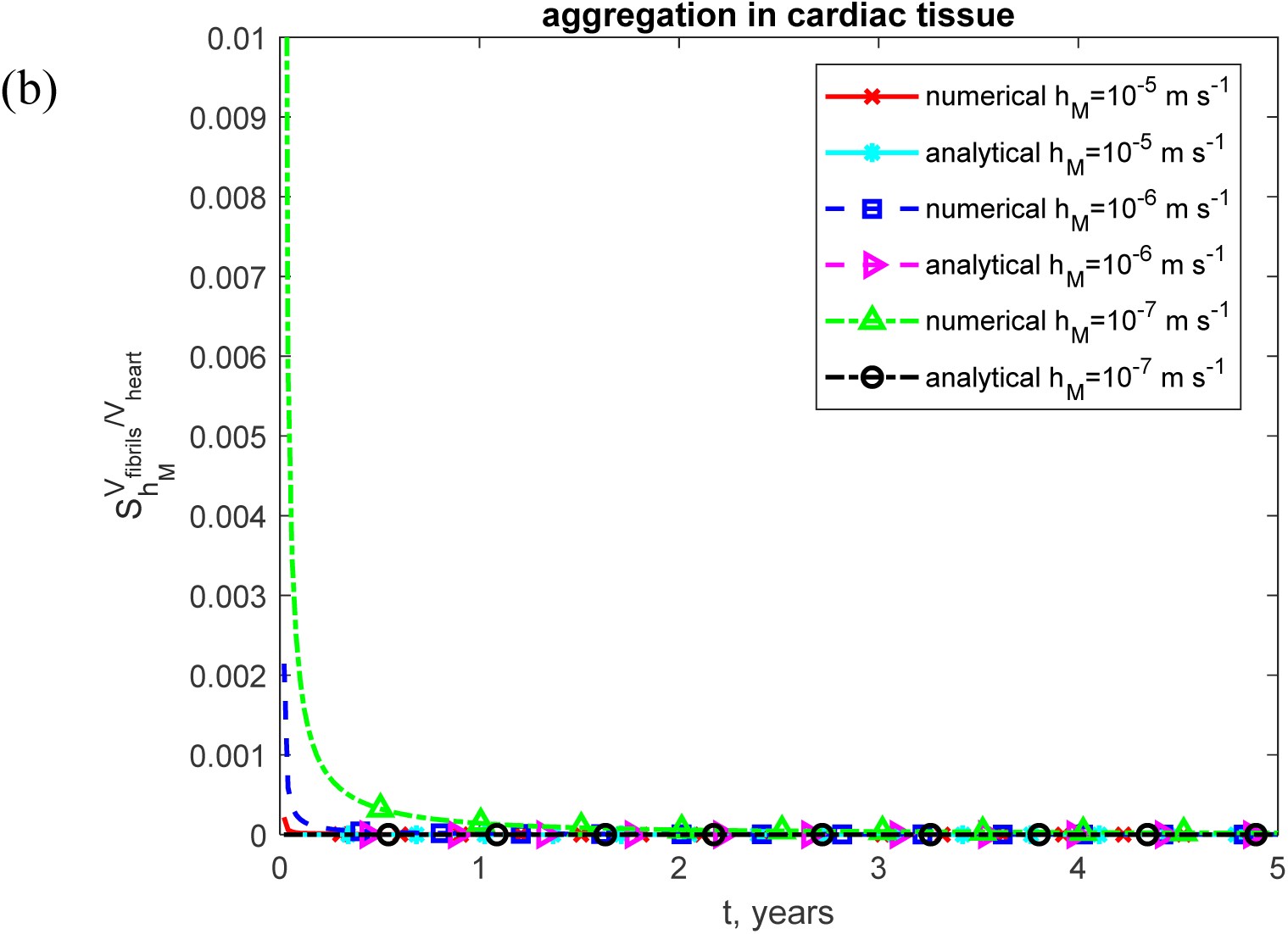
(a) Dimensionless sensitivity of the fraction of heart volume occupied by TTR amyloid fibrils to the half-deposition time for TTR-derived fibrils circulating in the blood plasma to deposit into cardiac tissue, *θ*_1/2,*I*_, shown as a function of time. This corresponds to the scenario in which TTR monomers misfold and assemble into protofibrillar structures within the bloodstream before depositing in the heart. (b) Dimensionless sensitivity of the fraction of heart volume occupied by TTR amyloid fibrils to the mass transfer coefficient characterizing the rate of deposition of TTR monomers into endocardium, *h_M_*, also shown as a function of time. This corresponds to the scenario in which TTR monomers remain in circulation until they directly bind to sites within the heart, forming amyloid deposits *in situ*.

**Fig. S19.**
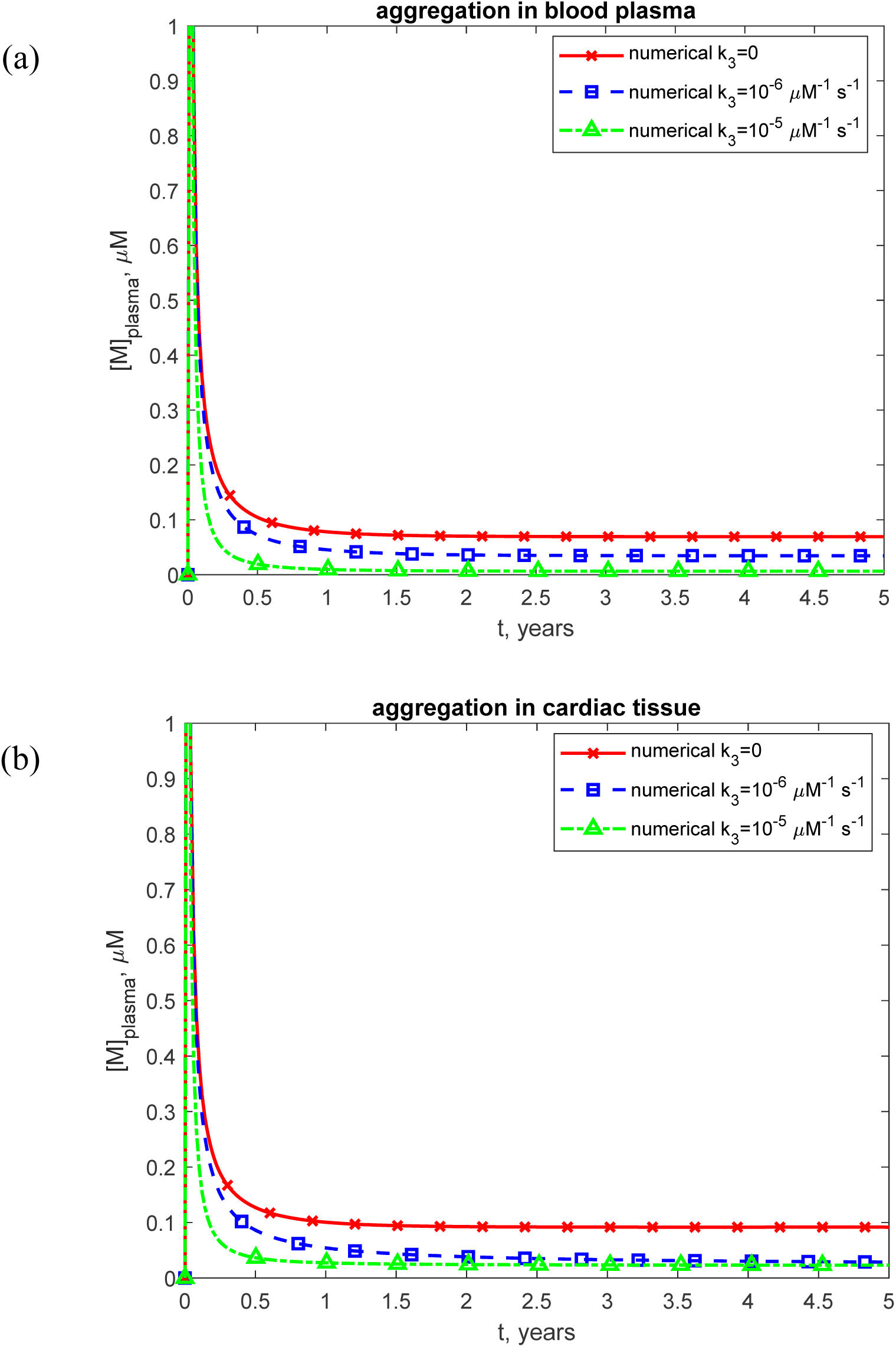
Molar concentration of TTR monomers in the blood plasma vs time for various values of *k*_3_. (a) Scenario when TTR monomers misfold and assemble into protofibrillar formations within the bloodstream before depositing in the heart. (b) Scenario when TTR monomers remain in circulation until they directly attach to sites within the heart, where they form amyloid deposits *in situ*.

**Fig. S20.**
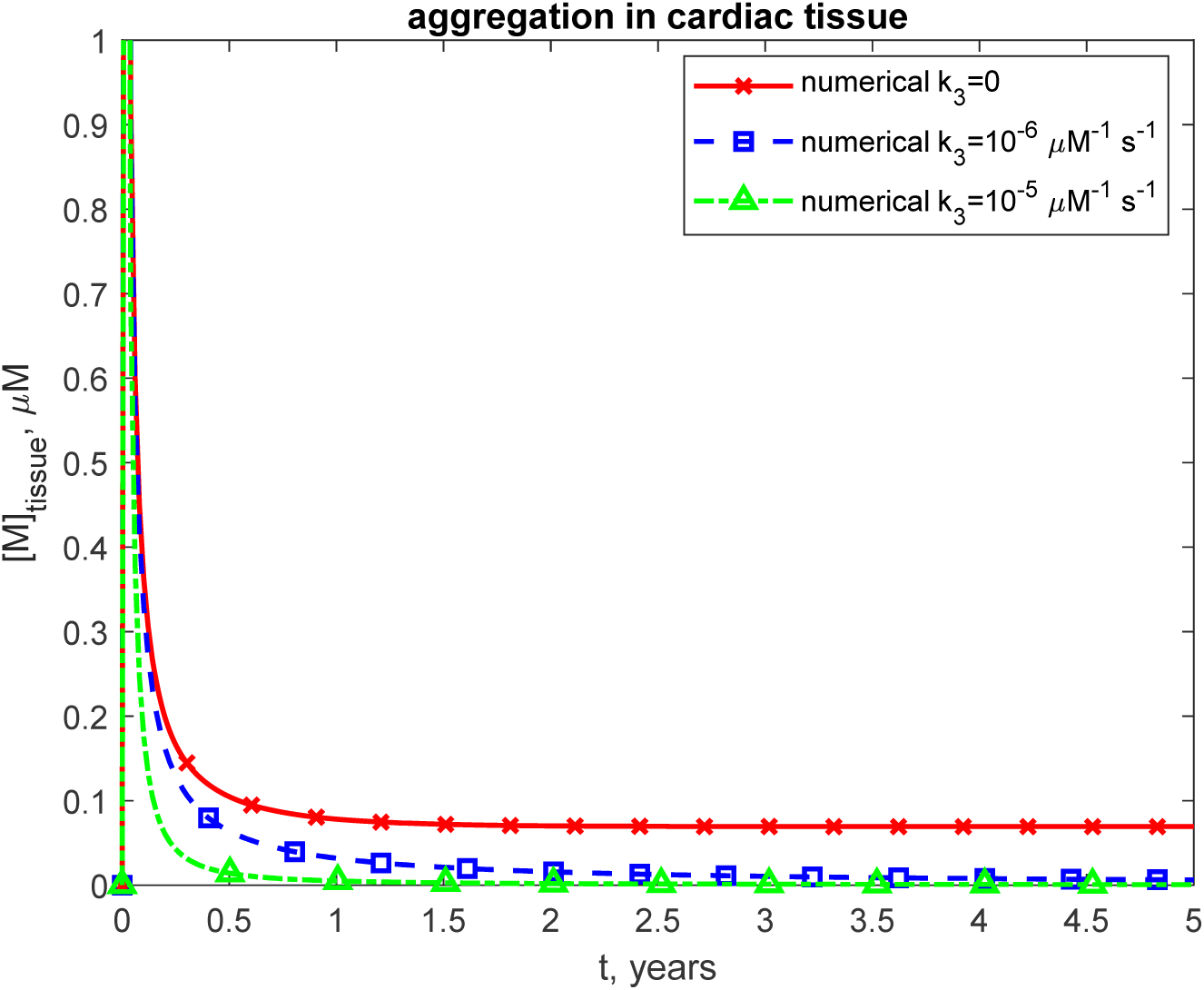
Molar concentration of TTR monomers in the cardiac tissue vs time for various values of *k*_3_. Scenario when TTR monomers remain in circulation until they directly attach to sites within the heart, where they form amyloid deposits *in situ*.

**Fig. S21.**
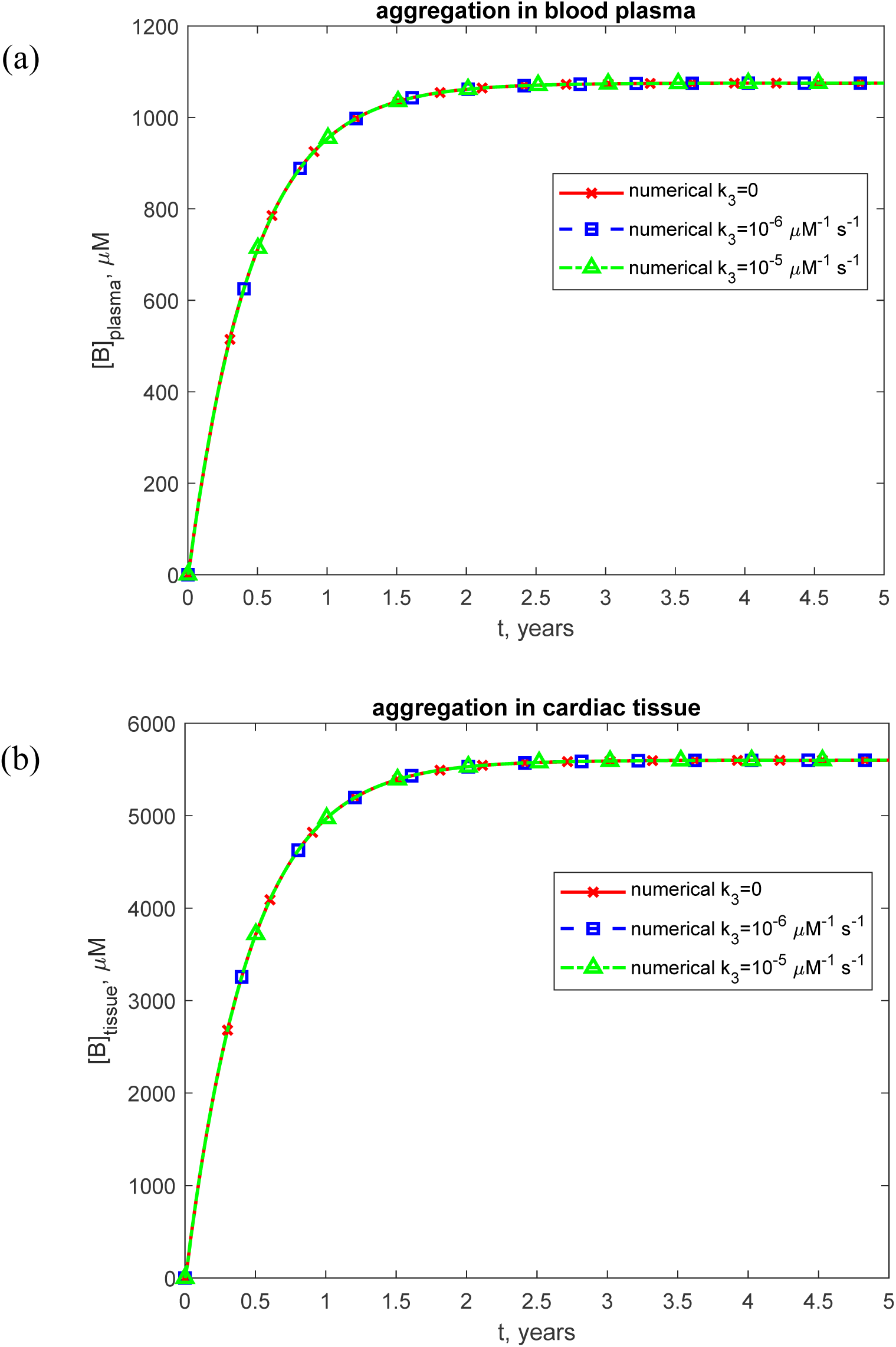
(a) Molar concentration of free TTR oligomers in the blood plasma vs time for various values of *k*_3_. Scenario when TTR accumulates in blood plasma. (b) Molar concentration of free TTR oligomers in cardiac tissue vs time for various values of *k*_3_. Scenario when TTR accumulates in cardiac tissue.

**Fig. S22.**
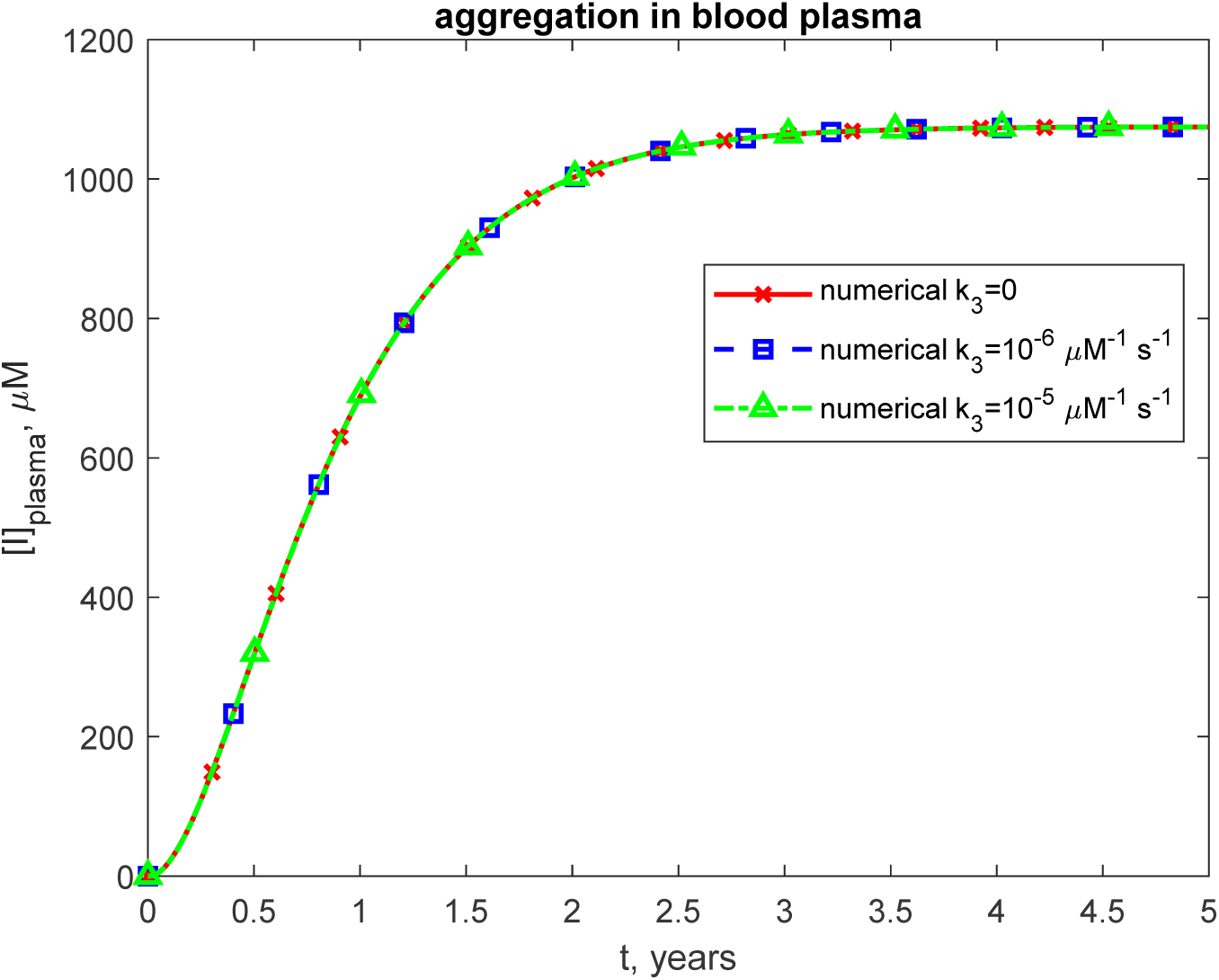
Molar concentration of TTR oligomers deposited into amyloid protofibrils circulating in the blood plasma vs time for various values of *k*_3_. Scenario when TTR monomers misfold and assemble into protofibrillar formations within the bloodstream before depositing in the heart.

**Fig. S23.**
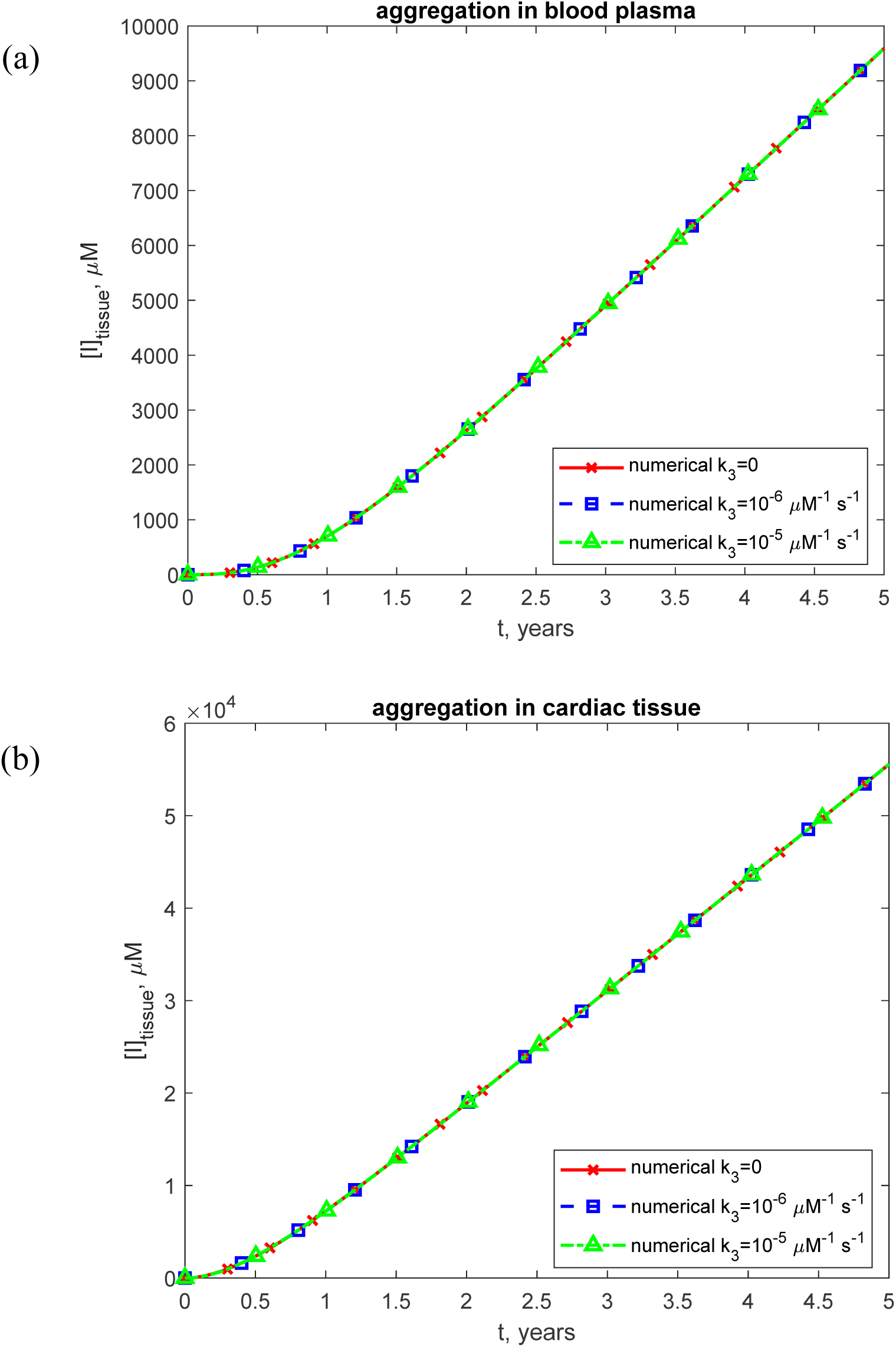
Molar concentration of TTR oligomers deposited into amyloid fibrils that infiltrated the cardiac tissue vs time for various values of *k*_3_. (a) Scenario when TTR monomers misfold and assemble into protofibrillar formations within the bloodstream before depositing in the heart. (b) Scenario when TTR monomers remain in circulation until they directly attach to sites within the heart, where they form amyloid deposits *in situ*.

**Fig. S24.**
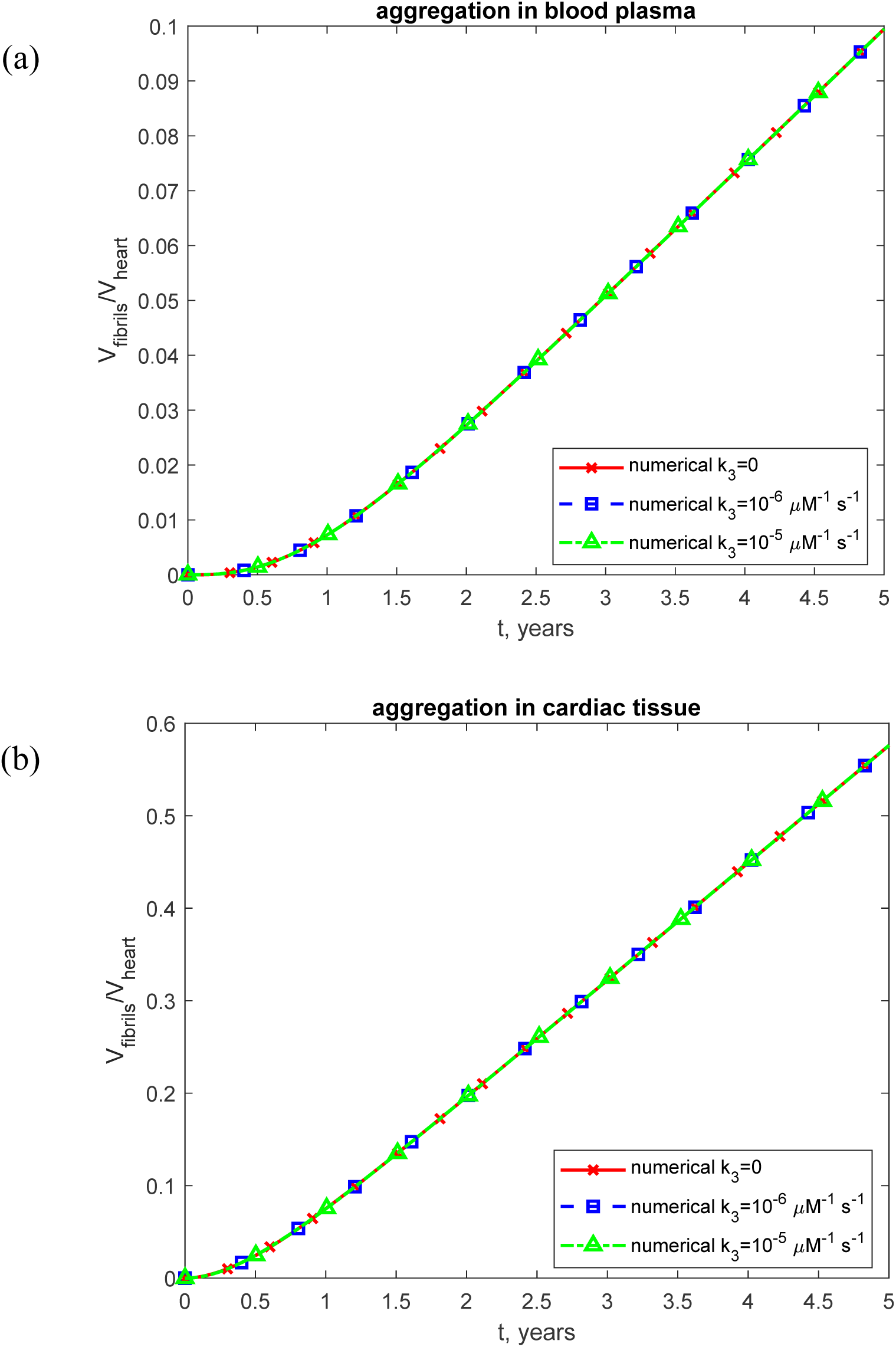
Volumetric ratio of deposited TTR fibrils to baseline myocardial volume for various values of *k*_3_. (a) As a function of the half-deposition time for TTR-derived fibrils circulating in the blood plasma to deposit into cardiac tissue. This corresponds to the scenario in which TTR monomers misfold and assemble into protofibrils within the bloodstream before depositing in the heart. (b) As a function of the mass transfer coefficient characterizing the rate at which circulating TTR monomers deposit into the endocardium. This represents the scenario in which TTR monomers remain in circulation until they bind directly to cardiac sites and form amyloid deposits *in situ*.

**Fig. S25.**
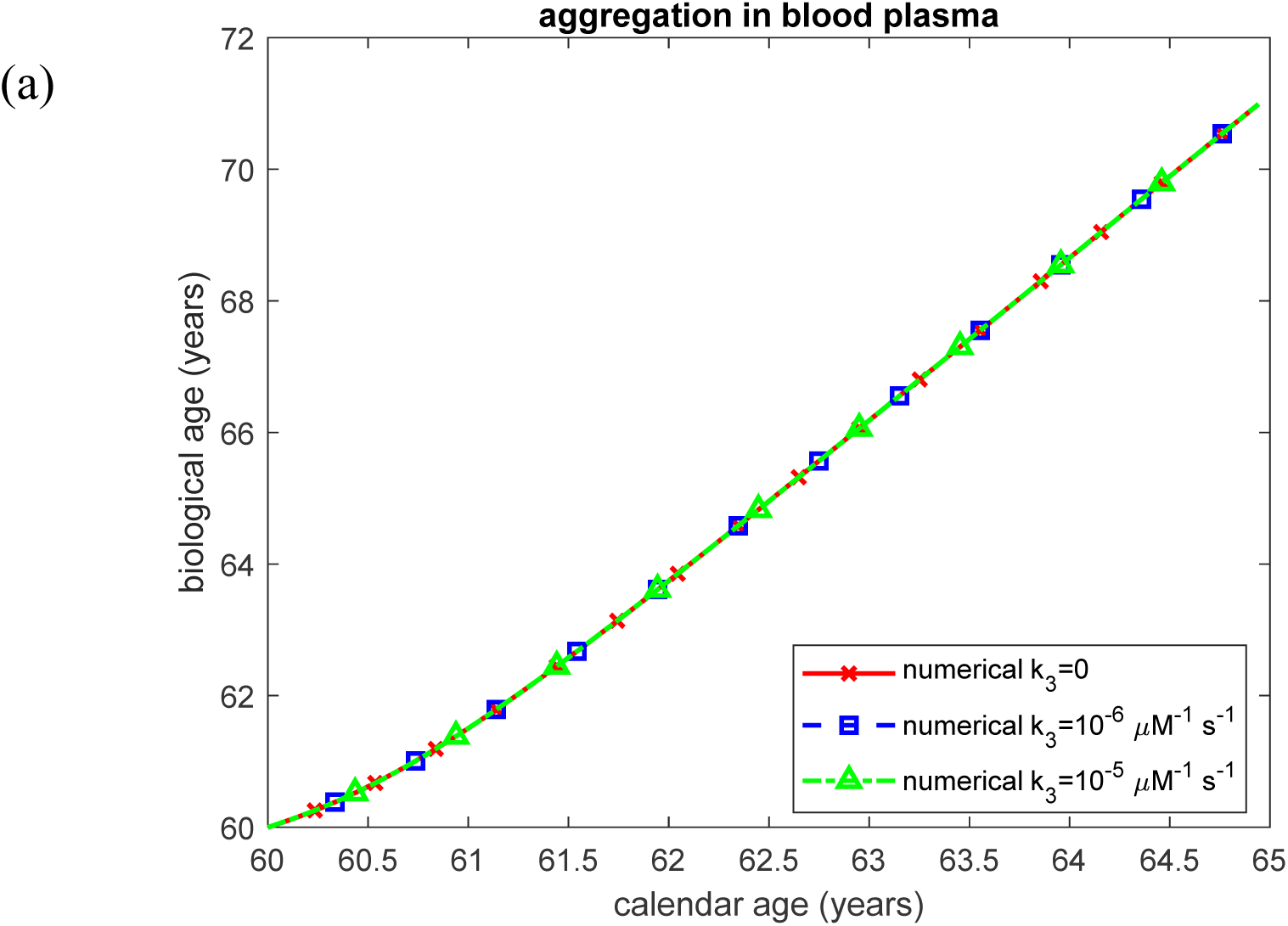
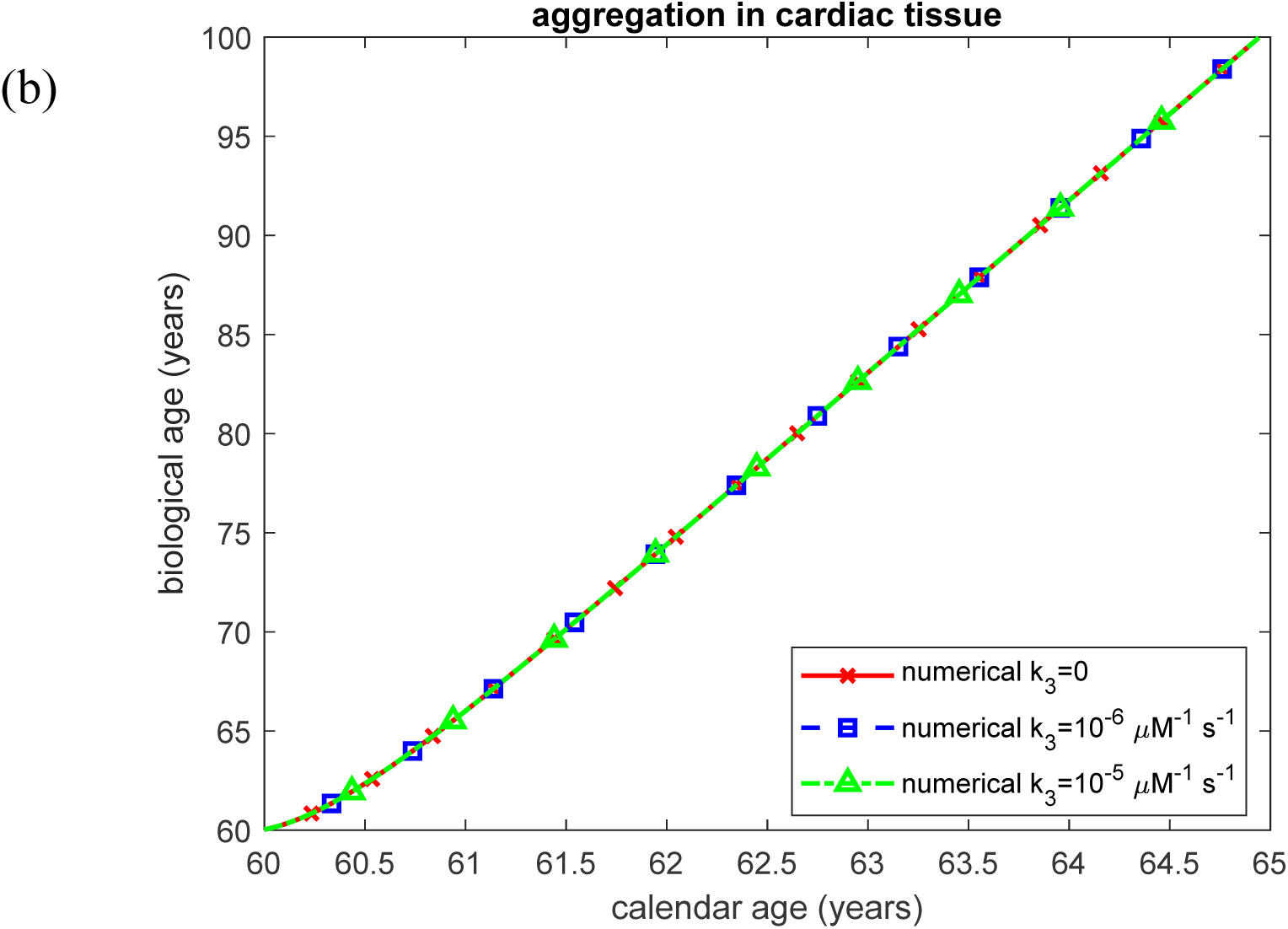
Biological age vs calendar age for various values of *k*_3_. (a) Scenario when TTR monomers misfold and assemble into protofibrillar formations within the bloodstream before depositing in the heart. (b) Scenario when TTR monomers remain in circulation until they directly attach to sites within the heart, where they form amyloid deposits *in situ*.

